# Integrated phylogenomics and fossil data illuminate the evolution of beetles

**DOI:** 10.1101/2021.09.22.461358

**Authors:** Chenyang Cai, Erik Tihelka, Mattia Giacomelli, John F. Lawrence, Adam Ślipiński, Robin Kundrata, Shûhei Yamamoto, Margaret K. Thayer, Alfred F. Newton, Richard A. B. Leschen, Matthew L. Gimmel, Liang Lü, Michael S. Engel, Diying Huang, Davide Pisani, Philip C.J. Donoghue

## Abstract

With over 380,000 described species and possibly several million more yet unnamed, beetles represent the most biodiverse animal order. Recent phylogenomic studies have arrived at considerably incongruent topologies and widely varying estimates of divergence dates for major beetle clades. Here we use a dataset of 68 single-copy nuclear protein coding genes sampling 129 out of the 194 recognized extant families as well as the first comprehensive set of fully-justified fossil calibrations to recover a refined timescale of beetle evolution. Using phylogenetic methods that counter the effects of compositional and rate heterogeneity we recover a topology congruent with morphological studies, which we use, combined with other recent phylogenomic studies, to propose several formal changes in the classification of Coleoptera: Scirtiformia and Scirtoidea *sensu nov*., Clambiformia *ser. nov.* and Clamboidea *sensu nov.*, Rhinorhipiformia *ser. nov*., Byrrhoidea *sensu nov.*, Dryopoidea *stat. res.*, Nosodendriformia *ser. nov.*, and Staphyliniformia *sensu nov*., alongside changes below the superfamily level. The heterogeneous former superfamily Cucujoidea is divided into three monophyletic groups: Erotyloidea *stat. nov*., Nitiduloidea *stat. nov*., and Cucujoidea *sensu nov.* Our divergence time analysis recovered an evolutionary timescale congruent with the fossil record: a late Carboniferous origin of Coleoptera, a late Paleozoic origin of all modern beetle suborders, and a Triassic–Jurassic origin of most extant families. While fundamental divergences within beetle phylogeny did not coincide with the hypothesis of a Cretaceous Terrestrial Revolution, many polyphagan superfamilies exhibited increases in richness with Cretaceous flowering plants.

## Introduction

Beetles (Coleoptera) are a textbook example of a hyperdiverse clade, known from more than 380,000 living species and upwards of 1.5 million awaiting description^1,2^, that display extraordinary morphological, taxonomic, and ecological diversity^3^. Constituting nearly a quarter of extant animal diversity on our shared planet, beetles play indispensable roles in nearly all terrestrial and freshwater ecosystems. The ecological dominance of beetles is reflected by their fossil record. The earliest unequivocal stem-beetles are Early Permian^4,5^, while crown beetles belonging to extant suborders (Adephaga, Archostemata, Myxophaga and Polyphaga) first occur in the late Permian^6,7^, and most extant families are first encountered in the fossil record in the Jurassic to Cretaceous^8–10^. A multitude of hypotheses have been proposed to explain beetle megadiversity, focusing principally on the importance of key anatomical innovations, co-diversification with other clades such as angiosperms during the Cretaceous Terrestrial Revolution^11^, and mass extinction events^9,12–15^. Tests of these hypotheses of the causes and consequences of beetle diversification require a robust time-calibrated phylogeny. However, considering the long evolutionary history, exceptional species richness, and unparalleled morphological disparity as well as apparent morphological convergence of beetles, resolving the phylogeny and timescale of Coleoptera evolution has proven challenging.

The lack of a consensus on higher-level relationships within Coleoptera has compromised attempts to derive an evolutionary timescale. To date, the majority of molecular phylogenetic studies of beetles with the most comprehensive taxon sampling^9,16–18^ have sampled eight or fewer genes, with a matrix length of less than 10,000 nucleotides^19^. While this limited gene sampling has helped shed light on the relationships of some subfamilies and families, it is insufficient to accurately resolve the deep evolutionary relationships within Coleoptera. As exemplified in the most comprehensive studies based on morphology^20^, eight gene markers^16^, mitochondrial genomes^21,22^, and more recent analyses of phylogenomic datasets^10,23^, and transcriptomes^23^, the interrelationships of the suborders and most series and superfamilies of the most diverse beetle suborder, Polyphaga, still lack consistency and sufficient statistical support; as such, relationships among many families remain effectively unresolved^19^. Compositional and rate heterogeneity are among the most common sources of phylogenetic incongruence^24–26^. Reducing site compositional heterogeneity in datasets combined with the utilization of evolutionary models incorporating compositional and rate heterogeneity, such as the CAT model, has been shown to fit data better and suppress long branch attraction (LBA) artifacts, one of the most prevalent type of systematic error in phylogenies^27,28^. Unlike site-homogeneous models, such as the widely-used WAG and LG models utilized in past analyses of coleopteran phylogeny, site-heterogeneous models such as the CAT model relax the assumption that the overall rate of substitution across sites in a sequence is constant and that substitution at different positions in an alignment occur independently^29^. As such, heterogeneous models provide a more faithful reflection of molecular evolution and commonly fit analysed datasets better than computationally simpler site-homogeneous models^30^. Consequently, site and rate heterogeneous models have been widely used to resolve complicated phylogenomic problems, such as deep and rapid radiations that are otherwise difficult to recover^30–33^. In different groups of beetles, site-heterogeneous models have recovered topologies that are highly-congruent, sometimes identical, to traditional morphology-based classification schemes^21,25,34^, thus contributing to resolving the often perceived ‘conflict’ between morphological and molecular phylogenies^35^. However, the site-heterogeneous CAT model has not been used widely for the analysis of protein-coding sequences of beetles.

Reconstructing the timescale of beetle evolution also has to explicitly account for sources of error in molecular clock estimates and the biases of the fossil record. Despite the rich fossil record of beetles, molecular clock analyses have generally incorporated few fossil calibrations, ranging from 7–34^9,10,16,23,36^, a practice that may lead to inaccurate divergence time estimates^37,38^ as the congruence between molecular clock estimates and the fossil record tends to increase logarithmically with an increasing number of calibrations used^39^. The selection and phylogenetic placement of fossil calibrations is another significant factor influencing divergence dates^40,41^ and past studies have either not followed best practices in justifying the phylogenetic position and stratigraphic age of the fossils^42^, or used fossils to calibrate more derived nodes than they truly represent^9,10,16^, skewing divergence time estimates. Finally, the choice of arbitrary maximum age constraints on nodes introduces uncertainties in inferring divergence dates^43^, which remain unquantified for beetles. Fossils can only provide minimum constraints on clade ages^44^, which is particularly important for taxa such as beetles that are restricted to a small number of fossil deposits with exceptional preservation^45^, making the criteria used to define maximum constraints on nodes are imposed an important consideration for molecular clock studies. These sources of bias have contributed to uncertainties in dating key events in beetle evolution, such as the date of origin of the beetle clade or the timing of Polyphaga radiation relative to the diversification of angiosperms – a key tenet of the Cretaceous Terrestrial Revolution hypothesis^11^.

Here we reconstruct a novel timescale of beetle evolution that integrates the fossil record and refined sampling of 68 nuclear protein coding (NPC) genes (16,206 amino acid sites) generated from Zhang et al.^10,46^ with the addition of *Rhinorhipus*^47^. We established 57 new calibrations selected in accordance with best practice recommendations^42^ that are fully justified with respect to their stratigraphic age and systematic position, more than have been used in any previous analysis of the timing of beetle evolution. Objective maximum age constraints have been specified based on the absence of beetle taxa in well-explored fossil deposits. We provide a well-resolved phylogeny of beetles with a comprehensive taxon sampling based on the site-heterogeneous CAT-GTR+G4 model. This phylogeny is more consistent with morphological data^20,25^ and whole genome analyses^23^ and we use it to propose formal changes to the classificatory scheme of beetles.

## Results

### Systematic bias

Prior to phylogenetic reconstruction, we subjected our 68-gene dataset to tests to determine possible sources of systematic error. Of the 376 analysed taxa, 115 failed the compositional homogeneity test (*p*<0.05) (Supplementary Table 1 available on Mendeley Data). Moreover, the dataset displayed a RCFV value of 0.0229, which is regarded as ‘high’ (>0.1)^48^. High RCFV values are indicative of pronounced amino acid compositional heterogeneity, which is characteristic of fast-evolving taxa^49^ (long branches) and may lead to misleading phylogenetic inference^24–26^. This was confirmed in preliminary analyses with reduced taxon sampling which showed that the recovery of relationships at deep nodes was heavily influenced by the model used in phylogenetic reconstruction. The site-homogeneous models LG, WAG, GTR20, and JTT recovered topologies with high statistical support, but differed profoundly from the better fitting site-heterogeneous model C10 (Tab. S1), with respect to the placement of numerous polyphagan groups, namely within Cucujiformia (Figs S1–S5). The recovered model-dependent phylogenetic signal is strongly suggestive of systemic incongruence of phylogenetic signal, attributable in part to LBA artifacts and other sources of systematic error such as the heterogeneity of the substitution process. Hence, we expect systematic bias to be an important source of incongruence in beetle phylogenomics, corroborating previous phylogenomic studies of beetle mitochondrial genomes^25,34,50^.

### Phylogenomic reconstruction

To overcome bias associated with compositional heterogeneity, we tested site-homogeneous and site-heterogeneous models for their fit to the whole analysed dataset with 376 taxa by running a cross-validation analysis in PhyloBayes. The site-heterogeneous model CAT-GTR+G4 fitted the dataset better than the site-homogeneous GTR model, variations of which have been used in some previous analyses of beetle phylogeny (CAT-GTR+G4> GTR; CV score = 8,094.35 ± 341.776). This is in line with previous studies that show that site-heterogeneous models consistently outperform site-homogeneous ones, albeit at the expense of a greater computational burden^30,33^. As such, we present the topology reconstructed by the site-heterogeneous model CAT-GTR+G4 implemented in PhyloBayes as our main tree. Topologies recovered with the site-homogeneous model (GTR20, Figs S7) and a variant of the site-heterogeneous CAT model (LG+C20, Figs S8), varied with respect to the support of key deep clades and recovered relationships. For example, the GTR and LG+C20 models recovered the suborders Archostemata and Myxophaga as sister to Adephaga, an alternative branching order of superfamilies within Cucujiformia inconsistent with recent analyses^10,16,23^, and differed with respect to the relationships among some polyphagan families such as Boganiidae. Nonetheless, the monophyly of major series and superfamilies has been supported by the site-homogeneous models as well as CAT-GTR+G4, which is further discussed below.

Our PhyloBayes analyses with the CAT-GTR+G4 model recovered deep relationships among suborders with Adephaga as the earliest diverging lineage, Adephaga (Polyphaga (Myxophaga, Archostemata)), a novel hypothesis^10,16^, albeit with low support (Fig. 3). As noted in Zhang et al.^10^, the two smallest coleopteran suborders, Archostemata and Myxophaga, were insufficiently sampled (each represented by a single taxon) and this may, in part, account for such a result. Despite extensive morphology-based phylogenetic studies^51–53^, the intersubordinal relationships should be considered tentative, awaiting future phylogenomic studies^19^. Consistent with Zhang et al.^10^, McKenna et al.^23^, and an analysis of ultraconserved elements^54^ within Adephaga, we recovered a paraphyletic ‘Hydradephaga’ with the aquatic Haliplidae and Gyrinidae forming a clade sister to other aquatic and terrestrial adephagans. A paraphyletic ‘Hydradephaga’ with Gyrinidae as the earliest diverging adephagan branch was also recovered by a number of morphological analyses of adults and larvae^55^.

Among Polyphaga, five families formed three branching clades sister to the remaining polyphagan families (Fig. 2), in congruence with all recent studies^9,10,16^. As in previous analyses^9,10,16,23^, Derodontidae was recovered as sister to Clambidae + Eucinetidae. In contrast to Zhang et al.^10^, our analysis also included the enigmatic Australian endemic family Rhinorhipidae (superfamily Rhinorhipoidea), sequenced by Kusy et al.^47^, which was strongly supported as sister to the remaining polyphagans. Thus, we can reject the recent hypothesis that it may be sister to Nosodendridae or to Elateriformia based on mitogenomes and Sanger sequence data analysed under site-homogeneous models^47^. This finding is congruent with the genome-scale analysis of McKenna et al.^23^ and selected analyses of Kusy et al.^47^.

**Fig. 1.**
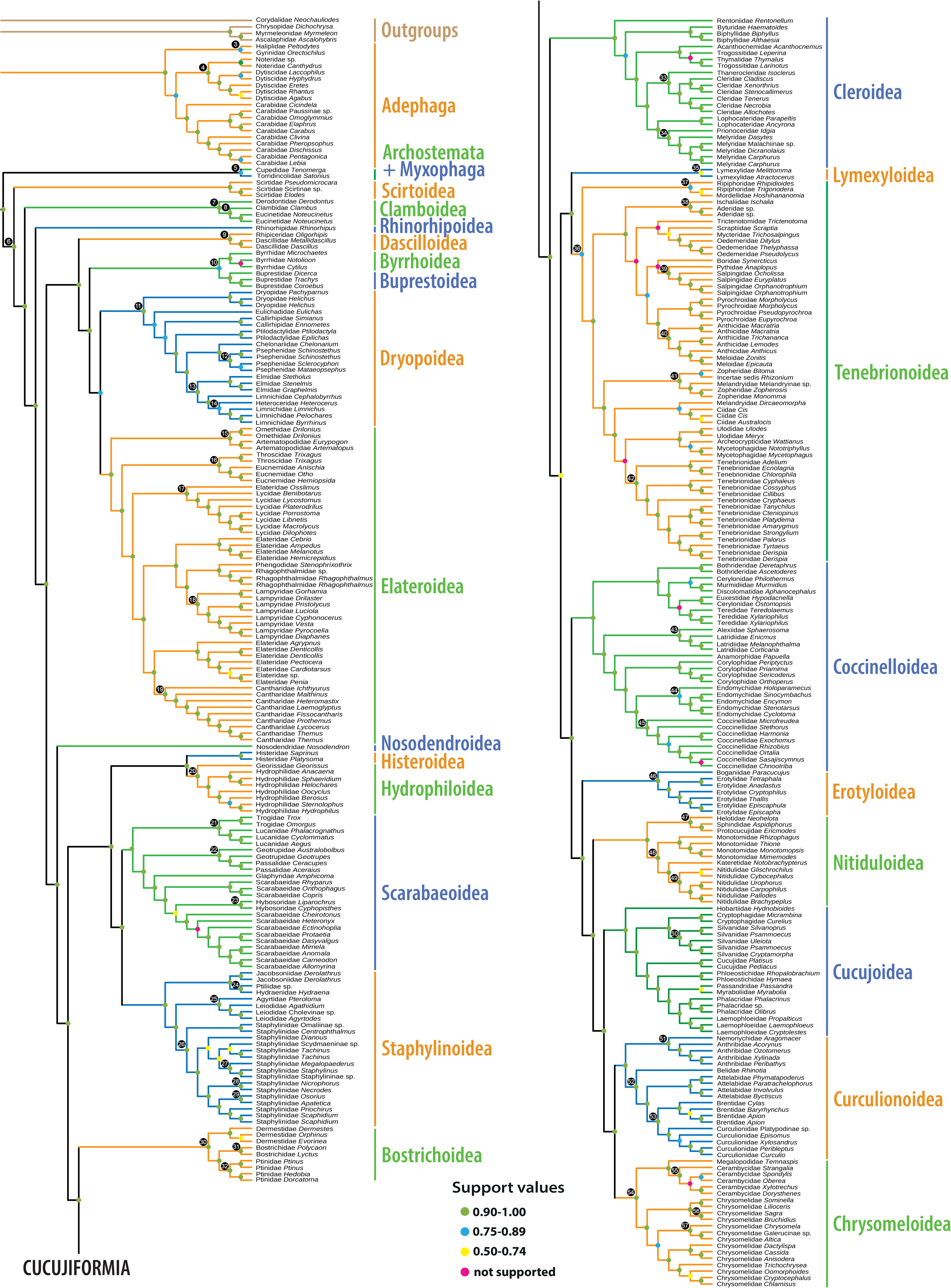
A full phylogeny of beetles displaying the systematic position of all sampled taxa analysed under the site heterogeneous CAT-GTR+G4 model. Branch lengths have been omitted for clarity. Newly proposed taxonomic changes are followed. Support values are shown as Bayesian posterior probabilities (BPP). Black numbered nodes indicate calibrations, see Supplementary Information for full list of calibrations.

**Fig. 2.**
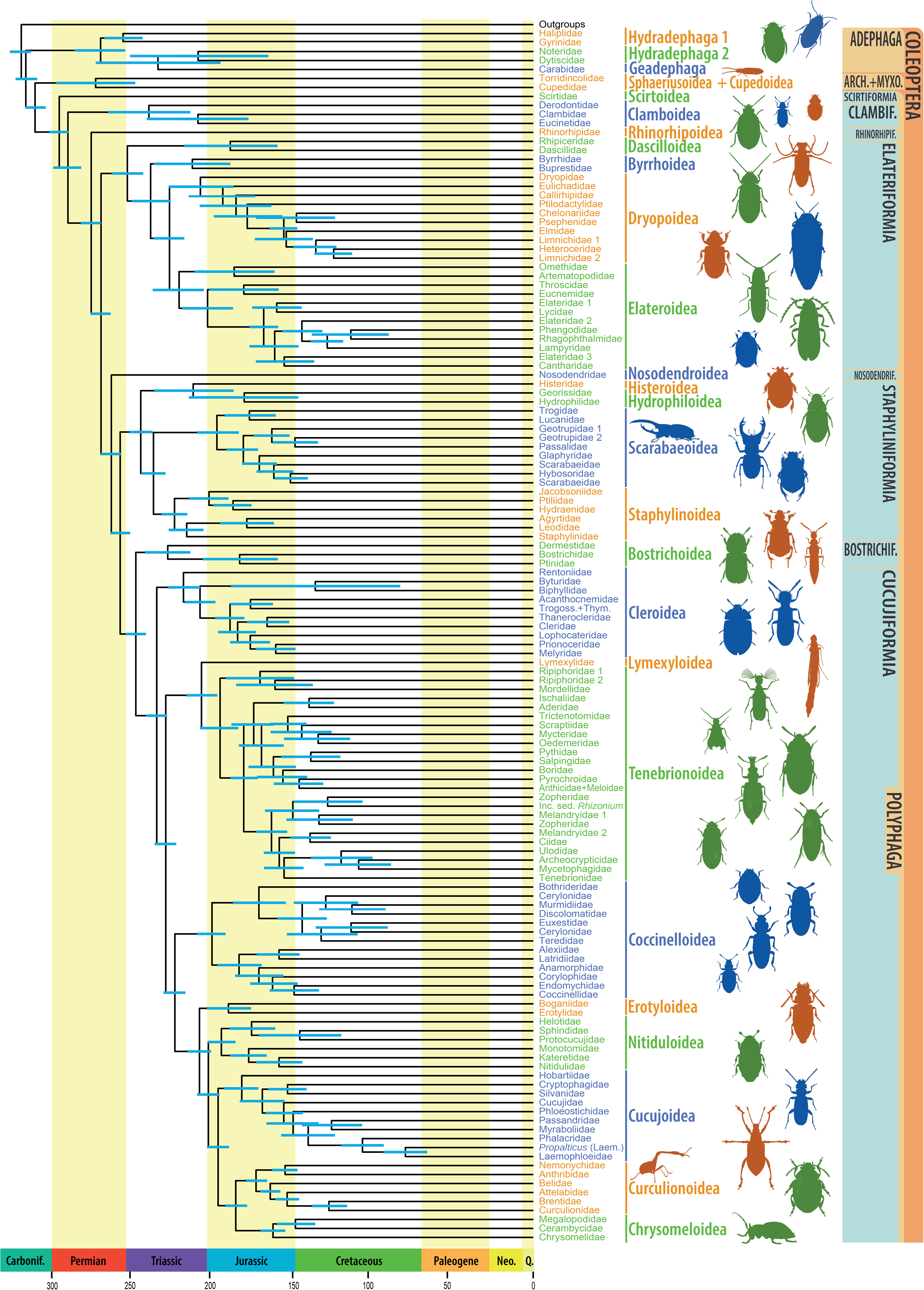
Timescale of beetle evolution displayed as a family-level tree adapted from Fig. 1. Ages were estimated based on 57 calibrated nodes, integrating the results of analyses using independent rates (IR) and autocorrelated rates (AC) molecular clock models in MCMCTree. Newly proposed taxonomic changes are followed. Abbreviations: Ar., Archostemata; Bostrichif., Bostrichiformia; Carbonif., Carboniferous; Clambifor., Clambiformia *ser. nov.*; Laem., Laemophloeidae; Myxo., Myxophaga; Neo., Neogene; Nosodendrif., Nosodendriformia *ser. nov.*, Rhinorhipif., Rhinorhipiformia *ser. nov.*; Trogoss. + Thym., Trogossitidae + Thymalidae; Q., Quaternary.

Next to Rhinorhipidae, the monophyletic series Elateriformia was recovered as the fourth polyphagan branch with strong support, a result consistent with genomic analyses^21,22,47^, rejecting a more derived position based on an eight-gene phylogeny^16,19^. Within Elateriformia, Dascilloidea were corroborated as the sister group to the remaining elateriform superfamilies, as in Zhang et al.^10^, Kusy et al.^56^, and McKenna et al.^23^, rejecting a weakly supported and ambiguous Dascilloidea-Byrrhidae relationship in Timmermans et al.^21^. The poorly supported monophyletic Byrrhoidea in Zhang et al.^10^ and McKenna et al.^16^ was not supported here, with Byrrhidae forming a sister group to Buprestidae. A polyphyletic Byrrhoidea was also recovered based on mitogenomes^21^, whole genomes^23^, and a small number of genes^9^, in which the name Dryopoidea was used for defining Byrrhoidea minus Byrrhidae, and Byrrhoidea *s. str.* for the moss-feeding Byrrhidae. As supported by numerous plesiomorphic characters (especially larval traits), Byrrhoidea *s. str.* and Buprestoidea have been long regarded to be isolated from other elateriforms^57–59^, in concert with our new topology. Additionally, Dryopoidea have been well defined by a unique rearrangement of tRNA gene order^60^. The interfamilial relationships among Dryopoidea are exactly the same as those based on a site-homogeneous model^10^, providing the best-supported relationships and clarifying uncertainties in the phylogeny of Dryopoidea based on a few genes^16,61^. Within Elateroidea,, the clade of ‘higher elateroids’ *sensu* Kundrata et al.^62^ was strongly supported, which is consistent with previous studies^9,10,16^. The clade Lycidae + Elateridae (Lissomini) was strongly supported to occupy a more basal position compared with Cantharidae, which is inconsistent with Zhang et al.^10^, but consistent with McKenna et al.^16^. Elateridae were recovered non-monophyletic as in other recent studies^10,16,23,63^.

Next to Elateriformia, Nosodendridae was strongly supported as sister to the remaining polyphagans (Staphyliniformia, Bostrichiformia, and Cucujiformia), in congruence with McKenna et al.^23^ and rejecting its placement within Elateriformia^9,16,23,62^.

Monophyly of the clade encompassing Histeroidea, Hydrophiloidea, Staphylinoidea (including Jacobsoniidae^10,16^), and Scarabaeoidea, collectively known as Haplogastra, was strongly supported (BPP = 1), consistent with various previous studies^10,16,21,64,65^. These results are congruent with the recent molecular clock analysis focused on Staphylinoidea that used 50 calibrations^66^. Histeroidea + Hydrophiloidea was recovered as sister to Staphylinoidea + Scarabaeoidea (BPP = 1), resolving challenging relationships difficult to reconstruct based on morphology^64^, mitogenomes^21^, or few genes^16,67^. Bostrichiformia were recovered as a sister group to Cucujiformia, corroborating previous results^16,21^.

Within Cucujiformia, the superfamily Cleroidea (including Biphyllidae and Byturidae), instead of Coccinelloidea^10,16^, was recovered as the earliest-diverging clade (BPP = 1). Lymexyloidea + Tenebrionoidea was moderately supported as sister to the rest of Cucujiformia (BPP = 0.66). Coccinelloidea formed a sister group to Cucujoidea *sensu* Robertson et al.^68^ + Phytophaga (Curculionoidea and Chrysomeloidea) (BPP = 1), representing a relationship never recovered before^9,10,16,21,68^. Intriguingly, the monophyly of the superfamily Cucujoidea *sensu* Robertson et al.^68^ was not supported, consistent with other phylogenomic studies^21,23^, but not recent studies based on site-homogeneous models (e.g. ref.^10,16,68^). The clade Boganiidae + Erotylidae was strongly supported as a sister group to other cucujoid families plus Phytophaga (BPP = 1). Helotidae, Sphindidae, Protocucujidae, Monotomidae, and the Nitidulidae group (Kateretidae, Nitidulidae, and Smicripidae) formed the next branch (BPP = 1). Within this clade, Cybocephalinae *stat. nov.* were recovered as members of Nitidulidae, as defined by morphology^69^, rejecting their previously proposed status as a separate family^10,70^. The remaining cucujoid families formed a monophyletic group, with Hobartiidae being the first branching lineage (BPP = 1). Monophyly of Phytophaga was strongly supported (BPP = 1), congruent with recent phylogenies^10,16,21^, but see^9,18^. Belidae were recovered as the second branch within Curculionoidea (BPP = 1), rejecting previous hypotheses that recovered them as a sister group to Nemonychidae + Anthribidae^71^ or to all other curculionoids^16^.

### Fossil calibrations

We derived 57 calibrations spread throughout the tree of beetles based on a combination of fossil, phylogenetic, stratigraphic, geochronological, and biogeographic evidence. Fossil calibration choice was conducted conservatively following best-practice^44^. Care was taken to calibrate nodes distributed equitably throughout the tree, informed by fossils that are the earliest phylogenetically-secure members of their respective clades based on the presence of unambiguous synapomorphies. Soft maximum node constraints were based on the absence of fossils in well-studied Lagerstätten to circumvent the use of arbitrary maxima.

For example, we selected *Ponomarenkium belmonthense* from the late Permian Newcastle Coal Measures at Belmont, Australia, to calibrate the node representing crown Coleoptera^72^. The type series is deposited in a public institutional collection, the Australian Museum in Sydney (holotype, 40278; paratype, 41618)^72^. The fossils possess open procoxal cavities, a narrow prosternal process, and lack a broad prothoracic postcoxal bridge; these characteristics place them into crown-Coleoptera. However, the combination of characters present in *Ponomarenkium* excludes it from the crown group of all four extant beetle suborders: internalized metatrochantin, metanepisternum only marginally part of the closure of the mesocoxal cavity, elytra without window punctures, antennae moniliform (Archostemata); transverse ridge of the mesoventrite, short metacoxae not reaching the hind margin of abdominal sternite III, absence of coxal plates (Adephaga); absence of a broad contact between the meso- and metaventrites (Myxophaga); and exposed propleuron (Polyphaga)^72^. The age of the Tatarian insect beds in the Newcastle Coal Measures at Belmont, from which the fossils originate, is derived from stratigraphic correlation and high-precision CA-TIMS U-Pb zircon dating and hence a conservative minimum age for the fossil can be taken from the top of the Changhsingian, dated at 251.902 Ma ± 0.024 Myr^72,73^, thus, 251.878 Ma. The maximum constraint on the root of Coleoptera was 307.1 Ma, based on the age of the Mazon Creek Lagerstätte^74^. Despite its highly diverse fossil insect assemblage that has been intensively studied for decades^106^, no unequivocal beetles are known from Mazon Creek or other younger Carboniferous insect Lagerstätte such as Commentry in France. The interpretation of *Adiphlebia* from Mazon Creek as the earliest beetle^75^ has been shown to be erroneous, based on clay artefacts attached to the wing membrane^76^. Full details of all 57 calibrations used are provided in the Supplementary Information. This comprehensive set of fully-justified calibrations serves as a basis for future divergence time analyses in Coleoptera, as well as our own. The suitability of our specified priors was evaluated by running an analysis without molecular data, yielding the effective priors for comparison^77^.

### Divergence time estimates

To derive an integrative timescale of beetles, we combined the results of alternative molecular clock models: independent rates (IR) and autocorrelated rates (AC). Both models yielded comparable results, with the IR clocks proposing earlier dates on average (Figs S9, S10).

We recovered a late Carboniferous origin of Coleoptera (322–306 Ma). This inferred divergence time is somewhat older than the age of the earliest unequivocal stem-group coleopteran, *Coleopsis archaica* from the Sakmarian Meisenheim Formation in Germany (~290 Ma)^5^, which has not been used as a calibration point. Despite a controversial Carboniferous record of putative stem Coleoptera and Coleopterida (i.e., Coleoptera + Strepsiptera)^66,67^ that have been recommended for calibration in some studies^103,104^, the affinities of these fossils have been questioned as they do not preserve unambiguous coleopteran apomorphies^4,105^.

Crown Adephaga are estimated to have originated in the latter half of the Permian (282–257 Ma). The earliest diverging adephagan family, Gyrinidae, originated in the Permian–Triassic. Adephagans colonized land once, between the Triassic and the Late Jurassic (248–162 Ma), when the clade including ground beetles and tiger beetles (Geadephaga) diverged from their aquatic ancestors.

The suborders Adephaga and Myxophaga diverged between the Pennsylvanian and mid-Triassic (314–236 Ma), in congruence with most other recent views of the group’s evolution (Fig. 4).

Polyphaga was estimated to have originated in a relatively narrow interval between the latest Carboniferous and Early Permian (307–286 Ma). These dates are significantly older than those estimated by previous studies^9,10,16^, which inferred a Permian to Triassic origin of the clade, but fall within the range estimated by Toussaint et al.^36^ and McKenna et al.^23^. The two earliest diverging coleopteran clades including Scirtidae and Clambidae diverged in the Late Triassic–Late Cretaceous (227–77 Ma) and Permian–latest Triassic (286–202 Ma), respectively. Despite their position on the polyphagan tree, these clades are scarcely represented in the pre-Paleogene fossil record^78^, which may contribute to their broad credibility intervals. The lineage comprising the isolated family Rhinorhipidae was inferred to be Permian in age (287–264 Ma).

The species-rich series Elateriformia diverged in the Late Permian to Middle Triassic (269–246 Ma). The last common ancestor of the Histeroidea, Hydrophiloidea, and Staphylinoidea clade diverged in the Early to Late Triassic (258–238 Ma). Staphylinidae, the most diverse beetle family^79^, originated between the Late Triassic and Early Jurassic (209–184 Ma), and subsequently diversified in the Early Jurassic to Late Cretaceous. The origin of Cleroidea was estimated as Late Triassic (236–209 Ma). The Lymexyloidea + Tenebrionoidea clade diverged between the Late Triassic and earliest Jurassic (227–201 Ma), with most families subsequently diverging in the Jurassic and Early Cretaceous. Coccinelloidea diverged in the Late Triassic to Early Jurassic (217–182 Ma). Major derived cucujiform branches originated roughly contemporaneously; the early diverging Boganiidae + Erotylidae clade is Late Triassic to Middle Jurassic (210–167 Ma) in age, Helotidae and kin are Late Triassic to Early Jurassic (210–180 Ma), and the most derived clade including Hobartiidae and relatives is Early to Middle Jurassic (198–164 Ma), with the lineages of most extant families originating in the Jurassic to Early Cretaceous. Phytophaga dates back to the Early to Middle Jurassic.

## Discussion

### Taxonomic implications

Molecular phylogenetic studies conducted over the last decade have lent support to several deep relationships among coleopteran groups suspected by earlier workers on the basis of morphological data alone. Considering that some of these relationships are strongly supported in different molecular studies^10,19,21,23,67^, and are thereby robust to taxon and gene sampling as well as to LBA artefacts, we herein propose several changes to the taxonomy of Coleoptera. All taxonomic changes, including morphological diagnoses, lists of included taxa, and detailed commentaries are provided in the Supplementary Information and are briefly summarized below. In total, 234 families are recognized of which 194 are extant. A hypothesis of the relationships among the higher taxonomic units within Coleoptera is outlined in Fig. 3.

**Fig. 3.**
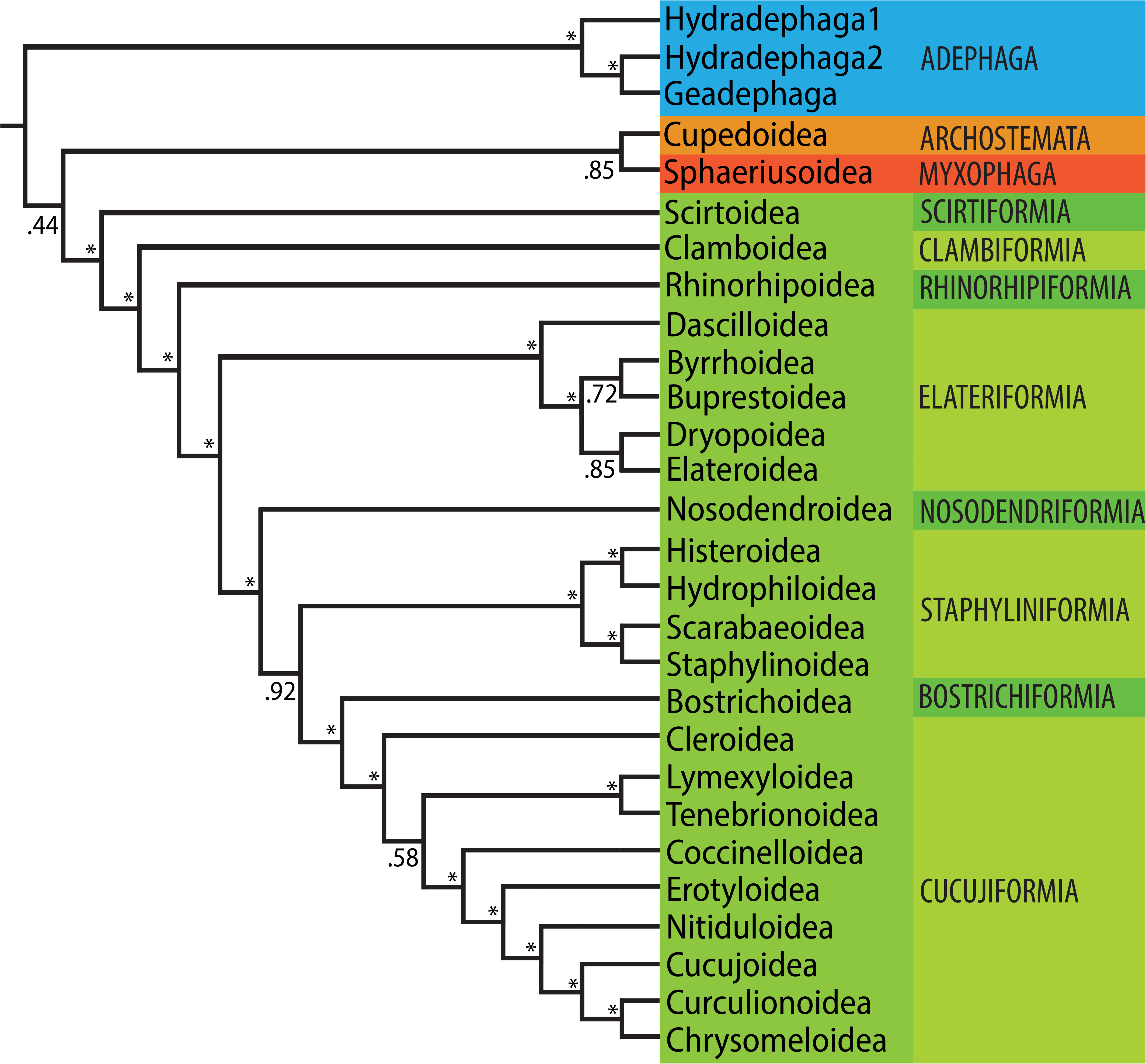
Proposed classification of Coleoptera showing the relationships of the suborders, series, and superfamilies of beetles. Asterisks (*) denote well-supported nodes with BPP ≥ 0.95.

**Fig. 4.**
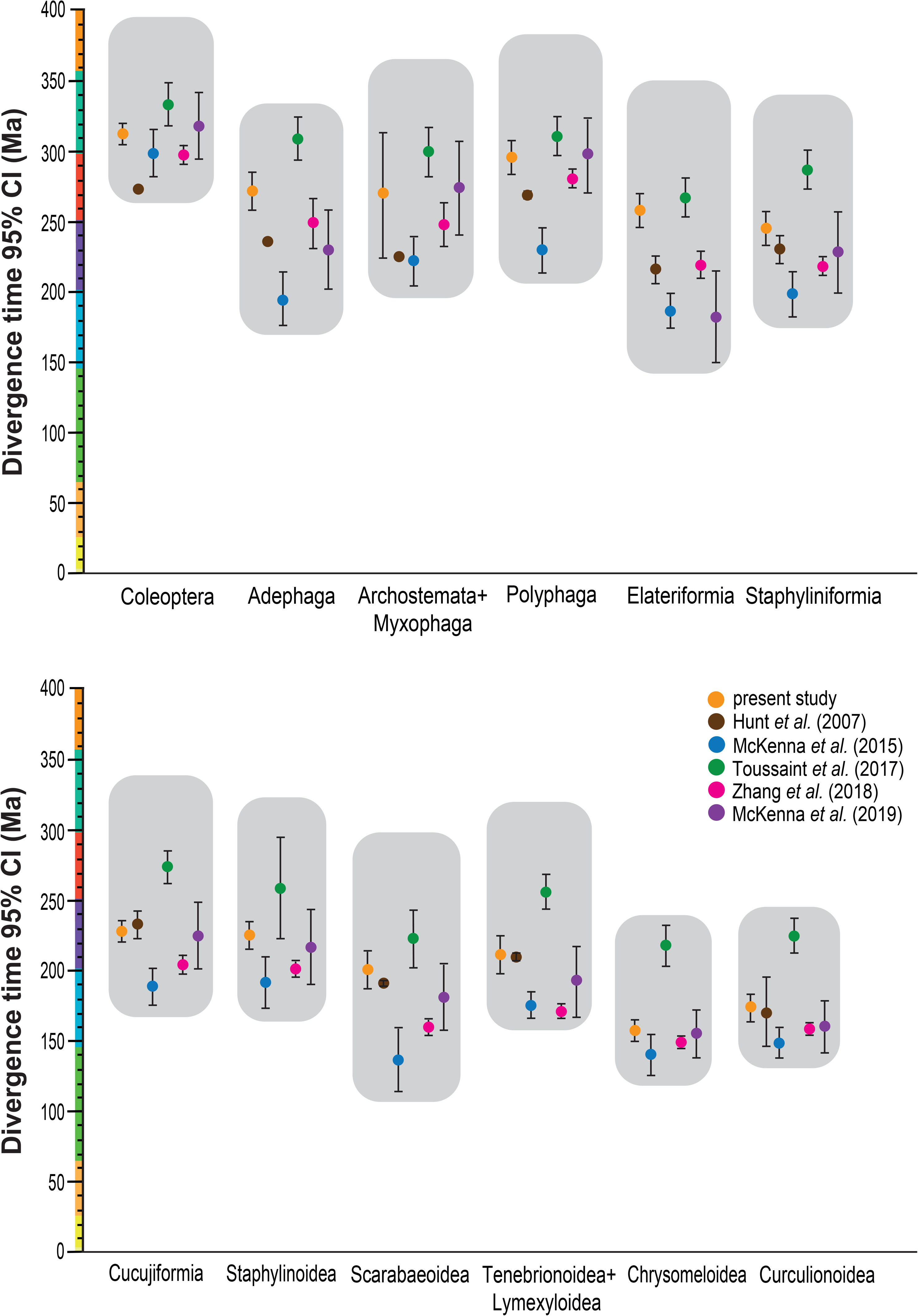
95% credibility intervals (CIs) for (a) the divergence of selected major beetle clades in the present and previous studies.

The traditional series Derodontiformia, classically defined as containing the three problematic families Derodontidae, Nosodendridae, and Jacobsoniidae, does not form a natural group in our analyses and other previous phylogenomic studies^10,23^. The monophyly of the group has been doubted based on morphological characters as well^20,58,80^. We consider Jacobsoniidae a member of Staphylinoidea, and Nosodendridae as belonging to a series of its own. Consequently, Scirtiformia and Scirtoidea *sensu nov*., and Clambiformia Cai and Tihelka *ser. nov.* and Clamboidea *sensu nov.* (with Clamboidea Fischer, 1821 taking priority over Derodontoidea LeConte, 1861) are redefined, the former comprising the extant families Decliniidae and Scirtidae, and the latter Clambidae, Derodontidae, and Eucinetidae.

The enigmatic Rhinorhipidae, represented in the recent fauna by only a single species endemic to northeastern Australia, are resolved with strong support as an isolated branch falling outside of Elateriformia (where it was placed by Lawrence and Newton^81^). We propose to include the family in a series of its own, Rhinorhipiformia Cai, Engel and Tihelka *ser. nov.*

Byrrhoidea in the current sense was recovered as paraphyletic, with Byrrhidae forming the sister group to Buprestidae and the remainder of the superfamily forming a sister group to Elateroidea. This split roughly corresponds to Crowson’s concept^57^ of Byrrhoidea as containing only a single family, Byrrhidae, and is not unexpected as the monophyly of Byrrhoidea has been doubted^23,61^. Thus, we regard Byrrhidae as constituting Byrrhoidea *sensu nov*. and treat the remaining former byrrhoid families within Dryopoidea *stat. res.* Relationships among clades within Elateroidea are not resolved with sufficient support to justify taxonomic revision, which is consistent with the results based on the anchored hybrid enrichment method using 2,260 single-copy orthologous genes by Douglas et al.^63^.

In congruence with other phylogenomic analyses^10,23^, Nosodendridae were recovered as an isolated lineage sister to Staphyliniformia, Bostrichiformia, and Cucujiformia, outside of Clambiformia *ser. nov.* where the family was provisionally placed by previous workers^58^. To maintain the monophyly of the remaining well-defined series, and considering the long-established morphological distinctiveness of the family, a new series is instituted for Nosodendridae, Nosodendriformia Cai and Tihelka *ser. nov*.

We recovered Scarabaeiformia (Scarabaeoidea) nested within Staphyliniformia, as a sister group to Staphylinoidea, in line with other genomic analyses^10,23^. As this would render the well-defined Staphyliniformia polyphyletic, we integrate Scarabaeiformia within Staphyliniformia *sensu nov*. This newly defined Staphyliniformia essentially corresponds to the Haplogastra concept proposed more than a century ago by Kolbe^82^ which was subsequently supported by a range of morphological studies of adult and larval characters (see Supplementary Information for full discussion).

The superfamily Cucujoidea is a diverse (~10,000 described species) and heterogeneous group of beetles, frequently considered the most problematic taxon in Coleoptera^83^. Its monophyly has been questioned and various authors in the past have suggested splitting it (see Supplementary Information for full review). In genomic analyses^10,23,84^, the group is consistently recovered as a grade consisting of three clades: (1) Boganiidae + Erotylidae; (2) Helotidae, Sphindidae, Protocucujidae, Monotomidae, Kateretidae, Nitidulidae, and Smicripidae; and (3) Hobartiidae, Cryptophagidae, Silvanidae, Cucujidae, Phloeostichidae, Passandridae, Myraboliidae, Phalacridae, Propalticidae (= Laemophloeidae), and Laemophloeidae. We consequently split the old Cucujoidea *sensu* Robertson et al.^68^ into three superfamilies: Erotyloidea *stat. nov.,* Nitiduloidea *stat. nov*., and Cucujoidea *sensu nov.* to provide a phylogenetically-sound, more balanced and practical classification of this diverse group that reflects its evolutionary history.

Our results support the downgrading of the carrion beetles from a family to a subfamily of Staphylinidae *sensu. nov*., as Silphinae *stat. nov.*, and reconsideration of other recent phylogenetic studies supports the restoration of Colonidae *stat. nov.* as a separate family, not subfamily of Leiodidae *sensu nov*. Moreover, click beetles (Elateridae), minute marsh-loving beetles (Limnichidae), earth-boring dung beetles (Geotrupidae), scarabs (Scarabaeidae), bark-gnawing beetles (Trogossitidae), ant-like beetles (Anthicidae), ironclad beetles (Zopheridae), and false darkling beetles (Melandryidae) were not recovered as natural groups, in congruence with some previous analyses^56,85^. These latter issues remain to be formally treated and will require a more extensive taxon sampling within and outside these groups to help redefine their boundaries.

### Carboniferous origin of Coleoptera

Despite a history of problematic purportedly coleopteran fossils from the Carboniferous^75,86^, the earliest unequivocal stem-group beetles appear in the fossil record in the early Permian^5^, providing few clues into the initial history of the order. Molecular clock studies have broadly converged on two principal models of beetle origins: the explosive model, suggesting that total-group Coleoptera originated and diversified in the Permian^9,10^, while the long-fuse model argues for a much earlier Carboniferous origin^36^, implying a long cryptic evolutionary history of the group undocumented in the fossil record. Our molecular clock studies lend support to the latter model, recovering a fully Carboniferous origin of Coleoptera and suggesting a 55–134 Myr gap in the fossil record between the origin of Coleoptera and the earliest fossil record of the order. The paucity of early beetle fossils may be because of the rarity of early beetles, their narrow ecological niche, lower fossilization potential, insufficient sampling of fossiliferous formations of this age, as well as secular biases in the rock record. The former two appear especially likely, since stem-beetles were rare in the Permian and coleopterans only came to dominate fossil insect assemblages in the Mesozoic^87^. Moreover, stem beetles were apparently not considerably more morphologically diverse than modern beetles^88,89^. The early Carboniferous is part of the ‘Hexapod gap’, a stratigraphic interval known for its lack of insect fossils^90^, which may further contribute to the lack a pre-Permian fossil record of beetles.

Adaptative radiations of successful clades in the fossil record are often associated with new morphological innovations that enable the colonization of new niches^91^. The heavily sclerotized bodies of beetles with forewings modified into hardened protective elytra are often cited to explain the diversification of beetles^8,9,14^. However, the long-fuse model, with a long cryptic history of Carboniferous–Permian beetles that apparently were neither abundant^87^ nor morphologically disparate^89^, argue for a more complex scenario of beetle diversification.

### Early diversification of beetles

The recovery of Adephaga as the earliest diverging clade of Coleoptera, and the aquatic families Gyrinidae and Haliplidae as the earliest diverging families within Adephaga, implies that the ancestral coleopteran may have been aquatic. This seems reasonable based on the fossil record because the Permian stem-coleopteran families †Tshekardocoleidae and †Phoroschizidae may have been fully or partly aquatic^92^ even though the life history of †Alphacoleoptera Engel, Cai, and Tihelka subord. nov. (refer to SI) remains mysterious^93^. Different phylogenetic reconstructions among the four suborders and the internal relationships of Polyphaga may refute the hypothesis for an aquatic ancestor^16^. Consistent with Zhang et al.^10^ and McKenna et al.^23^, Scirtiformia are resolved as the basal-most polyphagans and are a group with mainly aquatic and semiaquatic larvae^94^, though some forms are fully terrestrial^95^. Their placement supports the reconstruction of an aquatic ancestor for Archostemata + Myxophaga and the Polyphaga. An understanding of the phylogenetic relationships within scirtiforms^96^ is critical for testing the aquatic-ancestor hypothesis.

Under the aquatic ancestor hypothesis, beetles subsequently invaded terrestrial ecosystems independently four times, once each in the ancestor of Geadephaga, Archostemata, some Myxophaga, and Polyphaga, although many beetles have subsequently returned to freshwater again. The radiation of basal Coleoptera was rapid and occurred over a period of 17–78 Ma; the split between Archostemata and Myxophaga was estimated as Permian (269–246 Ma), while Polyphaga diverged at the Carboniferous–Permian boundary (314–236 Ma). The basal diversification of crown Coleoptera and their invasion of land occurred in the Permian, a time of drastic environmental change that saw the replacement of Carboniferous hygrophytic and permanently wetted terrestrial ecosystems inhabited by (semi)aquatic alphacoleopterans with seasonally dry ecosystems^97^. Paleozoic plant families were replaced by more modern Mesozoic groups and ecological communities became more complex, with a higher number of trophic levels^97,98^. It is therefore possible that the Permian ecosystem change played a pivotal role in shaping the early diversification of beetles^5,92^. Early diverging beetle lineages survived the End-Permian mass extinction event to diversify in the Mesozoic, most beetle superfamilies having diverged by the Jurassic.

### Cretaceous co-diversification with angiosperms

Angiosperms replaced the previously dominant gymnosperms during the Cretaceous, in a period known as the Cretaceous Terrestrial Revolution (KTR)^99,100^. Co-diversification with flowering plants has been proposed as a mechanism explaining the species-richness of herbivorous clades such as the weevils^9,15^ and beetles have been regarded as the earliest pollinators of angiosperms^15,101,102^. Our molecular clock analyses suggest that major beetle clades were present before the KTR. This is corroborated by the beetle fossil record12; elateroids, staphylinoids and weevils in particular have a diverse fossil record since the Jurassic^8,80^. Nonetheless, some scarabaeoid and cucujiform clades underwent diversification during the Late Jurassic to Early Cretaceous, partly overlapping with the diversification of major angiosperms clades in the Early to mid-Cretaceous^104,105^.

Besides directly affecting herbivorous lineages, the diversification of the angiosperms in the Cretaceous precipitated a diversification of vertebrate herbivores and predators^11^. Our molecular clock estimates corroborate the controversial idea, famously portrayed in numerous paleontological reconstructions, that coprophagous beetles, namely geotrupids (dung beetles) and scarabaeoids (scarabs), may have been associated with Cretaceous herbivorous dinosaurs^23,106,107^. Our analyses also corroborate a Cretaceous origin of the bioluminescent lampyroid clade^108^, temporally overlapping with the diversification of visually hunting predators such as anurans and stem-group birds during the KTR^109^. At the same time, some Mesozoic beetle families have their last appearance in the fossil record during the KTR, highlighting complex dynamics of transitioning from a gymnosperm- to angiosperm-dominated world^110^.

Overall, our results provide support for a more nuanced view^15^ of the KTR as an event that did not increase the superfamilial diversity of beetles, since most major beetle clades diverged by this time. Instead, the diversification of angiosperms was followed by clade-specific radiations in some beetle groups, such as scarabs^101^ or weevils^13^, in response to newly formed niches.

### K-Pg mass extinction

The impacts of the Cretaceous-Paleogene (K-Pg) mass extinction on beetles remain controversial, as herbivorous insects would be expected to have experienced elevated extinction levels given their close association with their host plants^111^. Our results suggest no diversification of beetles, at the level of families, in the aftermath of the K-Pg crisis, corroborating previous paleontological macroevolutionary studies suggesting that the mass extinction was hardly devastating for beetles^12,87,112^. It is however possible that the K-Pg extinction may have had different impacts on lower taxonomic ranks111 that have to be assessed by future studies focusing on specific herbivorous clades.

### Summary

Our new beetle phylogeny corroborates many relationships inferred by phylomitogenomic analyses using models accounting for compositional and rate heterogeneity^21^, but the resolution is significantly improved by a careful selection of single-copy NPC genes analysed with methods accounting for compositional and rate heterogeneity. We recovered with credibility the phylogenetic positions of the enigmatic Rhinorhipidae and recently recognized Coccinelloidea, as well as the paraphyly of Cucujoidea *sensu* Robertson et al.^68^. We also unraveled the isolated position of the peculiar Byrrhidae, consistent with the traditional classification of elateriform beetles^57,58,113^. Our molecular clock analyses suggest a Carboniferous origin of Coleoptera and a Paleozoic origin of all four beetle suborders. A Carboniferous origin of Coleoptera implies a 55–134 Ma long “beetle gap” in the fossil record. Early diverging beetle lineages survived the End-Permian mass extinction event to diversify in the Mesozoic, with major clades having originated by the Jurassic. Most major beetle clades were present by the Late Jurassic, although some groups diversified during the Cretaceous, in concert with the radiation of angiosperms.

We provide a revised treatment of the higher classification of Coleoptera that reflects the findings of phylogenomic studies conducted over the past decade. Scirtiformia and Scirtoidea *sensu nov*. are restricted to Decliniidae and Scirtidae, while Clambiformia Cai and Tihelka *ser. nov.* and its single constituent superfamily Clamboidea *sensu nov.* is considered to contain the extant families Clambidae, Derodontidae, and Eucinetidae. The other members of the former Derodontoidea, Nosodendridae and Jacobsoniidae, are placed into Nosodendriformia Cai and Tihelka *ser. nov.* and Staphylinoidea, respectively. To maintain the monophyly of the well-defined Elateriformia, we erect the new series Rhinorhipiformia Cai, Engel and Tihelka *ser. nov.* for the family Rhinorhipidae. Scarabaeoidea is formally incorporated into Staphyliniformia *sensu nov*. The former Cucujoidea is divided into three superfamilies: Erotyloidea *stat. nov.,* Nitiduloidea *stat. nov*., and Cucujoidea *sensu nov.* Silphinae *stat. nov.* are treated as a subfamily of Staphylinidae *sensu. nov*., and Colonidae *stat. nov.* is restored as an independent family and not a subfamily of Leiodidae. In addition, we recognize that the extinct suborder †Protocoleoptera (established for †Protocoleidae) comprises a group of polyneopterans allied to earwigs (Dermaptera) rather than beetles, and that the younger †Archecoleoptera were defined on the basis of a temporal fauna rather than any characters, taxa, or phylogenetic details. Accordingly, the earliest fossil beetles are here included in the new extinct suborder †Alphacoleoptera subord. nov. (refer to SI)

## Methods

### Dataset collation

We used the published NPC gene sequences from Zhang et al.^10^ supplemented with the *Rhinorhipus* sequences from Kusy et al.^47^. Zhang et al.^10^ presented and analysed both nucleotide and amino acid alignments of their data; their best tree was based upon a concatenated amino acid data set of 95 NPC genes. We excluded 27 genes that contain up to 21 copies in some beetle genomes and could not be homologized confidently^47^. All NPC genes were individually aligned using the “Translation Align” option with the FFT-NS-i-× 2 algorithm of MAFFT 7.2^114^. Ambiguously aligned regions were trimmed with BMGE 1.1 (-m BLOSUM30)^115^. The sequences were then concatenated into a supermatrix (376 taxa, 16,206 sites) using FASconCAT^116^. The concatenated supermatrix (68 NPC genes, 376 taxa) consisted of 16,206 amino acids. The dataset and output files are available from Mendeley Data (http://dx.doi.org/10.17632/7v27xcyv99.2).

### Compositional heterogeneity and model-specific artifacts

General statistics for the dataset, compositional homogeneity tests (*x*^2^ test of heterogeneity), and compositional heterogeneity (RCFV, relative composition frequency variability) were computed using BaCoCa v.1.105^117^. To assess the prevalence of model-specific incongruences in beetle phylogeny, we generated a reduced dataset with 136 taxa, sampling only a single representative for each family, and applied the site-homogeneous models LG, JTT, and WAG alongside the site-heterogeneous models C10 and C30. Analyses were conducted in IQ-TREE 1.6.3^118^ with 1,000 bootstraps. The fit of these models to the dataset was assessed using ModelFinder^119^ implemented in IQ-TREE 1.6.3.

### Model selection and phylogenetic analysis

We tested the impact of accounting for compositional heterogeneity on phylogenetic inferences of Coleoptera. We tested the site-heterogeneous infinite mixture model CAT-GTR+G4, which has been shown both theoretically and empirically to suppress long-branch attraction artefacts^24,27^, and the site-homogeneous model GTR20 and the model LG+C20, variants which have been used in recent phylogenomic studies of Coleoptera^10,16,23^. While the C20 model accounts for site-specific substitutional heterogeneity, just like the CAT model, it differs from the latter in that it offers only 20 substitutional categories, while the CAT model offers an unconstrained number of categories estimated from the data. To test whether the CAT-GTR+G4 model fits the decisive amino acid alignment better than the GTR model, a variant of which was used by Zhang et al.^10^, a Bayesian cross-validation analysis was run with ten replicates in PhyloBayes. CAT-GTR+G4 was run in PhyloBayes MPI 1.7^120^; two independent Markov chain Monte Carlo (MCMC) chains were run until convergence for up to 14 months. The models GTR20 and LG+C20 were implemented in IQ-TREE 1.6.3^118^ with 1,000 bootstraps.

### Fossil calibrations

We selected 57 calibrations spread throughout the tree of beetles based on a combination of fossil, phylogenetic, stratigraphic, geochronological, and biogeographic evidence (Tab. S2). Fossil calibration choice was conducted conservatively using coherent criteria by Parham et al.^42^. Care was taken to select fossils that are the earliest members of their respective clades and are distributed equitably throughout the tree. Soft maximum node constraints were based on the absence of fossils in well-studied Lagerstätten to circumvent the use of arbitrary maxima. The fossils were selected to calibrate nodes based on the presence of unambiguous synapomorphies.

### MCMCTree analysis

Divergence time estimation was performed using the approximate likelihood calculation in MCMCtree implemented in PAML 4.7^121^, incorporating softbound fossil calibrations on nodes on the tree^122^. We obtained 200,000 trees with a sampling frequency of 50 and discarded 10,000 as burn-in. Default parameters were set as follows: ‘cleandata=0’, ‘BDparas=1 1 0’, ‘kappa_gamma=6 2’, alpha_gamma=1 1’, ‘rgene_gamma= 2 20’, ‘sigma2_gamma=1 10’, and ‘finetune=1: 0.1 0.1 0.1 0.01 .5’. Convergence was tested in Tracer73^123^ by comparing estimates from the two independent chains. Analyses were run using both autocorrelated rate (AU) and independent rate (IR) clock models using uniform prior distributions to reflect ignorance of the true clade between the minimum and soft maximum constraints. In all analyses, the soft minimum and maximum bounds were augmented by a 2.5% tail probability. To ensure our priors were appropriate, we ran the MCMCTree analysis without sequence data to calculate the effective priors, compared to the specified priors.

**Tab. 1.** Divergence dates (95% CI of the posterior distribution of age estimates, in Ma) of the crown groups of beetle suborders and series from independent rates (IR) and autocorrelated (AC) molecular clock analyses with uniform prior distributions.

| **Clade** | **IR** | **AC** |
| --- | --- | --- |
| Coleoptera | 321–306 | 322–318 |
| Adephaga | 287–259 | 288–266 |
| Archostemata – Myxophaga | 286–236 | 314–285 |
| Polyphaga | 301–288 | 307–286 |
| Scirtiformia | 177–77 | 227–90 |
| Clambiformia | 254–202 | 286–239 |
| Rhinorhipiformia | 287–271 | 283–264 |
| Elateriformia | 269–246 | 268–249 |
| Nosodendriformia | 276–258 | 271–253 |
| Staphyliniformia | 258–241 | 257–238 |
| Bostrichiformia | 237–203 | 250–228 |
| Cucujiformia | 249–231 | 233–220 |

## Data availability

All analysed datasets and outputs are available from MendeleyData (http://dx.doi.org/10.17632/7v27xcyv99.2). All sequences are deposited in GenBank under the accession numbers provided by Zhang et al.^10^ and Kusy et al.^47^.

## Code Availability

No custom code has been generated during this study.

## Acknowledgements

We are grateful to Ladislav Bocak (Palacký University, Czech Republic) for comments on an earlier version of the manuscript. Support for the present study was provided by the Strategic Priority Research Program of the Chinese Academy of Sciences (XDB26000000 and XDB18000000), the National Natural Science Foundation of China (41672011 and 41688103), the Second Tibetan Plateau Scientific Expedition and Research (2019QZKK0706), and a Newton International Fellowship from the Royal Society awarded to C.C. and P.C.J.D. This study was partly supported by a Grant-in-Aid for JSPS Fellows (20J00159) given to S.Y. from the Japan Society for the Promotion of Science (JSPS), and the European Union’s Horizon 2020 research and innovation programme under the Marie Skłodowska-Curie grant agreement (764840 to D.P. and M.G.).

## Author contributions

C.C. and P.C.J.D. conceived and designed the project. C.C. conducted phylogenomic reconstruction, C.C. and E.T. selected calibrations, E.T. run molecular clock and cross-validation analyses. C.C. and E.T. drafted the manuscript to which M.G., J.F.L., A.S., R.K., S.Y., M.K.T., A.F.N., R.A.B.L., M.L.G., L.L., M.S.E., D.H., D.P. and P.C.J.D. contributed. All authors read and approved the final version of the paper.

## Additional information

Supplementary information is available for this paper at … Correspondence and requests for materials should be addressed to C.C. or P.C.J.D.

## Appendix 1.

Overview of the updated higher classification of Coleoptera.

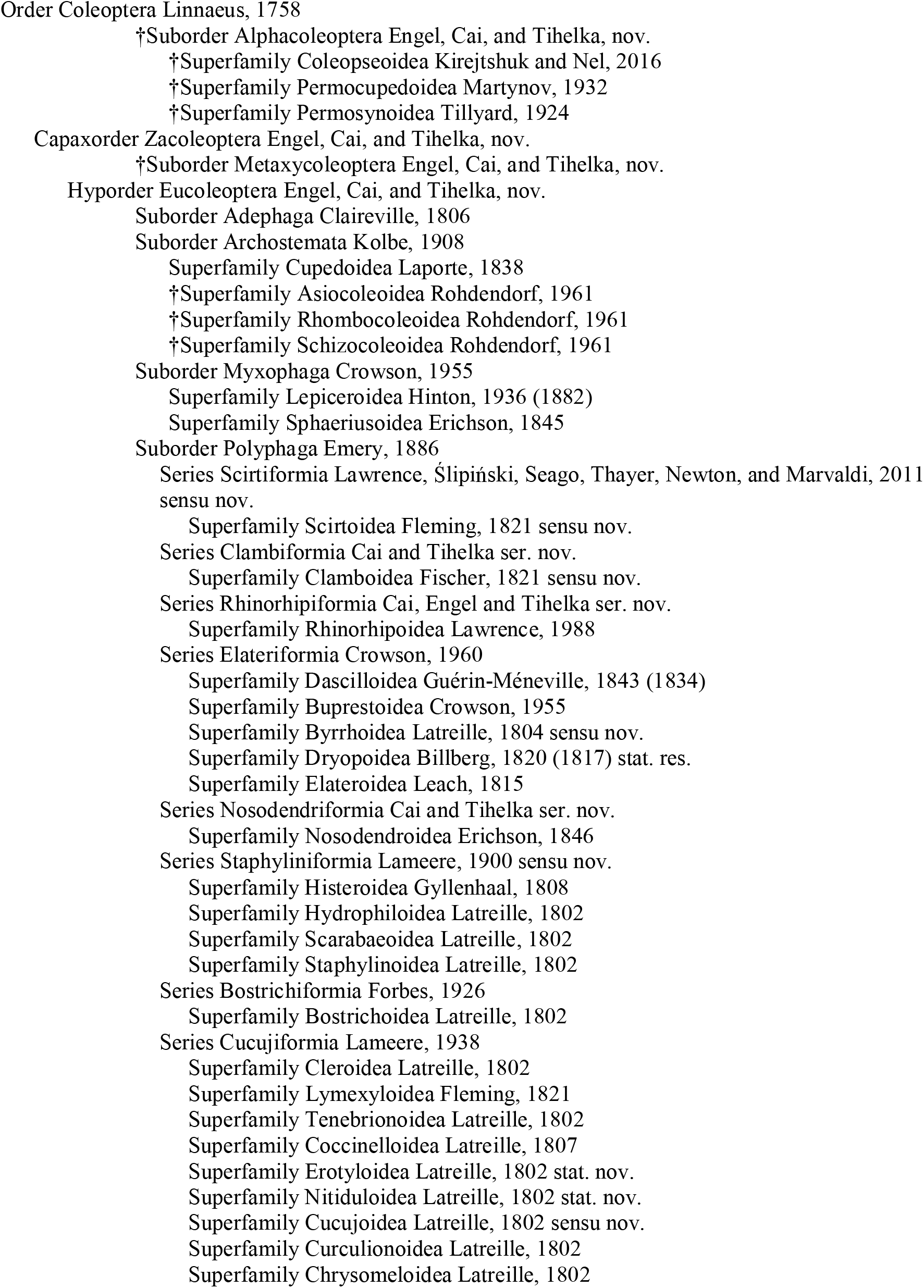

