## Supplemental Information for "Integrated phylogenomics and fossil data illuminate the evolution of beetles"

### Taxonomic treatment

SCIRTIFORMIA Lawrence, Ślipiński, Seago, Thayer, Newton and Marvaldi, 2011 / Superfamily SCIRTOIDEA Fleming, 1821 **sensu nov.**

***Type genus:*** *Scirtes* Illiger, 1807.

***Diagnosis:*** Adult head distinctly declined. Accessory ventral process of mandible present (partim). Ventral side of head with subgenal ridges. Prosternum very short in front of procoxae. Pterothoracic ventrites connected by membranes. Muscles *M. episterno-trochanteralis 2 (M. 70)*, *M. episterno-coxalis 3 (M. 103)*, and *M. furca-coxalis posterior 3 (M. 109)* absent. Functional spiracles on abdominal segment VIII absent. Hind wing AA4 joining anal fold posteriorly. Mesothoracic trochantins externally exposed. Metacoxal plates weakly to moderately enlarged. Larva with mandibular mola. Larval galea non-articulated.

***Included taxa:*** †Boleopsidae Kirejtshuk and Nel, 2013; Decliniidae Nikitsky, Lawrence, Kirejtshuk and Gratshev, 1994; †Elodophthalmidae Kirejtshuk and Azar, 2008; Scirtidae Fleming, 1821.

***Discussion:*** The series Scirtiformia and the superfamily Scirtoidea are herein restricted to only the extant families Scirtidae and Decliniidae, as well as the poorly-known extinct families †Elodophthalmidae from Lower Cretaceous Lebanese amber and †Boleopsidae from lower Eocene French (Oise) amber, whose placement however remains preliminary (Kirejtshuk and Azar 2008; Kirejtshuk and Nel 2013; Lawrence 2019a). The remaining families formerly affiliated with Scirtoidea are removed and placed into a redefined Clambiformia *ser. nov*. (which includes the single superfamily Clamboidea *sensu nov*.).

The basal position of the families Clambidae, Decliniidae, Derodontidae, Eucinetidae, and Scirtidae within Polyphaga was long recognised (Crowson 1955, 1960; Lawrence and Newton 1982; Lawrence et al. 1995b). However, disentangling the relationships among these families has proved challenging. Given that phylogenomic studies (Zhang et al. 2018; McKenna et al. 2019) have now firmly established Scirtidae as the earliest diverging polyphagan lineage, this renders the traditional series Scirtiformia and the superfamily Scirtoidea *sensu* Lawrence et al. (1995b) polyphyletic with the traditional Derodontoidea nested within it, thus mandating a taxonomic change to establish the two as natural groups reflecting evolutionary relationships. Adult Scirtidae are distinguished from Clambiformia *ser. nov*. by an enlarged metacoxal meron and a rounded radial cell of the hind wing, both of which may represent apomorphies of the family. Scirtids further have cone-shaped, distinctly elongated mesocoxae, an elongated metafurcal stalk, and have separated *M. noto-epimeralis (L 34)* and *M. epimero-subalaris (M. 52)* (Friedrich and Beutel 2006). The larval mandible of Scirtidae bears three putative apomorphies: flattening, suppression of mandibular teeth, and presence of a strong ventral hook at base (Zwick and Zwick 2008).

It should be noted that the position of the rare and monogeneric Decliniidae endemic to eastern Russia and Japan, not sampled in the present analysis, is not fully settled. It was consistently recovered as sister to Scirtidae in morphological (Lawrence et al. 2011) and molecular analyses of six genes (McKenna et al. 2015b). Their sister relationship is supported by the hind wing $AA_4$ joining anal fold posteriorly, a character otherwise found only in Adephaga. Adults of both families also lack functional spiracles on the eighth abdominal segment (Lawrence et al. 2011). Contrary to this arrangement, Anton et al. (2016) pointed out two potential synapomorphies of Clambidae and Decliniidae: a distinct ventral antennal furrow and a longitudinal carina on the laterodorsal surface of the mandibles. It is clear that the assessment of the systematic position of Decliniidae will require the sequencing of more genes and further morphological studies, especially of adult musculature. For now, the position of the family should be considered as tentative.

CLAMBIFORMIA Cai and Tihelka **ser. nov**. / Superfamily CLAMBOIDEA Fischer, 1821 **sensu nov.**

***Type genus:*** *Clambus* Fischer von Waldheim, 1821.

***Diagnosis:*** Adult head distinctly declined (prognathous in Derodontidae). Accessory ventral process of mandible present (partim). Ventral side of head with subgenal ridges (lost in Derodontidae). Prosternum very short in front of procoxae. Pterothoracic ventrites connected by membranes. Muscles *M. episterno-*

*trochanteralis 2 (M. 70)*, *M. episterno-coxalis 3 (M. 103)*, and *M. furca-coxalis posterior 3 (M. 109)* absent. Functional spiracles on abdominal segment VIII present. Hind wing AA4 not joining anal fold posteriorly. Incision between anal field and medial field very deep (in Derodontidae) to moderately deep (in remaining families). Mesothoracic trochantins exposed (except in *Eucinetus*). Metacoxal plates weakly to greatly enlarged. Larva with mandibular mola. Larval galea non-articulated.

***Included taxa:*** Clambidae Fischer, 1821; Derodontidae LeConte, 1861; Eucinetidae Lacordaire, 1857; †Mesocinetidae Kirejtshuk and Ponomarenko, 2010.

***Discussion:*** The series Clambiformia *ser. nov*. and the superfamily Clamboidea *sensu nov*. as defined herein include Clambidae, Derodontidae, and Eucinetidae, previously placed into the non-monophyletic Scirtiformia / Scirtoidea and Derodontiformia / Derodontoidea *sensu* Lawrence and Newton (1982), Lawrence (1991), Beutel (1996) and Lawrence et al. (2010). When Derodontoidea was defined by Lawrence and Newton (1982), it was understood primarily as a taxon of convenience for Derodontidae and the two other problematic families, Nosodendridae and Jacobsoniidae, and was noted to be "almost certainly paraphyletic". The taxonomic position of the two above-mentioned families was indeed phylogenetically exceptionally problematic and they were historically placed in a range of positions within Polyphaga, sometimes as *incertae sedis* (reviewed in Lawrence et al. 1995a, 2010). Recent phylogenomic studies have strongly favored the placement of Jacobsoniidae within Staphyliniformia (Zhang et al. 2018; McKenna et al. 2019), which is also supported by morphological characters such as similar hind wing structures and the maxillary galea of adults and larvae (Crowson 1959; 1960). The problematic Nosodendridae represent an isolated lineage sister to Staphyliniformia, Bostrichiformia, and Cucujiformia (Zhang et al. 2018; McKenna et al. 2019).

Because Derodontidae is consistently recovered with high support as sister to Clambidae and Eucinetidae (McKenna et al. 2015b, 2019; Zhang et al. 2018), we herein define the series Clambiformia *ser. nov*. and the superfamily Clamboidea *sensu nov*. to only include the families Clambidae, Derodontidae, and Eucinetidae. This new series is similar in composition to Crowson's (1981) Eucinetiformia, which included Scirtidae, Eucinetidae, and Clambidae. Morphological similarity between Derodontidae and the abovementioned families, mainly in terms of adult musculature, has been discussed by Kukalová-Peck and Lawrence (2004). Among the important characters uniting Derodontidae with Clambidae and Eucinetidae is the membranous connection of the pterothoracic ventrite. This character is only visible when the mesocoxae are removed from their cavities. It is present in Archostemata (Baehr 1975), but absent in Myxophaga while in Adephaga the two segments are secondarily joined by the interlocking of the meso- and metaventrites. Following Friedrich and Beutel (2006), we consider it to represent a putative shared plesiomorphy of Scirtiformia / Scirtoidea *sensu nov.* and Clambiformia *ser. nov*. / Clamboidea *sensu nov*. It was subsequently gained independently in some polyphagan groups, such as Leiodidae (Friedrich and Beutel 2006; Ge et al. 2007). The absence of *M. episterno-trochanteralis 2 (M. 70)* is a feature shared by Scirtoidea and Clamboidea *sensu nov*., in the sense presented here, although it has also been secondarily lost in some groups of polyphagan beetles such as Nosodendridae, and some groups in Scarabaeoidea, Pyrochroidae, and Dermestidae, while the absence of *M. episterno-coxalis 3 (M. 103)* can be considered a derived feature of the two groups (Friedrich and Beutel 2006; Ge et al. 2007). Following Ge et al. (2007), the shared loss of these three muscles, which are otherwise widespread in the remaining Polyphaga, can be considered another potential apomorphy of Scirtiformia / Scirtoidea *sensu nov.* and Clambiformia *ser. nov*. / Clamboidea *sensu nov.*

The remaining diagnostic features of Clambiformia *ser. nov*. are mainly plesiomorphic characters found also in Scirtiformia / Scirtoidea that exclude these taxa from the remaining polyphagan series. Nonetheless, it should be noted that even Scirtiformia / Scirtoidea *sensu* Lawrence et al. (1995b) was difficult to define and its diagnosis was likewise based in a large part on plesiomorphies (Lawrence and Newton 1995; Lawrence 2001). An externally exposed mesothoracic trochantin is also present in some Staphylinidae, Myxophaga, some Adephaga, and most Archostemata (Baehr 1975; Hansen 1997; Larsén 1966; Friedrich and Beutel 2006). Coxal plates are also present in Adephaga, Myxophaga, and some polyphagan groups such as Dascillidae and Ptiliidae (Hansen 1997; Beutel and Leschen 2005; Friedrich and

Beutel 2006). An accessory ventral process of the adult mandible is also present Dascillidae and Scarabaeoidea (Beutel et al. 2018).

The extinct family Mesocinetidae from the Upper Jurassic of Mongolia (Kirejtshuk and Ponomarenko 2010) shares similarities with eucinetids and may be their sister group (Lawrence 2019b). The family is therefore treated as a member of Clambiformia *ser. nov*., although this placement should be considered as only tentative.

### RHINORHIPIFORMIA Cai, Tihelka and Engel **ser. nov.**

***Type genus:*** *Rhinorhipus* Lawrence, 1988.

***Diagnosis:*** As for family Rhinorhipidae (Lawrence 1988, Kusy et al. 2018a): adult head hypognathous, with long temporal regions, lacking transverse occipital ridge or epicranial suture. Cranium with a short, median occipital endocarina and raised antennal insertions. Frontoclypeal region strongly declined and lacking a frontoclypeal suture. Clypeus long and narrow. Corpotentorium very broad. Oral cavity surrounded by setae. Labrum highly reduced, membranous. Mandibles long and with a setose dorsal cavity at base. Maxillae highly reduced, membranous, and setose. Pronotum without lateral carinae, anteriorly constricted and with a pair of elongate, vertical cavities. Procoxae slender and conical, trochantins completely visible, promesothoracic interlocking mechanism weakly developed. Mesothorax with a moderately developed mesoventral cavity reaching to the middle of the ventrite and a pair of well-developed procoxal housings on the mesanepisterna. Mesocoxae greatly projecting, mesotrochantin setose. Metaventrite with a moderately short discrimen. Metendosternite with long, curved lateral arms, anterior process with foramen at its base and a pair of expanded, ear-like, ventrolateral processes. Each elytron with 12 rather complete rows of deep punctures; epipleuron virtually absent, represented by slight thickening without lateral internal locking groove. Hind wing with r-m meeting stub of Rs; CuA3+4 meeting AA3, closing off anal (wedge) cell, cell acute; jugal lobe absent. Legs with enlarged and mesally produced metatrochanters, apically expanded metatibiae; tarsi simple, without pads, brushes, or membranous lobes; pretarsi with a well-developed empodium with two or three setae, and pectinate pretarsal claws (ungues). Abdominal tergite VIII of female dorsally concealed by tergite VII. Six Malphigian tubes. Larvae unknown.

***Included taxa:*** Rhinorhipidae Lawrence, 1988.

***Discussion:*** The enigmatic family Rhinorhipidae, represented in the modern fauna by a single species known only from northern New South Wales and southern Queensland (Lawrence 1988) is without a doubt one of the most mysterious beetle families. It possesses a suite of unusual characters that make resolving its systematic position difficult. Morphological analyses suggest an affinity to Elateriformia (Lawrence 1988; Lawrence et al. 1995b). As such, it has been included in Elateroidea (Bouchard et al. 2011; Lawrence et al. 2011) or placed as Elateriformia *incertae sedis* (Lawrence and Newton 1995). In a comprehensive morphological analysis, Lawrence (1988) recovered *Rhinorhipus* as sister to Elateroidea, while Lawrence et al. (1995b) recovered it as a sister to Byrrhoidea (all characters) or as sister to Buprestidae + Dascillidae (adult characters). A more recent extensive analysis of adult and larval morphological characters has recovered the family as sister to the elateriform family Rhipiceridae (Lawrence et al. 2011). Kusy et al. (2018a) recovered the family as the earliest diverging group within Elateroidea or as sister to Elateriformia and the rest of the derived Polyphaga. Consequently, the authors erected the superfamily Rhinorhipoidea to accommodate the group. In the more extensive analyses of McKenna et al. (2019) and our own analyses which account for long-branch artifacts, Rhinorhipidae are sister to Elateriformia and the remaining Polyphaga. Because it falls outside of Elateriformia, and would render the group paraphyletic, we erect the new series Rhinorhipiformia Cai, Engel and Tihelka *ser. nov*. for this unique family.

### BYRRHOIDEA Latreille, 1804 **sensu. nov.**

***Type genus:*** *Byrrhus* Linnaeus, 1767.

***Diagnosis:*** Adult body strongly sclerotized, highly convex and compact, often possessing mechanisms for compaction such as cavities for the reception of legs and fitting of hypognathous head against the anterior edge of the prosternum. Head strongly deflexed at base and deeply inserted into prosternum. Mandibles short and broad. Clypeus reduced. Base of labrum on lower plane than apex and usually forming a deep transverse impression between labrum and an abrupt frontal ridge. Eyes small and finely faceted, without interfacetal setae. Pro-mesothoracic interlocking device absent. Prosternum anteriorly not forming a produced chin-piece, but a head rest sometimes present. Procoxal and mesocoxal cavities widely separated. Transverse metaventral suture asbent. Epipleuron abruptly narrowed. Basal three (less often two) abdominal ventrites connate.

***Included taxa:*** Byrrhidae Latreille, 1804.

***Discussion:*** Byrrhids have been traditionally regarded as isolated groups within Elateriformia and treated as the sole members of Byrrhoidea (Crowson 1955, 1982; Beutel 2016). Lawrence (1988) subjected Elateriformia to the first cladistic analysis and expanded Byrrhoidea to include buprestids, byrrhids, and dryopoids. Lawrence and Britton (1991) subsequently resurrected Buprestoidea, which is supported by our results and retained in our classification system. The monophyly of Byrrhoidea as defined by Lawrence and Newton (1995) proved challenging to confirm based on adult, larval, and combined morphological datasets (Beutel, 1995; Lawrence et al., 1995, 2011; Costa et al., 1999). Early molecular studies based on a handful of genes have likewise yielded mixed results (Hunt et al., 2007; Bocak et al., 2014; McKenna et al., 2015). Analyses based on 89 (McKenna et al., 2019) and 68 genes (herein) have supported a clade unifying the former Buprestoidea and Byrrhoidea, together forming a sister group to a clade largely consistent with Dryopoidea of Crowson (1960, 1973, 1978, 1982) and Kasap and Crowson (1975). Hence, we herein resurrect the old concept of Byrrhoidea of Crowson (1955).

While different topologies supporting Buprestidae inside the old Byrrhoidea have been recovered by other recent morphological and molecular studies (e.g., Lawrence 1988; Lawrence et al. 2011; Bocakova et al. 2007; Bocak et al., 2014, Kundrata et al. 2014, 2017), it should be noted that the relationships of the 'byrrhoid-buprestoid complex' represent a lasting controversy in coleopteran systematics that should be addressed with more complete molecular and taxon sampling in future works.

### DRYOPOIDEA Billberg, 1820 (1817) **stat. res.**

***Type genus:*** *Dryops* A. G. Olivier, 1791.

***Diagnosis:*** Adult with frontoclypeal suture mostly distinct. Hind margin of pronotum crenulate or not. Procoxae projecting. Tarsomere 5 often as long as the basal 4 tarsomeres. Functional spiracles on 8th abdominal segment absent. Wing folding of "dryopoid" type. Most larvae with a well-developed and usually setose prostheca, short and wide prepharyngeal tube, broad and sclerotised gula, maxilla and labium closely connected, mandibular mola absent, labrum present, and a characteristic abdominal apex formed by tergite IX.

***Included taxa:*** Callirhipidae Emden, 1924; Chelonariidae Blanchard, 1845; Cneoglossidae Champion, 1897; Dryopidae Billberg, 1820 (1817); Elmidae Curtis, 1830; Eulichadidae Crowson, 1973; Heteroceridae MacLeay, 1825; Limnichidae Erichson, 1846; Lutrochidae Kasap and Crowson, 1975; Protelmidae Jeannel, 1950; Psephenidae Lacordaire, 1854; Ptilodactylidae Laporte, 1838.

***Discussion:*** Dryopoidea was defined by Crowson (1955, 1960, 1973, 1978, 1982) and Kasap and Crowson (1975) as a heterogeneous elateriform group that included beetles whose larvae are often aquatic and adults predominantly terrestrial. This early concept of the superfamily contained the families Chelonariidae, Dryopidae, Elmidae, Eulichadidae, Heteroceridae, Limnichidae, Lutrochidae, Psephenidae, and Ptilodactylidae. Lawrence and Newton (1982) added Callirhipidae which were earlier part of Artematopoidea or Rhipiceroidea, and Lawrence (1988) transferred here Cneoglossidae from Cantharoidea. Lawrence (1988) placed dryopoid beetles in a redefined broader concept of Byrrhoidea, which was adopted by subsequent authors (Lawrence and Newton 1995, Beutel 2016). Characters shared by most dryopoid larvae were discussed by Beutel (1995). On the molecular level, the superfamily is characterised by a unique rearrangement of tRNA gene order (Timmermans and Vogler 2012). Dryopoidea was monophyletic in Hunt et al. (2007), McKenna et al. (2015b; 2019), in the mitogenome analyses of Timmermans and Vogler (2012), and Timmermans et al. (2016), Zhang et al. (2018), and the present study.

NOSODENDRIFORMIA Cai and Tihelka **ser. nov.**

***Type genus:*** *Nosodendron* Latreille, 1804.

***Diagnosis:*** As for the family Nosodendridae (Beutel 1996; Ge et al. 2007; Leschen and Beutel 2010; Lawrence and Ślipiński 2013): Body oval and black, strongly convex dorsally, flattened ventrally. Head prognathous, inserted into the prothorax, with an enlarged mentum concealing the mouthparts. Frontoclypeal suture absent. Eyes large, almost meeting dorsally. Antennal insertions concealed from above by frontal projections. Antennae with a compact three-segmented apical club. All three segments of the thorax shorter than the maximum width of the metaventrite. Prothorax in front of procoxae short. Prosternal process long and wide, extending to mesoventrite. Cavities between hypomeron and prolegs present, further cavities for housing legs present on pterothorax and ventrites. Mesothoracic discriminal line absent. Mesocoxal cavities separated by a broad intercoxal process. Protrochantins concealed. *M. noto-trochantinalis* absent. Procoxae and procoxal cavities transverse. Metacoxal plates well-developed and complete. Metaventrite with more or less complete transverse suture. Radial cross-vein 3 absent, anal field strongly enlarged. Tarsal formula 5-5-5. Larva with labrum and clypeus fused, abdominal apex formed by posterior edge of segment VIII, and anal pads present.

***Included taxa:*** Nosodendridae Erichson, 1846.

***Discussion:*** The systematic placement of the problematic family Nosodendridae, the wounded-tree beetles, has always represented a persistent problem in systematic coleopterology. Earlier authors placed Nosodendridae into Byrrhoidea and Dascilloidea (Hayes and Chu 1946; Endrödy-Younga 1989), mainly due their gross similarity to byrrhids which also have an adult with a compact and convex body form. Crowson (1959) placed the family into the superfamily Dermestoidea. Lawrence and Newton (1995) considered the family to belong to Bostrichoidea, while Lawrence et al. (2010) placed the family into their Derodontiformia.

Ge et al. (2007) proposed several autapomorphies of Nosodendridae: the three segments of the thorax collectively shorter than the width of the metaventrite; accessory ridge laterally produced into a short, blunt tooth; presence of a hypomeral antennal groove; apical plate of the trochantinopleura distinctly enlarged; strongly shortened precoxal portion of the prosternum; wide prosternal process between procoxae; protrochantins concealed; transverse procoxae and procoxal cavities; mesothoracic discriminal line absent; broad intercoxal process separating mesocoxal cavities; anterior pronotal extension; dentiform process of the pronotal accessory ridge; the extensive apical plate of the trochantinopleura; radial cross-vein 3 absent; anal field strongly enlarged; *M. noto-trochantinalis* absent; and the beetle's strongly reduced flight musculature. The larvae of *Nosodendron* possess the following autapomorphies: labrum and clypeus fused; abdominal apex formed by posterior edge of segment VIII; and anal pads present (Beutel 1996).

STAPHYLINIFORMIA Lameere, 1900 **sensu nov.**

***Type genus:*** *Staphylinus* Linnaeus, 1758.

***Diagnosis:*** Antennae filiform or capitate, rarely serrate or pectinate. Mandibles more-or-less exposed, not concealed under labrum when abducted. Maxillary muscle *M. craniobasimaxillaris* present. Sternite II visible only as a lateral rudiment in nearly all representatives. Aedeagus never of cucujoid type. Metendosternite never of hylecoetoid type. Mesothoracic spiracles concealed under the hypomeron (most representatives). Radial cell primitively eyelet-like but weakened proximally by the obliteration of the base of RA 3+4. Radial bar with a single apical hinge, reinforced by the pinching together of RA 3+4 with the anterior wing margin. RA4 and RP 1 approaching one another and remaining parallel or fused together to form a Y-shaped support for the anterior apical wing margin and large apical field. Posterior coxae with oblique, non-excavate posterior face (absent in some staphylinoids). Protibia often strongly spinose or toothed outwardly. Tarsal formula usually 5-5-5 and simple. Malpighian tubules free, usually four present (Kukalová-Peck and Lawrence 1993; Hansen 1997; Lawrence and Ślipiński 2013).

***Included taxa:*** Histeroidea Gyllenhaal, 1808; Hydrophiloidea Latreille, 1802; Scarabaeoidea Latreille, 1802; Staphylinoidea Latreille, 1802.

***Discussion:*** Staphyliniformia, as presented here in its revised form, roughly corresponds to Kolbe's (1908) concept of Haplogastra, which included superfamilies united by having sternite II visible only as a lateral rudiment and pleural sclerite – Hydrophiloidea, Staphylinoidea, Histeroidea and Scarabaeoidea (Grebennikov and Scholtz 2004). The remaining Polyphaga (Kolbe's Heterogastra) has sternite II completely membranous. The Haplogastra concept was subsequently followed, with various modifications, by Peyerimhoff (1933), Jeannel and Paulian (1944), and Crowson (1955, 1960). The concept was later revived in analyses of wing characters (Kukalová-Peck and Lawrence 1993; referred to as the 'hydrophiloid lineage') and combined analysis of adult and larval morphological characters (Hansen 1997) and also appears to be well-supported by molecular data (Caterino et al. 2005; Song et al. 2010; McKenna et al. 2015b, 2019; Zhang et al. 2018).

The constriction of the hypopharynx between the maxillary bases is hourglass-shaped in cross sections in most staphyliniforms, as defined here, although exceptions are known in Silphidae (as Silphinae herein), some Leiodidae, and some Staphylinidae (Beutel and Leschen 2005). The early-diverging scarabaeoid genus *Glaresis* shares its paired ligular lobes, strongly sclerotised and setose anteriorly, with Hydrophiloidea (Anton and Beutel 2004; Beutel and Leschen 2005). In pupae, the presence of a maximum of only four functional abdominal spiracles has been suggested as a possible apomorphy of Staphyliniformia including Scarabaeoidea, or at least a large part of this group since some members such as some Scarabaeoidea, Hydrophilidae, Synteliidae, and Tachyporini have more (Lawrence and Newton 1982, Newton 2020). Anton and Beutel (2012) cited some potential apomorphies supporting Staphyliniformia *sensu nov*. They noted that scarabaeoids share with Hydrophiloidea antenna with asymmetrically widened club segments and discussed the possibility of a shared origin of the deep sensory pouch on the third club segment in Staphylinoidea and Scarabaeoidea. Staphyliniformia *sensu nov*. is also characterised by the shared presence of *M. craniobasimaxillaris*, an unusual maxillary muscle apparently found only in members of this group. Scarabaeoidea and some other members of the series also share the sclerotised labrum at least 2.5 times as broad as long, combined length of the scapus and pedicellus exceeding one third of the entire antennal length, and a cupuliform antennal segment VIII preceding three-segmented club. The same condition was suggested as an autapomorphy of Hydrophiloidea. Further characters typical of Staphyliniformia *sensu nov.* were listed by Lawrence and Ślipiński (2013).

### EROTYLOIDEA Latreille, 1802 **stat. nov.**

***Type genus:*** *Erotylus* Fabricius, 1775.

***Diagnosis:*** Adults with procoxal cavities internally open or closed by a slender bar. Glandular openings on the body, especially at the fore- and hind angles of the pronotum and prosternum. Tergite VIII in female dorsally concealed by tergite VII. Tarsal formula 5-5-5 in both sexes. Penis not divided into basipenis and distipenis, usually curved in lateral view, anterior edge usally with two penile struts. Adults phytophagous (Boganiidae, many Erotylidae), some pollinivorous, and mycophagous. Larvae with frontal arms lyriform (V-shaped in some Erotylidae), mesal surface of mandible with well-developed mola (some), maxillary articulating area present, hypopharyngeal sclerome present (missing in some Erotylidae). Some characters modified from Robertson et al. (2015).

***Included taxa:*** Boganiidae Sen Gupta and Crowson, 1966; Erotylidae Latreille, 1802.

***Discussion:*** The large and diverse Erotylidae was confirmed as monophyletic (Robertson et al. 2004; Wegrzynowicz 2002; Leschen 2003), as was the less diverse Gondwanan Boganiidae (Escalona et al. 2015). A close relationship between Erotylidae and Boganiidae was not generally suspected until these groups were recovered as sister taxa by Marvaldi et al. (2009) in an analysis of two rDNA fragments, albeit with considerably limited taxon sampling. In a four-gene analysis, Bocak et al. (2014) recovered Boganiidae as nested within Erotylidae. Genomic analyses have, however, unequivocally recovered Boganiidae and Erotylidae as a clade sister to the remaining Cucujoidea *sensu lato* (Zhang et al. 2018; McKenna et al. 2019; present study).

Boganiidae was not been sampled in many past analyses of Cucujoidea and so the set of characters supporting the superfamily is not extensive. Some Boganiidae (Escalona et al. 2015) and Erotylidae have glandular ducts (Crowson 1990, 1991; McHugh et al. 1997; Leschen 2003), some of which penetrate deep within the body. Similar ducts are present in several other families that are now placed in Coccinelloidea (Discolomatidae) and Cucujoidea (Cryptophagidae), and they also bear resemblance to ducts present in Chrysomeloidea and elsewhere (e.g, Drilling et al. 2013). Tightly bundled glandular ducts, referred to as a microtubulate condition by Leschen (2003), are present in Boganiinae (Boganiidae) and Pharaxonothinae (Crowson 1990, 1991; Leschen 2003). The penis, which is not divided into two parts that characterise several members of the Cucujoidea (e.g., Leschen et al. 2005; Lawrence et al. 2011) is of a single piece and often curved in lateral view, resting in the abdomen on its side. The anterior edge usually bears two elongate struts, which are short in Boganiinae, but elongate in most remaining taxa and fused into a single process. The extent to which aedeagal characters could be considered potential synapomorphies requires additional study.

Both Boganiidae and some phytophagous Erotylidae also share some ecological similarities, even though the ancestral state, at least of Erotylidae, may be fungus feeding. Both contain species that visit cycads in high numbers and probably contribute to their pollination (Crowson 1991; Leschen and Buckley 2007; Suinyuy et al. 2009; Prado et al. 2011). Given the discovery of Cretaceous boganiids preserved alongside cycad pollen, it is likely that the association between Erotyloidea *stat. nov*. and cycads is over 100 Ma old (Cai et al. 2018b).

Some of the families traditionally included in the 'basal Cucujoidea' that lack genetic data may eventually be placed into Erotyloidea; but at present there is no compelling morphological evidence to place them here.

Similarities between the monogeneric *Lamingtonium* and Boganiidae + Erotylidae were discussed by Sen Gupta and Crowson et al. (1969), and the external habitus and fungal feeding behaviour of the group are suggestive. However, the true affinities of the family are difficult to demonstrate conclusively (Lawrence et al. 2011), they lack the glandular ducts present in the families above, and the genitalia differ (Lawrence and Leschen 2003). Lawrence and Leschen (2003) discussed the possible relationship between Erotylidae, Lamingtoniidae, and Cyclaxyridae based on larval characters, and the sister relationship of Tasomosalpingidae and Cyclaxyridae was shown by Leschen et al. (2005). Cyclaxyridae lack glandular ducts and the Tasmosalpingidae have trichome-like structure at the posterior margin of the prothorax that interrupts the pronotal carina but doesn't appear to have the glandular ducts similar to Erotylidae or Boganiinae. Among the aforementioned families, the bipartite penis may not be present, and similar penile struts present in most erotylids also occur in Cyclaxyridae and Tasmosalpingidae, but the former has been placed among Cucujoidea by McKenna et al. (2015b) and Robertson et al. (2015) that placed it as sister taxon to Myraboliidae.

### NITIDULOIDEA Latreille, 1802 **stat. nov.**

***Type genus:*** *Nitidula* Fabricius, 1775.

***Diagnosis:*** Adults rarely with elongate rostrum. Dorsal part of mandible without setose cavity (except Sphindidae). Subantennal grooves well-developed (except Nitidulidae: Calonecrinae, Smicripidae). Antennae never fully filiform or moniliform, and a club is always present. Gular sutures never confluent (except some Nitidulidae, some Monotomidae). Ventral or lateral portion of prothorax on each side never with incomplete notosternal suture or without sutures (except some Nitidulidae, some Sphindidae, some Monotomidae). Cuticular glands, where present, consist of single ducts. Procoxal cavities internally open (most). Tarsal formula 5-5-5 in female and 5-5-5 or 5-5-4 in male (rarely 4-4-4). Ventrite 1 never much longer than ventrite 2 (except some Nitidulidae and Monotomidae). Abdominal tergite VII exposed in dorsal view and tergite VIII in male with sides curved ventrally forming a genital capsule (in Kateretidae, Monotomidae, Nitidulidae, and Smicripidae). Tergite VIII in female dorsally concealed by tergite VII, tergite X in male completely membranous. Spiculum gastrale present (absent in Sphindidae and some Monotomidae). Larvae with frontal arms lyriform (most), mesal surface of mandible with well-developed mola (most), maxillary articulating area present (most), hypopharyngeal sclerome present (most).

***Included taxa:*** Helotidae Chapuis, 1876; Kateretidae Kirby, 1837; Monotomidae Laporte, 1840; Nitidulidae Latreille, 1802; Protocucujidae Crowson, 1954; Smicripidae Horn, 1880; Sphindidae Jacquelin du Val, 1860.

***Discussion:*** The former superfamily Cucujoidea *sensu* Crowson (1955) and Lawrence and Newton (1982), more or less corresponding to the old Clavicornia, is a very heterogeneous group containing mostly minute to moderately sized brown to dark beetles that are difficult to place elsewhere (Leschen et al. 2005). As such, the old Cucujoidea has been considered to represent a taxonomically complicated 'wastebasket group' that may or may not be monophyletic (Leschen and Ślipiński 2010). There have historically been ample proposals to divide the group, which were reviewed in detail by Lawrence et al. (1995a). Robertson et al. (2015) provided the first step to splitting the traditional Cucujoidea with the use of molecular data, erecting the superfamily Coccinelloidea for ladybird beetles, handsome fungus beetles, and their relatives, and removing the Biphyllidae and Byturidae to Cleroidea. No other divisions of Cucujoidea have been formalised since then. It is notable that in the latter part of his career, Crowson considered the establishment of a separate superfamily for sap beetles and their allies, Nitiduloidea (Audiso 1993; Crowson 1995), although this was never proposed formally. Nonetheless, the group of families surrounding Nitidulidae has been consistently recovered in morphology-based phylogenies and has often been referred to as the 'nitidulid lineage', 'nitidulid series' or 'nitidulid assemblage' (Ślipiński 1998; Leschen et al. 2005; Cline et al. 2014; McKenna et al. 2015b). Genome-scale analyses have unambiguously established the existence of a clade containing the 'nitidulid lineage' and the more basal families Helotidae, Sphindidae, Protocucujidae, and Monotomidae (McKenna et al. 2019).

The monophylum consisting of Monotomidae, Nitidulidae, Kateretidae, and Smicripidae is supported by abdominal tergite VII exposed in dorsal view and tergite VIII in males with sides curved ventrally forming a genital capsule (Leschen et al. 2005). LeConte (1861) treated this group as a unified Nitidulidae. Crowson (1955) also considered Monotomidae to belong to the nitidulid group, although he later revised this idea (unpublished, cited in Leschen et al. 2005). The clade containing only Nitidulidae, Kateretidae, and Smicripidae is strongly supported by the frontoclypeal suture present (most), anterior cervical sclerite not contiguous with head capsule or placed in paired emarginations, galea absent (most), anterior pronotal angles absent or not produced forwards, anterior tendons of metendosternite narrowly separated, abdominal tergite VII exposed in dorsal view, tergite VIII in males with sides curved ventrally forming a genital capsule, paired endocarinae absent in the ventral head capsule of larvae, four or fewer stemmata, apex of larval mandible bidentate (most), mesal surface of larval mandible with asperate or tuberculate mola, labial palpi 1-segmented in larva, and larval hypopharyngeal sclerome an irregular molar-like tooth (most) (Leschen et al. 2005). Of these, the absence of a galea is apparently autapomorphic for the group. The basal clade Protocucujidae + Sphindidae has traditionally been well-accepted (Sen Gupta and Crowson 1979; McHugh 1993; Ślipiński 1998) and is supported by the following shared morphological characters: frontoclypeal suture present, apex of mandible tridentate, anterior portion of prosternum at midline as long as prosternal process, procoxal cavity internally open, metendosternal laminae well developed, 5 free veins present in medial area of the hind wing, 5-5-4 male tarsal formula, aedeagus with U-shaped sclerite near apex, loss of articulated parameres, internal sac with basal complex sclerite, labrum of larvae vertical, apex of larval mandible tridentate, and ventral epicranial ridges of larval head capsule absent (Thomas 1984a; Ślipiński 1998; Beutel and Ślipiński 2001; Leschen et al. 2005). Similarities between the two taxa are especially evident in the South American genus *Protosphindus* Sen Gupta and Crowson (1979).

Characters shared by nitidulid families are listed above in the diagnosis section. Identifying characters that support the whole of the eco-morphologically remarkably diverse Nitiduloidea, as defined here, is more difficult. This is not surprising, however, because the same was the case for the old Cucujoidea *sensu* Crowson (1955) and Lawrence and Newton (1982) which likewise lacked a set of unique and widespread apomorphies (Kirejtshuk 2000; Leschen and Ślipiński 2010). Nonetheless, commonalities of the whole Nitiduloidea *stat. nov*. have been noted by coleopterists for over a century. Sharp and Muir (1912) noted similarities of the aedeagus of Helotidae, Monotomidae, and Nitidulidae, although they noted that their taxon sampling was not wide enough to draw definitive conclusions.

Because of the inherent difficulties associated with studying the phylogeny of cucujoid beetles, the division of Cucujoidea *sensu* Robertson et al. (2015) proposed by us into three separate superfamilies leaves several enigmatic families of uncertain placement that were not included in our analyses because of the unavailability of sequences and lack of consensus in morphology-based phylogenies. In the morphological phylogenetic analysis of Leschen et al. (2005), Tasmosalpingidae formed a monophyletic group with Cyclaxyridae but was simultaneously also close to Nitiduloidea based on all characters or adult characters only. When larval characters were analysed alone, it was recovered as sister to Phalacridae. The placement of the fossil families †Parandrexidae and †Wabbelidae has not been formally tested (Kirejtshuk 1994; Escalona et al. 2015; Alekseev, 2017). These fossil taxa are treated as Cucujiformia incertae sedis, as they cannot be placed for sure in either Erotyloidea stat. nov., Nitiduloidea stat. nov., or Cucujoidea sensu nov. with certainty.

### CUCUJOIDEA Latreille, 1802 **sensu nov.**

***Type genus:*** *Cucujus* Fabricius, 1775.

***Diagnosis:*** Adults typically with dorsoventrally flattened bodies (except Cyclaxyridae, some Passandridae, Phalacridae). Head sometimes with elongate rostrum, longer than wide (some Silvanidae, some Laemophloeidae). Dorsal part of mandible without a setose cavity (present in some Silvanidae, some Phloeostichidae). Subantennal grooves weakly developed or absent (except some Passandridae, some Laemophloeidae, Myraboliidae). Antennae filiform or moniliform (some Silvanidae, Cucujidae, some Passandridae, some Laemophloeidae). Tentorium with a medial process (except Myraboliidae, Phalacridae, and Laemophloeidae). Gular sutures at least partly confluent (only in Passandridae). Ventral or lateral portion of prothorax on each side with incomplete notosternal suture or without sutures (some Cryptophagidae, some Silvanidae, some Phloeostichidae, Laemophloeidae). Cuticular glands, where present, consist of single ducts. Procoxal cavities internally open (except some Passandridae, Phalacridae). Ventrite 1 sometimes much longer than ventrite 2 (some Cryptophagidae, some Silvanidae, Myraboliidae). Tergite VIII in female dorsally concealed by tergite VII, tergite X in male completely membranous. Unequal anterior tibial spurs (some, in Passandridae, Phalacridae and Laemophloeidae). Tarsal formula 5-5-5 in female and 5-5-5 or 5-5-4 in male (rarely 4-4-4 or 4-5-4). Penis may be clearly divided into basipenis and distipenis (Leschen, pers. obs.) or undivided with a single, often broad penile strut. Larvae with frontal arms lyriform (most), mesal surface of mandible with well-developed mola (most), maxillary articulating area present (most), hypopharyngeal sclerome present (most).

***Included taxa:*** Agapythidae Sen Gupta and Crowson, 1969; Cavognathidae Sen Gupta and Crowson, 1966; Cryptophagidae Kirby, 1826; Cucujidae Latreille, 1802; Cyclaxyridae Gimmel, Leschen and Ślipiński, 2009; Hobartiidae Sen Gupta and Crowson, 1966; Laemophloeidae Ganglbauer, 1899; Lamingtoniidae Sen Gupta and Crowson, 1969; Myraboliidae Lawrence and Britton, 1991; Passandridae Blanchard, 1845; Phalacridae Leach, 1815; Phloeostichidae Reitter, 1911; Priasilphidae Crowson, 1973; Silvanidae Kirby, 1837; Tasmosalpingidae Lawrence and Britton, 1991.

***Discussion:*** Cucujoidea, in the strict sense as conceived here, corresponds to the most derived clade of the old Cucujoidea *sensu* Crowson (1955) and Lawrence and Newton (1982) and the 'basal Cucujoidea' of Leschen et al. (2005) recovered in our and previous genome-scale molecular studies (Zhang et al. 2018; McKenna et al. 2019). As with the former Cucujoidea *sensu* Crowson (1955) and Lawrence and Newton (1982) or Cucujoidea *sensu* Robertson et al. (2015), finding definitive apomorphies of the group is difficult. The definitions of many of the cucujoid families discussed here have been considerably fluid in the past, often lacking definitive apomorphies, but most concern groups broadly defined in recent decades as Cucujidae and Phleoeostichidae. Nonetheless, Cucujoidea *sensu nov.* contains several clades that were strongly suspected to be monophyletic based on morphological studies, in some cases long before the advent of molecular systematics. These include Laemophloeidae, Phalacridae, and Passandridae (Thomas 1984b); Leschen's (1996) 'cucujid line' consisting of Cucujidae, Hobartiidae, Phloeostichidae, Cryptophagidae, Cavognathidae, and Silvanidae, following Thomas (1984a); and the 'basal Cucujoidea' of Leschen et al. (2005). Cucujoidea *sensu nov*. is also mostly congruent with the 'cucujid series' as recovered in molecular studies by Hunt et al. (2007), Bocak et al. (2014), McElrath et al.

(2015), and Robertson et al. (2015).

The relationships among the family groups and potential synapomorphies for Cucujoidea have been discussed in several papers, but none of the characters are conclusive synapomorphies, as discussed in Leschen (1996), relevant chapters in Leschen et al. (2010), and the study by McElrath et al. (2015) where some salient features of the adults were reconstructed over the Cucujoidea tree. Some characters like the presence or absence of the tentorial spine in the adult or the number of tarsungular setae in the larva (Leschen 1996) are too variable within the group to be identified as a synapomorphy (Leschen 1996). One feature relevant for defining Cucujoidea may be the presence of a bipartite or subdivided penis (Leschen, unpubl.). The penis in Cucujiformia is highly diverse (Lawrence and Slipinski 2013) and many groups have a single or double penile strut. Several members of the Cucujoidea as defined here have a bipartite penis (Cavognathidae, Priasilphidae, Cucujidae, Agapythidae, and Silvanidae; Leschen et al. 2005; Lawrence et al. 2011) and the anterior part could be homologous to the penile strut. Examination of dissections of Hobartiidae and Passandridae and images of Myraboliidae indicate that these may also have a subdivided penis, whereas the remaining families clearly have an undivided penis (Cyclaxyridae, Lamingtoniidae, Phloeostichidae, Phalacridae and Tasmosalpingidae). If we examine this variation in light of our tree and those of Zhang et al. (2018) and McKenna et al. (2019), a bipartite aedeagus is optimised as a synapomorphy, the undivided type (in different forms) as derived. These aedeagal types are similarly optimised on the trees in McElrath et al. (2015) and Robertson et al. (2015).

### †Alphacoleoptera Engel, Cai, and Tihelka **subord. nov.**

***Type family:*** †Tshekardocoleidae Rohdendorf, 1944.

***Diagnosis:*** Procoxal cavities closed posteriorly; mesothorax shortened and metathorax enlarged; forewings complete, flat over abdomen at reset, with wide epipleura, meeting along and putatively locking along sutural margin except apices acute and typically not meeting along sutural margin (thus, largely but not completely covering metasoma), venation complete, reticulate, longitudinal sectors typically forking (sometimes zigzagged), with multiple cells between veins, small field proximally between Sc and RA (except Permosynoidea), anal vein following sutural margin (posterior trailing margin opposite leading edge), with shoulder placed proximally on A1+2; mesoventrite longer than prosternum; tarsal formula 5-5-5; telescopic ovipositor scarcely sclerotized.

***Included taxa:*** Superfamily †Permocupedoidea Martynov, 1932, Superfamily †Permosynoidea Tillyard, 1924, and Superfamily †Coleopseoidea Kirejtshuk and Nel, 2016.

***Etymology:*** The name of this suborder is a combination of Greek *álpha* (*ἄλφα*), first letter of the Ancient Greek alphabet and an allusion to "first", and Coleoptera, itself a combination of Greek *koleós* (*κολεός*), meaning, "sword-sheath", and *pterá* (*πτερά*, plural of pterón, *πτερόν*), meaning, "wings".

***Discussion:*** This group has often been referred to as †Protocoleoptera Tillyard, 1924. Unfortunately, Protocoleoptera was established explicitly for the extinct family †Protocoleidae Tillyard, 1924, a group today known to be a lineage of polyneopteran insects allied to the earwigs (Dermaptera). Another extinct famly, †Permophilidae Tillyard, 1924, from the same fossil beds was later established as †Paracoleoptera Laurentiaux, 1953, and also considered a lineage allied to the beetles. †Permophilidae, like †Protocoleidae, are polyneopterans (*e.g.*, Forbes, 1928; Kukalová, 1966). Accordingly, †Protocoleoptera and †Paracoleoptera are objective synonyms of †Protelytroptera, are unrelated to Coleoptera, and, in fact, do not belong to the Eumetabola. Neither †Protocoleoptera nor †Paracoleoptera can be used as suborders of Coleoptera. Similarly, Crowson (1981) established an extinct suborder, †Archecoleoptera, which he considered to exclude the most primitive of beetles but be confined to the Permian. Crowson still considered †Protocoleoptera to be a lineage of early beetles including the †Tshekardocoleidae and others, and therefore excluded them from his ill-defined †Archecoleoptera. †Archecoleoptera were not established on the basis of an specific taxon or characters, but was meant to exclude at the least †Tshekardocoleidae and other early extinct beetles. Thus, the basalmost and extinct suborder of beetles remains without an appropriate name and one is therefore established here.

### Zacoleoptera Engel, Cai, and Tihelka **capaxord. nov.**

***Diagnosis:*** Procoxal cavities distinct and open; narrow prosternal process present.
***Included taxa:*** Suborder Metaxycoleoptera nov. and Hyporder Eucoleoptera nov.
***Etymology:*** The name of this capaxorder is a combination of Aeolian Greek *zá* (*ζά*), meaning, "very", and Coleoptera.

### †METAXYCOLEOPTERA Engel, Cai, and Tihelka **subord. nov.**

***Type family:*** †Ponomarenkiidae Yan et al., 2017.
***Diagnosis:*** Fossil beetles with: antenna moniliform; elytra without window punctures; propleuron exposed; absence of coxal plates; absence of broad contact between meso- and metaventrites; procoxal cavities distinct and open; narrow prosternal process present; internalized metatrochantin; metanepisternum only marginally part of closure of mesocoxal cavity; transverse ridge of mesoventrite; short metacoxae not reaching hind margin of abdominal sternite III. They share with Eucoleoptera open procoxal cavities and a narrow prosternal process.
***Included taxa:*** Family †Pronomarenkiidae Yan et al., 2017.
***Etymology:*** The name of this suborder is a combination of Greek *metaxú* (*μετᾰξῠ*), meaning, "between", and Coleoptera.
***Discussion:*** This extinct group is likely sister to the Eucoleoptera, *i.e.*, the clade comprising all extant beetle suborders.

### EUCOLEOPTERA Engel, Cai, and Tihelka **hypord. nov.**

***Type family:*** Carabidae Latreille, 1802.
***Diagnosis:*** Preëpisternal cervical sclerites lacking; procoxal cavities present and distinctly countersunk (secondarily modified in several clades); broad prothoracic postcoxal bridge lacking; narrow prosternal process; mesothoracic discrimen absent (secondarily reversed in various clades); mesothoracic meron absent; mesoscutellar-elytron locking device present; hind wing triangular (pivitol) fold proximal to RA3+4 (*i.e.*, not dividing vein), with vein typically incompletely proximally; mesocoxal cavities deep and completely demarcated mesally and posteriorly by distinct ridge (secondarily reduced numerous times); metascutal fissure present; alacristae present; metapostnotum undivided; metathoracic meron absent; metacoxae transverse, with metacoxal cavities distinct, completely demarcated posteriorly by carina; reduced costal crossvenation; sternite I reduced; female tergite IX divided into paraprocts on either side of tergite X.
***Included taxa:*** Adephaga Clairville, 1806, Archostemata Kolbe, 1908, Myxophaga Crowson, 1955, and Polyphaga Emery, 1886.
***Etymology:*** The name of this hyporder is a combination of Greek *eû* (*εὖ*), meaning, "good" or "true", and Coleoptera.
***Discussion:*** This group comprises all extant suborders of Coleoptera, distinct from the exclusively Permian alphacoleopterans.

### Notes on the classification of beetles

†Magnocoleidae Hong, 1998 *syn. nov*. – The monotypic fossil family †Magnocoleidae was described by the preeminent Chinese palaeoentomologist Professor Youchong Hong (1998) based on isolated elytra from the Lower Cretaceous Qingshila Formation of China, which were named as *Magnocoleus huangjiapuensis*. No further species belonging to the family have been described since, probably owing to the poorly accessible original description published in Chinese with only a brief English summary in a regional geological bulletin. Hong compared the new family with Permian, Triassic, and Jurassic protocoleopterans and archostematans, and differentiated it based on elytral venation characters. However, it should be noted that the elytral wing venation pattern of *Magnocoleus* falls well within the range of morphological variation observed in Mesozoic Cupedidae and Ommatidae. Using Ponomarenko's (1969) key to Archostemata *sensu lato*, the presence of five principal elytral veins, together with the shortened ones, and the complete reduction of CuP places *Magnocoleus* into Cupedidae or Ommatidae. Indeed, in a phylogenetic analyses of elytral characters *Magnocoleus* was recovered as sister to Ommatidae and Cupedidae (Lubkin 2007). We thus consider †Magnocoleidae as a junior synonym of Cupedoidea and *Magnocoleus* as *incertae sedis* within that superfamily. The lack of complete body fossils makes it difficult to determine whether the beetles belonged to Ommatidae or Cupedidae, as the two families cannot be reliably distinguished on the basis of elytral venation alone (Lawrence 1999).

Heteroceridae MacLeay, 1825. – The variegated mud-loving beetles were recovered as nested within Limnichidae with moderate support. Heterocerids are characterised by a set of morphological characters that represent extreme adaptations for life in near-water environments, such as fossorial legs and body covered with dense water-repellent setae (Vanin et al. 2016). Historically, the monophyly of Limnichidae, the minute marsh-loving beetles, has been doubted on the basis of both morphological and molecular data (Hinton 1939; Crowson 1978; Beutel 1995; Costa et al. 1999; Bocak et al. 2014). Crowson (1978) regarded Heteroceridae as a sister group to Limnichidae. In a morphological phylogenetic analysis, Beutel (1995) recovered Limnichidae as the sister group to Heteroceridae based on larval morphological characters, while Costa et al. (1999) placed Heteroceridae in a clade with Dryopidae + paraphyletic Limnichidae. Heteroceridae was placed within a paraphyletic Limnichidae in Kundrata et al.'s (2017) analysis of four DNA fragments and the phylogenomic studies of Zhang et al. (2018) and an 89 gene dataset in McKenna et al. (2019). In our analyses, *Heterocerus* was recovered as sister to Limnichinae with full support, although the subfamilies Hyphalinae and Thaumastodinae were not sampled. In the analysis of McKenna et al. (2019), *Heterocerus* was sister to Cephalobyrrhinae.

Regardless of the precise sister group of Heteroceridae it shares with apparently all Limnichidae the presence of a frontoclypeal suture on the adult head, absence of the radial cell in the hindwing, asymmetrical base of the basal piece of the aedeagus, as well as the following larval synapomorphies: head with dense vestiture of long and short setae, presence of accessory anterior tentorial bridge, and a short antenna with a large hyaline sensorium (Crowson 1978; Beutel 1995; Costa et al. 1999). Crowson (1978) furthermore pointed out that the larvae in both families share mandibles with a plumose prostheca and retractable anal tracheal gills in at least some taxa.

We treat Heteroceridae as a separate family, although its incorporation within Limnichidae should be considered if future studies arrive at identical conclusions with increased gene sampling and higher statistical support.

Jacobsoniidae Heller, 1926. – The family Jacobsoniidae is herein treated as a member of the superfamily Staphylinoidea. Jacobsoniidae have been placed in a variety of basal polyphagan superfamilies in the past, namely in Bostrichiformia (Lawrence and Newton 1995), Cucujiformia (Sen Gupta 1979) and Derodontiformia (Lawrence and Leschen 2010). Recently, Lawrence (2016) removed Jacobsoniidae from Derodontiformia and treated it as series and superfamily *incertae sedis*.

The placement of some or all of members of Jacobsoniidae within Staphylinoidea is supported by morphological and molecular studies (Lawrence et al. 2011; McKenna et al. 2015b, 2019; Toussaint et al. 2017; Zhang et al. 2018). The three very different-looking genera have not been included together in any recent analyses, and only two (*Derolathrus* and *Saphophagus*) have genetic data available. Ptiliidae, which together with Hydraenidae formed the sister group to Jacobsoniidae in our analyses, shares with the family reduced hindwing venation and very long fringe hairs forming a "feather-wing", with a complex folding pattern aided by spicules on the abdominal tergites (Crowson 1960; Yamamoto et al. 2017). The jacobsoniid larval galea also suggests a relationship with Staphylinoidea (Crowson 1960). It should nonetheless be noted that our analyses are only based on the jacobsoniid genus *Derolathrus*; however, *Saphophagus* was placed in the same position in McKenna et al. (2019). Due to the considerable morphological differences between the three genera, it is not certain whether Jacobsoniidae as presently defined is in fact monophyletic (Löbl and Burckhardt 1988), and genetic data for *Sarothrias* (senior synonym of the type genus of the family) would test this hypothesis and settle the superfamily placement of the family.

COLONIDAE Horn, 1880 (1859) *stat. nov*. – The leiodid subfamily Coloninae is herein treated as a separate family of Staphylinoidea, in agreement with some historical treatments of Leiodidae and related families (e.g., Jeannel 1936; Perkovsky 1997). In addition to having a brick-wall pattern of abdominal intersegmental membranes, unique outside of Staphylinidae sensu nov. as noted below under Silphinae, colonines lack the conspicuous leiodid character of an "interrupted club" with smaller antennomere 8 (e.g., Perkovsky 1997), and have many other characters that are unique or unusual within Leiodidae, including cervical sclerites absent, procoxal cavities closed by fusion of the pronotal and prosternal processes, abdominal spiracles 7-8 absent, and others noted by Newton (1998). Recent molecular phylogenetic studies have also placed Coloninae as an isolated staphylinoid group well outside of an otherwise monophyletic Leiodidae, either as a sister to Hydraenidae+Ptiliidae (McKenna et al. 2015a), or sister to Agyrtidae+Leiodidae (minus Coloninae) (Lü et al. 2020). Recent discovery of mid-Cretaceous fossils (Cai and Huang 2017; Yamamoto and Takahashi 2018) further supports this separation by showing that an unreduced antennomere 8 was already present then and not likely a later secondary development within Leiodidae as previously proposed (e.g., Hansen 1997).

SILPHINAE Latreille, 1806 *stat. nov.* – The family Silphidae *sensu* Sikes (2016) is herein treated as a subfamily of Staphylinidae *sensu nov*. Difficulties in clearly distinguishing the two groups were pointed out by coleopterists for almost a hundred years; Hatch (1927) treated the burying beetles as a subfamily of Staphylinidae, but his proposal was not widely accepted. A number of morphological and molecular studies conducted since have firmly established Silphinae *stat. nov.* as either nested within Staphylinidae or at most a sister group to that family, together forming a monophyletic clade (Hansen 1997, Beutel and Molenda 1997; Beutel and Leschen 2005; Caterino et al. 2005; Hunt et al. 2007; Grebennikov and Newton 2009, 2012; Lawrence et al. 2011; McKenna et al. 2015a, 2015b, 2019; Zhang et al. 2018; Lü et al. 2020; Song, et al. 2021). The sister group of Silphinae *stat. nov*. is not resolved, as past studies have suggested a range of topologies often placing the subfamily as sister to the remaining staphylinids (e.g., Grebennikov and Newton 2012) or associated with the subfamily Phloeocharinae (Caterino et al. 2005) or Tachyporinae (McKenna et al. 2015a, 2015b; Lü et al. 2020; Song et al. 2021; Yamamoto 2021). In our analyses, the burying beetles were recovered as sister to a clade containing Osoriinae, Apateticinae, and Scaphidiinae. This rather derived placement within Staphylinidae is consistent with the hypothesis of Lawrence and Newton (1982).

The inclusion of Silphinae *stat. nov*. within Staphylinidae *sensu nov.* is supported by a typical brick-wall pattern of minute sclerites in the abdominal intersegmental membranes, which may represent a synapomorphy of the two groups, although it also occurs in the enigmatic leiodid genus *Colon* Herbst (Hansen 1997) and is not present in many groups (Thayer 2016). The two taxa share a number of morphological characters, most likely correlated with reduced elytra that are abbreviated apically and associated with a more flexible abdomen: a unique wing hinge and asymmetrical folding pattern, aedeagus usually everted asymmetrically and with a large basal bulb and internal muscles for evaginating the internal sac, larvae without labral tormae and mandibular molae but with a modified tentorium whose posterior arms attach directly to the ventral surface of the head and whose bridge is directed posteriorly (Lawrence and Newton 1982; Newton 1997; Sikes et al. 2016). We consider this evidence, together with the results of molecular phylogenetic studies, as sufficient for formally treating the carrion beetles as members of Staphylinidae *sensu nov*.

The former subfamilies of Silphinae *stat. nov*. are downgraded to tribal level: Silphini Latreille, 1806 and Nicrophorini Kirby, 1837 (both *stat. nov*.).

CLEROIDEA Latreille, 1802. – Gimmel et al. (2019) recently conducted the first in-depth molecular analysis of the relationships within the superfamily Cleroidea, sampling four genes across a wide range of taxa. While the taxon sampling in our study was insufficient to test all their newly proposed classification changes in the superfamily, we did recover their newly proposed Thymalidae, represented by *Thymalus*, nested within Trogossitidae *sensu stricto* of Gimmel et al. (2019). While *Thymalus* was historically placed within the trogossitid subfamily Peltinae (now Peltidae), it does not fit the diagnosis of Trogossitidae in the new, restricted sense, which includes only taxa with broadly closed procoxal cavities (Gimmel et al. 2019). We defer this issue to a subsequent study that includes more taxa of the Trogossitidae (*sensu stricto*) and the other subfamily of Thymalidae, the Decamerinae.

CYBOCEPHALINAE Jacquelin du Val, 1858 *stat. nov*. – The family Cybocephalidae is herein treated as subfamily Cybocephalinae stat. nov. within Nitidulidae. The rank and systematic position of this peculiar group was recently discussed by Kirejtshuk and Mantič (2015) and Lawrence (2019c).

MELOIDAE Gyllenhal, 1810. – The family Meloidae *sensu* Bologna et al. (2010) was recovered nested within Anthicidae. Due to their unusual life habits and a similarly unusual morphology, the systematic position of blister beetles has long been considered unresolved. A relationship with the family Anthicidae has been hypothesised by a number of coleopterists in the past (Crowson 1955; Abdullah 1965a, b, 1969; Selander 1966, 1991; Bologna and Pinto 2001). The subtribe Protomeloina has been regarded by Abdullah (1965a, b) as a link between the family and the remaining Anthicidae. The same author also proposed that blister beetles are in fact derived anthicids (Abdullah 1969). Meloidae have been recovered as nested within Anthicidae in the phylogenomic analyses of Zhang et al. (2018) and McKenna et al. (2019) as well as in some analyses of fewer genes (Gunter et al. 2014). Both families share a strongly deflexed head, a somewhat inflated vertex, a narrow neck, and a pronotum lacking lateral carinae (Bologna et al. 2010). Indeed, some meloids and anthicids can be easily confused and unsurprisingly there is a long history of taxa transferred between the two groups. It is interesting to note that some anthicid beetles are attracted to blister beetles and feed on the cantharidin they secrete, which is possibly used for predator defense and/or as a mating attractant, leading Abdullah (1964) to postulate that the production of this substance may have been ancestral to the two groups. This hypothesis seems supported by the fact that some male anthicids have been recorded to secrete cantharidin (Schütz and Dettner 1992). At present, we regard Meloidae as a separate family, but note that the accumulation of further molecular evidence may result into the group being joined within Anthicidae. An alternative treatment may be to divide Anthicidae, which currently contains several sufamilies with apparently disparate morphology, depending on future morphological and molecular evidence.

### Classification of beetles (families and subfamilies)

#### Order COLEOPTERA Linnaeus, 1758

†Suborder **ALPHACOLEOPTERA** Engel, Cai, and Tihelka, *nov*.

†Superfamily **Coleopseoidea** Kirejtshuk and Nel, 2016
†Family Coleopseidae Kirejtshuk and Nel, 2016

†Superfamily **Permocupedoidea** Martynov, 1932
†Family Tshekardocoleidae Rohdendorf, 1944
†Family Labradorocoleidae Ponomarenko, 1969
†Family Oborocoleidae Kukalová, 1969
†Family Permocupedidae Martynov, 1932
†Subfamily Permocupedinae Martynov, 1932
†Subfamily Taldycupedinae Rohdendorf, 1961

†Superfamily **Permosynoidea** Tillyard, 1924
†Family Ademosynidae Ponomarenko, 1968
†Family Permosynidae Tillyard, 1924

Capaxorder **ZACOLEOPTERA** Engel, Cai, and Tihelka, *nov*.

Suborder Metaxycoleoptera Engel, Cai, and Tihelka, nov.
†Family Ponomarenkiidae Yan, Lawrence, Beattie and Beutel, 2017

Hyporder **EUCOLEOPTERA** Engel, Cai, and Tihelka, *nov*.

Suborder **ADEPHAGA** Claireville, 1806

†Family Tritarsusidae Hong, 2002
†Family Triaplidae Ponomarenko, 1977
Family Haliplidae Aubé, 1836
Family Gyrinidae Latreille, 1810
Subfamily Spanglerogyrinae Folkerts, 1979
Subfamily Gyrininae Latreille, 1810
Subfamily Heterogyrinae Brinck, 1955
†Family Coptoclavidae Ponomarenko, 1961
†Subfamily Timarchopsinae Wang, Ponomarenko and Zhang, 2010
†Subfamily Charonoscaphinae Ponomarenko, 1977
†Subfamily Coptoclavinae Ponomarenko, 1961
†Subfamily Coptoclaviscinae Soriano, Ponomarenko and Delclòs, 2007
†Subfamily Hispanoclavinae Soriano, Ponomarenko and Delclòs, 2007
Family Noteridae Thomson, 1860
Subfamily Notomicrinae Branden, 1884
Subfamily Noterinae Thomson, 1860
Family Meruidae Spangler and Steiner, 2005
Family Aspidytidae Ribera, Beutel, Balke and Vogler, 2002
Family Amphizoidae LeConte, 1853
Family Hygrobiidae Régimbart, 1879
Family Dytiscidae Leach, 1815
Subfamily Matinae Branden, 1885
Subfamily Lancetinae Branden, 1885
Subfamily Agabinae Thomson, 1867

Subfamily Colymbetinae Erichson, 1837
Subfamily Copelatinae Branden, 1885
Subfamily Laccophilinae Gistel, 1848
Subfamily Cybistrinae Sharp, 1880
Subfamily Dytiscinae Leach, 1815
Subfamily Coptotominae Branden, 1885
Subfamily Hydrodytinae Miller, 2001
Subfamily Hydroporinae Aubé, 1836
†Subfamily Palaeogyrininae Schlechtendal, 1894
†Subfamily Liadytiscinae Prokin and Ren, 2010
Family Trachypachidae Thomson, 1857
†Subfamily Eodromeinae Ponomarenko, 1977
Subfamily Trachypachinae Thomson, 1857
Family Cicindelidae Latreille 1802
Family Carabidae Latreille, 1802
†Subfamily Protorabinae Ponomarenko, 1977
†Subfamily Conjunctiinae Ponomarenko, 1977
Subfamily Nebriinae Laporte, 1834
Subfamily Carabinae Latreille, 1802
Subfamily Loricerinae Bonelli, 1810
Subfamily Omophroninae Bonelli, 1810
Subfamily Elaphrinae Latreille, 1802
Subfamily Migadopinae Chaudoir, 1861
Subfamily Hiletinae Schiødte, 1848
Subfamily Scaritinae Bonelli, 1810
Subfamily Broscinae Hope, 1838
Subfamily Apotominae LeConte, 1853
Subfamily Siagoninae Bonelli, 1813
Subfamily Melaeninae Csiki, 1933
Subfamily Gehringiinae Darlington, 1933
Subfamily Trechinae Bonelli, 1810
Subfamily Patrobinae Kirby, 1837
Subfamily Psydrinae LeConte, 1853
Subfamily Nototylinae Bänninger, 1927
Subfamily Paussinae Latreille, 1806
Subfamily Brachininae Bonelli, 1810
Subfamily Harpalinae Bonelli, 1810
†Family Colymbotethidae Ponomarenko, 1994
†Family Parahygrobiidae Ponomarenko, 1977
†Family Liadytidae Ponomarenko, 1977

Suborder **ARCHOSTEMATA** Kolbe, 1908

Superfamily **Cupedoidea** Laporte, 1838
= †Family Magnocoleidae Hong, 1998 **syn. nov.**
Family Crowsoniellidae Iablokoff-Khnzorian, 1983
Family Cupedidae Laporte, 1838
Subfamily Priacminae Crowson, 1962
†Subfamily Mesocupedinae Ponomarenko, 1969
Subfamily Cupedinae Laporte, 1838
Family Micromalthidae Barber, 1913
Family Ommatidae Sharp and Muir, 1912

†Subfamily Brochocoleinae Hong, 1982
Subfamily Ommatinae Ponomarenko, 1969
Subfamily Tetraphalerinae Crowson, 1962
†Family Triadocupedidae Ponomarenko, 1966

†Superfamily **Asiocoleoidea** Rohdendorf, 1961
†Family Asiocoleidae Rohdendorf, 1961
†Family Tricoleidae Ponomarenko, 1969

†Superfamily **Rhombocoleoidea** Rohdendorf, 1961
†Family Rhombocoleidae Rohdendorf, 1961

†Superfamily **Schizocoleoidea** Rohdendorf, 1961
†Family Schizocoleidae Rohdendorf, 1961
†Family Phoroschizidae Bouchard and Bousquet, 2020
†Family Catiniidae Ponomarenko, 1968

Suborder **MYXOPHAGA** Crowson, 1955

Superfamily **Lepiceroidea** Hinton, 1936 (1882)
Family Lepiceridae Hinton, 1936 (1882)

Superfamily **Sphaeriusoidea** Erichson, 1845
Family Torridincolidae Steffan, 1964
Subfamily Torridincolinae Steffan, 1964
Subfamily Deleveinae Endrödy-Younga, 1997
†Family Triamyxidae Triamyxidae Qvarnström, Fikáček, Wernström, Huld, Beutel, Arriaga-Varela, Ahlberg, and Niedźwiedzki, 2021
Family Hydroscaphidae LeConte, 1874
Family Sphaeriusidae Erichson, 1845

Superfamily *incertae sedis*

Suborder **POLYPHAGA** Emery, 1886

Series SCIRTIFORMIA Lawrence, Ślipiński, Seago, Thayer, Newton, and Marvaldi, 2011 **sensu nov.**

Superfamily **Scirtoidea** Fleming, 1821 **sensu nov.**
Family Decliniidae Nikitsky, Lawrence, Kirejtshuk and Gratshev, 1994
Family Scirtidae Fleming, 1821
†Subfamily Mesernobiinae Engel, 2010
Subfamily Nipponocyphoninae Lawrence and Yoshitomi, 2007
Subfamily Stenocyphoninae Lawrence and Yoshitomi, 2007
Subfamily Scirtinae Fleming, 1821
†Family Elodophthalmidae Kirejtshuk and Azar, 2008
†Family Boleopsidae Kirejtshuk and Nel, 2013

Series CLAMBIFORMIA Cai and Tihelka **ser. nov.**

Superfamily **Clamboidea** Fischer, 1821 **sensu nov.**
Family Derodontidae LeConte, 1861
Subfamily Peltasticinae LeConte, 1861
Subfamily Derodontinae LeConte, 1861
Subfamily Laricobiinae Mulsant and Rey, 1864

Family Clambidae Fischer, 1821
Subfamily Calyptomerinae Crowson, 1955
Subfamily Acalyptomerinae Crowson, 1979
Subfamily Clambinae Fischer von Waldheim, 1821
Family Eucinetidae Lacordaire, 1857
†Family Mesocinetidae Kirejtshuk and Ponomarenko, 2010

Series RHINORHIPIFORMIA Cai, Engel and Tihelka **ser. nov.**

Superfamily **Rhinorhipoidea** Lawrence, 1988
Family Rhinorhipidae Lawrence, 1988

Series ELATERIFORMIA Crowson, 1960

Superfamily **Dascilloidea** Guérin-Méneville, 1843 (1834)
Family Dascillidae Guérin-Méneville, 1843 (1834)
Subfamily Dascillinae Guérin-Méneville, 1843 (1834)
Subfamily Karumiinae Escalera, 1913
Family Rhipiceridae Latreille, 1834
Subfamily Rhipicerinae Latreille, 1834
Subfamily Sandalinae Jakobson, 1913

Superfamily **Byrrhoidea** Latreille, 1804 **sensu nov.**
Family Byrrhidae Latreille, 1804
Subfamily Byrrhinae Latreille, 1804
Subfamily Syncalyptinae Mulsant and Rey, 1869
Subfamily Amphicyrtinae LeConte, 1861
†Subfamily Lidryopinae Kirejtshuk and Nel, 2016

Superfamily **Buprestoidea** Crowson, 1955
Family Schizopodidae LeConte, 1859
Family Buprestidae Leach, 1815
Subfamily Julodinae Lacordaire, 1857
Subfamily Polycestinae Lacordaire, 1857
Subfamily Galbellinae Reitter, 1911
Subfamily Chrysochroinae Laporte, 1835
Subfamily Buprestinae Leach, 1815
Subfamily Agrilinae Laporte, 1835
†Subfamily Parathyreinae Alexeev, 1994

Superfamily **Dryopoidea** Billberg, 1820 (1817) **stat. res.**
Family Lutrochidae Kasap and Crowson, 1975
Family Dryopidae Billberg, 1820 (1817)
Family Eulichadidae Crowson, 1973
Family Callirhipidae Emden, 1924
Family Ptilodactylidae Laporte, 1838
Subfamily Cladotominae Pic, 1914
Subfamily Anchytarsinae Champion, 1897
Subfamily Aploglossinae Champion, 1897
Subfamily Araeopidiinae Lawrence, 1991
Subfamily Ptilodactylinae Laporte, 1838

Family Cneoglossidae Champion, 1897
Family Chelonariidae Blanchard, 1845
Family Psephenidae Lacordaire, 1854
Subfamily Afroeubriinae Lee, Satô, Shepard and Jäch, 2007
Subfamily Eubriinae Lacordaire, 1857
Subfamily Psephenoidinae Bollow, 1938
Subfamily Eubrianacinae Jakobson, 1913
Subfamily Psepheninae Lacordaire, 1854
Family Protelmidae Jeannel, 1950
Family Elmidae Curtis, 1830
Subfamily Larainae LeConte, 1861
Subfamily Elminae Curtis, 1830
Family Limnichidae Erichson, 1846
Subfamily Hyphalinae Britton, 1971
Subfamily Limnichinae Erichson, 1846
Subfamily Cephalobyrrhinae Champion, 1925
Subfamily Thaumastodinae Champion, 1924
Family Heteroceridae MacLeay, 1825
Subfamily Elythomerinae Pacheco, 1964
Subfamily Heterocerinae MacLeay, 1825

Superfamily **Elateroidea** Leach, 1815
Family Artematopodidae Lacordaire, 1857
Subfamily Electribiinae Crowson, 1975
Subfamily Allopogoniinae Crowson, 1973
Subfamily Artematopodinae Lacordaire, 1857
Family Omethidae LeConte, 1861
Subfamily Driloniinae Crowson, 1972
Subfamily Matheteinae LeConte, 1881
Subfamily Omethinae LeConte, 1861
Subfamily Telegeusinae Leng, 1920
Family Brachypsectridae Horn, 1881
Family Throscidae Laporte, 1840
Family Eucnemidae Eschscholtz, 1829
Subfamily Perothopinae Lacordaire, 1857
Subfamily Phyllocerinae Reitter, 1905
Subfamily Pseudomeninae Muona, 1993
Subfamily Palaeoxeninae Muona, 1993
Subfamily Phlegoninae Muona, 1993
Subfamily Anischiinae Fleutiaux, 1936
Subfamily Melasinae Fleming, 1821
Subfamily Eucneminae Eschscholtz, 1829
Subfamily Macraulacinae Fleutiaux, 1923
Family Cerophytidae Latreille, 1834
Family Jurasaidae Rosa, Costa, Kramp and Kundrata, 2020
Family Elateridae Leach, 1815
Subfamily Elaterinae Leach, 1815
Subfamily Tetralobinae Laporte, 1840
Subfamily Omalisinae Lacordaire, 1857
Subfamily Agrypninae Candèze, 1857
Subfamily Hapatesinae Kusy, Motyka and Bocak, 2021
Subfamily Pityobiinae Hyslop, 1917

Subfamily Lissominae Laporte, 1835
Subfamily Thylacosterninae Fleutiaux, 1920
Subfamily Campyloxeninae Costa, 1975
Subfamily Hypnoidinae Schwarz, 1906
Subfamily Dendrometrinae Gistel, 1848
Subfamily Negastriinae Nakane and Kishii, 1956
Subfamily Cardiophorinae Candèze, 1859
Subfamily Oestodinae Hyslop, 1917
Subfamily Hemiopinae Fleutiaux, 1941
Subfamily Physodactylinae Lacordaire, 1857
Subfamily Subprotelaterinae Fleutiaux, 1920
Subfamily Morostominae Dolin, 2000
Subfamily Parablacinae Kundrata, Gunter, Douglas and Bocak, 2016
†Subfamily Protagrypninae Dolin, 1973
Family Sinopyrophoridae Bi and Li, 2019
Family Lycidae Laporte, 1838
Subfamily Dexorinae Bocak and Bocakova, 1989
Subfamily Calochrominae Lacordaire, 1857
Subfamily Erotinae Leconte, 1881
Subfamily Ateliinae Kleine, 1928
Subfamily Lycinae Laporte, 1838
Subfamily Lyropaeinae Bocak and Bocakova, 1989
Subfamily Metriorrhynchinae Kleine, 1926
Family Iberobaeniidae Bocak, Kundrata, Fernández and Vogler, 2016
†Family Berendtimiridae Winkler, 1987
†Family Cretophengodidae Li, Kundrata, Tihelka and Cai, 2021
Family Phengodidae LeConte, 1861
Subfamily Cydistinae Paulus, 1972
Subfamily Phengodinae LeConte, 1861
Subfamily Mastinocerinae LeConte, 1881
Subfamily Penicillophorinae Paulus, 1975
Family Rhagophthalmidae Olivier, 1907
Family Lampyridae Rafinesque, 1815
Subfamily Luciolinae Lacordaire, 1857
Subfamily Pterotinae LeConte, 1861
Subfamily Ototretinae McDermott, 1964
Subfamily Lamprohizinae Kazantsev, 2010
Subfamily Chespiritoinae Ferreira, Keller and Branham, 2020
Subfamily Cyphonocerinae Crowson, 1972
Subfamily Psilocladinae McDermott, 1964
Subfamily Amydetinae Olivier in Wytsman, 1907
Subfamily Cheguevariinae Kazantsev, 2007
Subfamily Photurinae Lacordaire, 1857
Subfamily Lampyrinae Rafinesque, 1815
†Family Mysteriomorphidae Alekseev and Ellenberger, 2019
Family Cantharidae Imhoff, 1856 (1815)
Subfamily Cantharinae Imhoff, 1856 (1815)
Subfamily Silinae Mulsant, 1862
Subfamily Dysmorphocerinae Brancucci, 1980
Subfamily Malthininae Kiesenwetter, 1852
Subfamily Chauliognathinae LeConte, 1861

Series NOSODENDRIFORMIA Cai and Tihelka **ser. nov.**

Superfamily **Nosodendroidea** Erichson, 1846
Family Nosodendridae Erichson, 1846

Series STAPHYLINIFORMIA Lameere, 1900 **sensu nov.**

Superfamily **Histeroidea** Gyllenhaal, 1808
Family Synteliidae Lewis, 1882
Family Sphaeritidae Shuckard, 1839
†Family Cretohisteridae Zhou, Caterino, Ślipiński and Cai, 2018
Family Histeridae Gyllenhal, 1808
Subfamily †Antigracilinae Zhou, Caterino, Ren and Ślipiński, 2020
Subfamily Niponiinae Fowler, 1912
Subfamily Abraeinae MacLeay, 1819
Subfamily Saprininae Blanchard, 1845
Subfamily Dendrophilinae Reitter, 1909
Subfamily Onthophilinae MacLeay, 1819
Subfamily Tribalinae Bickhardt, 1914
Subfamily Histerinae Gyllenhal, 1808
Subfamily Haeteriinae Marseul, 1857
Subfamily Chlamydopsinae Bickhardt, 1914

Superfamily **Hydrophiloidea** Latreille, 1802
Family Hydrophilidae Latreille, 1802
Subfamily Hydrophilinae Latreille, 1802
Subfamily Chaetarthriinae Bedel, 1881
Subfamily Enochrinae Short and Fikáček, 2013
Subfamily Acidocerinae Zaitzev, 1908
Subfamily Cylominae Zaitzev, 1908
Subfamily Sphaeridiinae Latreille, 1802
Family Helophoridae Leach, 1815
Family Epimetopidae Zaitzev, 1908
Family Georissidae Laporte, 1840
Family Hydrochidae Thomson, 1859
Family Spercheidae Erichson, 1837

Superfamily **Scarabaeoidea** Latreille, 1802
†Family Alloioscarabaeidae Bai, Ren and Yang, 2012
†Family Septiventeridae Bai, Ren, Shih and Yang, 2013
Family Lucanidae Latreille, 1804
†Subfamily Protolucaninae Nikolajev, 2007
Subfamily Aesalinae MacLeay, 1819
†Subfamily Ceruchitinae Nikolajev, 2006
Subfamily Syndesinae MacLeay, 1819
Subfamily Lampriminae MacLeay, 1819
Subfamily Lucaninae Latreille, 1804
†Subfamily Paralucaninae Nikolajev, 2000
†Subfamily Litholampriminae Nikolajev, 2015
Family Trogidae MacLeay, 1819
†Subfamily Kresnikinae Tihelka, Huang and Cai, 2019
Subfamily Troginae MacLeay, 1819
Subfamily Omorginae Nikolajev, 2005
Family Glaresidae Prudhomme de Borre, 1886

Family Pleocomidae LeConte-, 1861
†Subfamily Cretocominae Nikolajev, 2002
†Subfamily Archescarabaeinae Nikolajev, 2010
Subfamily Pleocominae LeConte, 1861
Family Bolboceratidae Mulsant, 1842
Subfamily Athyreinae Lynch Arribálzaga, 1878
Subfamily Bolboceratinae Mulsant, 1842
Family Diphyllostomatidae Holloway, 1972
Family Geotrupidae Latreille, 1802
Subfamily Taurocerastinae Germain, 1897
Subfamily Geotrupinae Latreille, 1802
†Family Passalopalpidae Boucher and Bai, 2016
Family Passalidae Leach, 1815
Subfamily Aulacocyclinae Kaup, 1868
Subfamily Passalinae Leach, 1815
Family Belohinidae Paulian, 1959
Family Ochodaeidae Streubel, 1846
†Subfamily Cretochodaeinae Nikolajev, 1995
Subfamily Ochodaeinae Streubel, 1846
Subfamily Chaetocanthinae Scholtz, 1988
Family Glaphyridae MacLeay, 1819
†Subfamily Cretoglaphyrinae Nikolajev, 2005
Subfamily Glaphyrinae MacLeay, 1819
Subfamily Amphicominae Blanchard, 1845
Family Hybosoridae Erichson, 1847
†Subfamily Mimaphodiinae Nikolajev, 2007
Subfamily Anaidinae Nikolajev, 1996
Subfamily Ceratocanthinae Martínez, 1968
Subfamily Hybosorinae Erichson, 1847
Subfamily Liparochrinae Ocampo, 2006
Subfamily Pachyplectrinae Ocampo, 2006
Family Scarabaeidae Latreille, 1802
†Subfamily Lithoscarabaeinae Nikolajev, 1992
Subfamily Chironinae Blanchard, 1845
Subfamily Aegialiinae Laporte, 1840
Subfamily Eremazinae Stebnicka, 1977
Subfamily Aphodiinae Leach, 1815
Subfamily Aulonocneminae Janssens, 1946
Subfamily Termitotroginae Wasmann, 1918
Subfamily Scarabaeinae Latreille, 1802
†Subfamily Prototroginae Nikolajev, 2000
†Subfamily Cretoscarabaeinae Nikolajev, 1995
Subfamily Dynamopodinae Arrow, 1911
†Subfamily Electrorubesopsinae Bai and Wang, 2019
Subfamily Phaenomeridinae Erichson, 1847
Subfamily Orphninae Erichson, 1847
Subfamily Allidiostomatinae Arrow, 1940
Subfamily Aclopinae Blanchard, 1850
Subfamily Melolonthinae Leach, 1819
Subfamily Oncerinae LeConte, 1861
Subfamily Podolasiinae Howden, 1997

Subfamily Rutelinae MacLeay, 1819
Subfamily Dynastinae MacLeay, 1819
Subfamily Cetoniinae Leach, 1815

Superfamily **Staphylinoidea** Latreille, 1802
Family Jacobsoniidae Heller, 1926
Family Ptiliidae Erichson, 1845
Subfamily Nossidiinae Sörensson and Delgado, 2019
Subfamily Ptiliinae Erichson, 1845
Family Hydraenidae Mulsant, 1844
Subfamily Orchymontiinae Perkins, 1997
Subfamily Prosthetopinae Perkins, 1994
Subfamily Hydraeninae Mulsant, 1844
Subfamily Ochthebiinae Thomson, 1859
Family Colonidae Horn, 1880 (1859) **stat. nov.**
Family Agyrtidae Thomson, 1859
Subfamily Agyrtinae Thomson, 1859
Subfamily Necrophilinae Newton, 1997
Subfamily Pterolomatinae Thomson, 1862
Family Leiodidae Fleming, 1821 **sensu nov.**
Subfamily Camiarinae Jeannel, 1911
Subfamily Catopocerinae Hatch, 1927 (1880)
Subfamily Leiodinae Fleming, 1821
Subfamily Cholevinae Kirby, 1837
Subfamily Platypsyllinae Ritsema, 1869
Family Staphylinidae Latreille, 1802 **sensu nov.**
Subfamily Silphinae Latreille, 1806 **stat. nov., sensu nov.**
Subfamily Apateticinae Fauvel, 1895
Subfamily Trigonurinae Reiche, 1866
Subfamily Scaphidiinae Latreille, 1806
Subfamily Piestinae Erichson, 1839
Subfamily Osoriinae Erichson, 1839
Subfamily Oxytelinae Fleming, 1821
†Subfamily Protactinae Heer, 1847
Subfamily Omaliinae MacLeay, 1825
Subfamily Microsilphinae Crowson, 1950
Subfamily Glypholomatinae Jeannel, 1962
Subfamily Empelinae Newton and Thayer, 1992
Subfamily Proteininae Erichson, 1839
Subfamily Micropeplinae Leach, 1815
Subfamily Neophoninae Fauvel, 1905
Subfamily Dasycerinae Reitter, 1887
Subfamily Protopselaphinae Newton and Thayer, 1995
Subfamily Pselaphinae Latreille, 1802
Subfamily Mycetoporinae Thomson, 1859
Subfamily Tachyporinae MacLeay, 1825
Subfamily Phloeocharinae Erichson, 1839
Subfamily Trichophyinae Thomson, 1858
Subfamily Habrocerinae Mulsant and Rey, 1876
Subfamily Aleocharinae Fleming, 1821
Subfamily Leptotyphlinae Fauvel, 1874
Subfamily Oxyporinae Fleming, 1821

Subfamily Megalopsidiinae Leng, 1920
Subfamily Steninae MacLeay, 1825
Subfamily Euaesthetinae Thomson, 1859
Subfamily Solieriinae Newton and Thayer, 1992
Subfamily Scydmaeninae Leach, 1815
Subfamily Pseudopsinae Ganglbauer, 1895
Subfamily Olisthaerinae Thomson, 1858
Subfamily Paederinae Fleming, 1821
Subfamily Staphylininae Latreille, 1802

Series BOSTRICHIFORMIA Forbes, 1926

Superfamily **Bostrichoidea** Latreille, 1802
Family Dermestidae Latreille, 1804
Subfamily Dermestinae Latreille, 1804
Subfamily Thorictinae Agassiz, 1846
Subfamily Orphilinae LeConte, 1861
Subfamily Trinodinae Casey, 1900
Subfamily Attageninae Laporte, 1840
Subfamily Megatominae Leach, 1815
Family Endecatomidae LeConte, 1861
Family Bostrichidae Latreille, 1802
Subfamily Dysidinae Lesne, 1894
Subfamily Polycaoninae Lesne, 1896
Subfamily Bostrichinae Latreille, 1802
Subfamily Psoinae Blanchard, 1851
Subfamily Dinoderinae Thomson, 1863
Subfamily Lyctinae Billberg, 1820
Subfamily Euderiinae Lesne, 1934
Family Ptinidae Latreille, 1802
Subfamily Eucradinae LeConte, 1861
Subfamily Ptininae Latreille, 1802
Subfamily Dryophilinae Gistel, 1848
Subfamily Ernobiinae Pic, 1912
Subfamily Anobiinae Fleming, 1821
Subfamily Ptilininae Shuckard, 1839
Subfamily Alvarenganiellinae Viana and Martínez, 1971
Subfamily Xyletininae Gistel, 1848
Subfamily Dorcatominae Thomson, 1859
Subfamily Mesocoelopodinae Mulsant and Rey, 1864

Series CUCUJIFORMIA Lameere, 1938

Superfamily **Cleroidea** Latreille, 1802
Family Rentoniidae Crowson, 1966
Family Byturidae Gistel, 1848
Subfamily Byturinae Gistel, 1848
Subfamily Platydascillinae Pic, 1914
Family Biphyllidae LeConte, 1861
Family Acanthocnemidae Crowson, 1964
Family Protopeltidae Crowson, 1966
Family Peltidae Kirby, 1837
Family Lophocateridae Crowson, 1964

Family Trogossitidae Latreille, 1802
Subfamily Calityinae Houlbert, 1922
Subfamily Egoliinae Lacordaire, 1854
Subfamily Larinotinae Slipinski, 1992
Subfamily Trogossitinae Latreille, 1802
Family Thymalidae Léveillé, 1888
Subfamily Thymalidae Léveillé, 1888
Subfamily Decamerinae Crowson, 1964
Family Phycosecidae Crowson, 1952
Family Prionoceridae Lacordaire, 1857
Family Mauroniscidae Majer, 1995
Family Rhadalidae LeConte, 1861
Subfamily Gietellinae Constantin and Menier, 1987
Subfamily Rhadalinae LeConte, 1861
Family Melyridae Leach, 1815
Subfamily Malachiinae Fleming, 1821
Subfamily Melyrinae Leach, 1815
Subfamily Dasytinae Laporte, 1840
Family Phloiophilidae Kiesenwetter, 1863
Family Chaetosomatidae Crowson, 1952
Family Thanerocleridae Chapin, 1924
Subfamily Zenodosinae Kolibáč, 1992
Subfamily Thaneroclerinae Chapin, 1924
Family Cleridae Latreille, 1802
Subfamily Tillinae Fischer von Waldheim, 1813
Subfamily Korynetinae Laporte, 1838
Subfamily Epiclininae Gunter, Leavengood and Bartlett, 2013
Subfamily Hydnocerinae Spinola, 1844
Subfamily Clerinae Latreille, 1802

Superfamily **Lymexyloidea** Fleming, 1821
Family Lymexylidae Fleming, 1821
Subfamily Hylecoetinae Germar, 1818
Subfamily Lymexylinae Fleming, 1821
Subfamily Atractocerinae Laporte, 1840
Subfamily Melittommatinae Wheeler, 1986

Superfamily **Tenebrionoidea** Latreille, 1802
Family Ripiphoridae Laporte, 1840
Subfamily Pelecotominae Guérin-Méneville, 1857
Subfamily Ptilophorinae Gerstaecker, 1855
Subfamily Hemirhipidiinae Heller, 1921
Subfamily Ripidiinae Gerstaecker, 1855
Subfamily Ripiphorinae Laporte, 1840
Family Mordellidae Latreille, 1802
Subfamily Ctenidiinae Franciscolo, 1951
Subfamily Mordellinae Latreille, 1802
†Family Apotomouridae Bao, Walczynska, Moody, Wang and Rust, 2018
Family Aderidae Csiki, 1909
Family Ischaliidae Blair, 1920
Family Trictenotomidae Blanchard, 1845
Family Scraptiidae Gistel, 1848
Subfamily Scraptiinae Gistel, 1848

Subfamily Anaspidinae Mulsant, 1856
Family Mycteridae Perty, 1840
Subfamily Mycterinae Perty, 1840
Subfamily Eurypinae Thomson, 1860
Subfamily Hemipeplinae Lacordaire, 1854
Family Oedemeridae Latreille, 1810
Subfamily Polypriinae Lawrence, 2005
Subfamily Calopodinae Costa, 1852
Subfamily Oedemerinae Latreille, 1810
Family Boridae Thomson, 1859
Subfamily Borinae Thomson, 1859
Subfamily Synercticinae Lawrence and Pollock, 1994
Family Pythidae Solier, 1834
Family Salpingidae Leach, 1815
Subfamily Othniinae LeConte, 1861
Subfamily Prostominiinae Grouvelle, 1914
Subfamily Agleninae Horn, 1878
Subfamily Inopeplinae Grouvelle, 1908
Subfamily Dacoderinae LeConte, 1862
Subfamily Aegialitinae LeConte, 1862
Subfamily Salpinginae Leach, 1815
Family Pyrochroidae Latreille, 1806
Subfamily Tydessinae Nikitsky, 1986
Subfamily Pilipalpinae Abdullah, 1964
†Subfamily Waidelotinae Bukejs, Alekseev and Pollock, 2019
Subfamily Pedilinae Lacordaire, 1859
Subfamily Pyrochroinae Latreille, 1806
Subfamily Pogonocerinae Iablokov-Khnzorian, 1985
Subfamily Agnathinae Lacordaire, 1859
Family Anthicidae Latreille, 1819
Subfamily Eurygeniinae LeConte, 1862
Subfamily Macratriinae LeConte, 1862
Subfamily Steropinae Jacquelin du Val, 1863
Subfamily Copobaeninae Abdullah, 1969
Subfamily Lemodinae Matthews, 1987
Subfamily Tomoderinae Bonadona, 1961
Subfamily Anthicinae Latreille, 1819
Subfamily Notoxinae Stephens, 1829
Family Meloidae Gyllenhal, 1810
Subfamily Eleticinae Wellman, 1910
Subfamily Meloinae Gyllenhal, 1810
Subfamily Tetraonycinae Böving and Craighead, 1931
Subfamily Nemognathinae Laporte, 1840
Family Stenotrachelidae Thomson, 1859
Subfamily Stenotrachelinae Thomson, 1859
Subfamily Cephaloinae LeConte, 1862
Subfamily Nematoplinae LeConte, 1862
Subfamily Stoliinae Nikitsky, 1985
Family Tetratomidae Billberg, 1820
Subfamily Tetratominae Billberg, 1820
Subfamily Piseninae Miyatake, 1960

Subfamily Penthinae Lacordaire, 1859
Subfamily Hallomeninae Gistel, 1848
Subfamily Eustrophinae Gistel, 1848
Family Melandryidae Leach, 1815
Subfamily Melandryinae Leach, 1815
Subfamily Osphyinae Mulsant, 1856 (1839)
Family Synchroidae Lacordaire, 1859
Family Prostomidae Thomson, 1859
Family Ciidae Leach, 1819
Subfamily Sphindociinae Lawrence, 1974
Subfamily Ciinae Leach, 1819
Family Ulodidae Pascoe, 1869
Family Archeocrypticidae Kaszab, 1964
Family Pterogeniidae Crowson, 1953
Family Mycetophagidae Leach, 1815
Subfamily Esarcinae Reitter, 1882
Subfamily Mycetophaginae Leach, 1815
Subfamily Bergininae Leng, 1920
Family Tenebrionidae Latreille, 1802
Subfamily Lagriinae Latreille, 1825 (1820)
Subfamily Nilioninae Oken, 1843
Subfamily Phrenapatinae Solier, 1834
Subfamily Zolodininae Watt, 1975
Subfamily Pimeliinae Latreille, 1802
Subfamily Blaptinae Leach, 1815
Subfamily Tenebrioninae Latreille, 1802
Subfamily Alleculinae Laporte, 1840
Subfamily Diaperinae Latreille, 1802
Subfamily Stenochiinae Kirby, 1837
Family Zopheridae Solier, 1834
Subfamily Colydiinae Billberg, 1820
Subfamily Zopherinae Solier, 1834
Family Promecheilidae Lacordaire, 1859
Family Chalcodryidae Watt, 1974
Family *incertae sedis*
Subfamily Afreminae Levey, 1985
Subfamily Lagrioidinae Abdullah and Abdullah, 1968

Superfamily **Coccinelloidea** Latreille, 1807
Family Bothrideridae Erichson, 1845
Family Cerylonidae Billberg, 1820
Subfamily Ostomopsinae Sen Gupta and Crowson, 1973
Subfamily Loebliorylonina Ślipiński, 1990
Subfamily Ceryloninae Billberg, 1820
Family Murmidiidae Jacquelin du Val, 1858
Family Discolomatidae Horn, 1878
Subfamily Notiophyginae Jakobson, 1915
Subfamily Discolomatinae Horn, 1878
Subfamily Aphanocephalinae Jakobson, 1904
Subfamily Cephalophaninae John, 1954
Subfamily Pondonatinae John, 1954
Family Euxestidae Grouvelle, 1908

Family Teredidae Seidlitz, 1888
Subfamily Teredinae Seidlitz, 1888
Subfamily Anommatinae Ganglbauer, 1899
Subfamily Xylariophilinae Pal and Lawrence, 1986
Family Alexiidae Imhoff, 1856
Family Akalyptoischiidae Lord, Hartley, Lawrence, McHugh, Whiting and Miller, 2010
Family Latridiidae Erichson, 1842
Subfamily Latridiinae Erichson, 1842
Subfamily Corticariinae Curtis, 1829
Family Anamorphidae Strohecker, 1953
Family Corylophidae LeConte, 1852
Subfamily Periptyctinae Ślipiński, Lawrence and Tomaszewska, 2001
Subfamily Corylophinae LeConte, 1852
Family Endomychidae Leach, 1815
Subfamily Merophysiinae Seidlitz, 1872
Subfamily Pleganophorinae Jacquelin du Val, 1858
Subfamily Leiestinae Thomson, 1863
Subfamily Xenomycetinae Strohecker, 1962
Subfamily Danascelinae Tomaszewska, 2000
Subfamily Cyclotominae Imhoff, 1856
Subfamily Endomychinae Leach, 1815
Subfamily Epipocinae Gorham, 1873
Subfamily Lycoperdininae Bromhead, 1838
†Family Tetrameropsidae Kirejtshuk and Azar, 2008
Family Mycetaeidae Jacquelin du Val, 1857
Family Eupsilobiidae Casey, 1895
Family Coccinellidae Latreille, 1807
Subfamily Microweiseinae Leng, 1920
Subfamily Monocoryninae Miyatake, 1988
Subfamily Coccinellinae Latreille, 1807

Superfamily **Erotyloidea** Latreille, 1802 **stat. nov.**
Family Boganiidae Sen Gupta and Crowson, 1966
Subfamily Paracucujinae Endrödy-Younga and Crowson, 1986
Subfamily Boganiinae Sen Gupta and Crowson, 1966
Family Erotylidae Latreille, 1802
Subfamily Cryptophilinae Casey, 1900
Subfamily Languriinae Hope, 1840
Subfamily Xenoscelinae Ganglbauer, 1899
Subfamily Pharaxonothinae Crowson, 1952
Subfamily Loberinae Bruce, 1951
Subfamily Erotylinae Latreille, 1802

Superfamily **Nitiduloidea** Latreille, 1802 **stat. nov.**
Family Helotidae Chapuis, 1876
Family Sphindidae Jacquelin du Val, 1860
†Subfamily Libanopsinae Kirejtshuk, 2015
Subfamily Protosphindinae Sen Gupta and Crowson, 1979
Subfamily Odontosphindinae Sen Gupta and Crowson, 1979
Subfamily Sphindiphorinae Sen Gupta and Crowson, 1979
Subfamily Sphindinae Jacquelin du Val, 1860
Family Protocucujidae Crowson, 1954

Family Monotomidae Laporte, 1840
Subfamily Rhizophaginae Redtenbacher, 1845
Subfamily Monotominae Laporte, 1840
Family Kateretidae Kirby, 1837
Family Nitidulidae Latreille, 1802
Subfamily Cryptarchinae Thomson, 1859
Subfamily Prometopiinae Böving and Craighead, 1931
Subfamily Amphicrossinae Kirejtshuk, 1986
Subfamily Carpophilinae Erichson, 1842
Subfamily Cybocephalinae Jacquelin du Val, 1858 **stat. nov.**
Subfamily Calonecrinae Kirejtshuk, 1982
Subfamily Epuraeinae Kirejtshuk, 1986
Subfamily Meligethinae Thomson, 1859
Subfamily Nitidulinae Latreille, 1802
Subfamily Maynipeplinae Kirejtshuk, 1998
Subfamily Cillaeinae Kirejtshuk and Audisio, 1986
Family Smicripidae Horn, 1880

Superfamily **Cucujoidea** Latreille, 1802 **sensu nov.**
Family Hobartiidae Sen Gupta and Crowson, 1966
Family Cryptophagidae Kirby, 1826
Subfamily Cryptophaginae Kirby, 1826
Subfamily Atomariinae LeConte, 1861
Family Silvanidae Kirby, 1837
Subfamily Brontinae Blanchard, 1845
Subfamily Silvaninae Kirby, 1837
Family Cucujidae Latreille, 1802
Subfamily Cucujinae Latreille, 1802
Subfamily Pediacinae Jin, Zwick, Ślipiński, Marris, Thomas and Pang, 2020
Subfamily Platisinae Jin, Zwick, Slipiński, Marris, Thomas and Pang, 2020
Family Phloeostichidae Reitter, 1911
Family Agapythidae Sen Gupta and Crowson, 1969
Family Priasilphidae Crowson, 1973
Family Cavognathidae Sen Gupta and Crowson, 1966
Family Lamingtoniidae Sen Gupta and Crowson, 1969
Family Tasmosalpingidae Lawrence and Britton, 1991
Family Cyclaxyridae Gimmel, Leschen and Ślipiński, 2009
Family Passandridae Blanchard, 1845
†Subfamily Mesopassandrinae Jin, Ślipiński, Zhou and Pang, 2019
Subfamily Passandrinae Blanchard, 1845
Family Myraboliidae Lawrence and Britton, 1991
Family Phalacridae Leach, 1815
Subfamily Phaenocephalinae Matthews, 1899
Subfamily Phalacrinae Leach, 1815
Family Laemophloeidae Ganglbauer, 1899

Series **Cucujiformia** Lameere, 1938 ***incertae sedis***
†Family Parandrexidae Kirejtshuk, 1994
†Family Wabbelidae Alekseev, 2017

Superfamily **Curculionoidea** Latreille, 1802
Family Cimberididae Gozis, 1882
Family Nemonychidae Bedel, 1882

†Subfamily Aepyceratinae Poinar, Brown and Legalov, 2017
Subfamily Rhinorhynchinae Voss, 1922
†Subfamily Brenthorrhininae Arnoldi, 1977
†Subfamily Distenorrhininae Arnoldi, 1977
†Subfamily Eobelinae Arnoldi, 1977
†Subfamily Paleocartinae Legalov, 2003
†Subfamily Metrioxenoidinae Legalov, 2009
†Subfamily Cretonemonychinae Gratshev and Legalov, 2009
†Subfamily Selengarhynchinae Gratshev and Legalov, 2009
†Subfamily Slonikinae Zherikhin, 1977
†Subfamily Eccoptarthrinae Arnoldi, 1977
Subfamily Nemonychinae Bedel, 1882
Family Anthribidae Billberg, 1820
Subfamily Urodontinae Thomson, 1859
Subfamily Choraginae Kirby, 1819
Subfamily Anthribinae Billberg, 1820
Family Belidae Schönherr, 1826
†Subfamily Montsecbelinae Legalov, 2015
Subfamily Oxycoryninae Schönherr, 1840
Subfamily Belinae Schönherr, 1826
†Family Mesophyletidae Poinar, 2008
†Subfamily Aepyceratinae Poinar, Brown and Legalov, 2017
†Subfamily Mesophyletinae Poinar, 2008
Family Attelabidae Billberg, 1820
Subfamily Attelabinae Billberg, 1820
Subfamily Apoderinae Jekel, 1860
Subfamily Rhynchitinae Gistel, 1848
Subfamily Isotheinae Scudder, 1893
Subfamily Pterocolinae Lacordaire, 1865
Family Caridae Thompson, 1992
†Subfamily Baissorhynchinae Zherikhin, 1993
Subfamily Chilecarinae Legalov, 2009
Subfamily Carinae Thompson, 1992
Family Brentidae Billberg, 1820
Subfamily Ithycerinae Schoenherr, 1823
Subfamily Brentinae Billberg, 1820
Subfamily Microcerinae Lacordaire, 1863
Subfamily Eurhynchinae Lacordaire, 1863
Subfamily Apioninae Schönherr, 1823
Subfamily Nanophyinae Gistel, 1848
Family Curculionidae Latreille, 1802
Subfamily Brachycerinae Billberg, 1820
Subfamily Dryophthorinae Schönherr, 1825
Subfamily Platypodinae Shuckard, 1839
Subfamily Bagoinae Thomson, 1859
Subfamily Hyperinae Marseul, 1863 (1848)
Subfamily Entiminae Schönherr, 1823
Subfamily Cyclominae Schönherr, 1826
Subfamily Curculioninae Latreille, 1802
Subfamily Baridinae Schönherr, 1836
Subfamily Ceutorhynchinae Gistel, 1848

Subfamily Conoderinae Schönherr, 1833
Subfamily Cossoninae Schönherr, 1825
Subfamily Molytinae Schönherr, 1823
Subfamily Cryptorhynchinae Schönherr, 1825
Subfamily Lixinae Schönherr, 1823
Subfamily Mesoptiliinae Lacordaire, 1863
Subfamily Scolytinae Latreille, 1804
Subfamily Orobitidinae Thomson, 1859
Subfamily Xiphaspidinae Marshall, 1920
†Family Ulyanidae Zherikhin, 1993
†Family Obrieniidae Zherikhin and Gratshev, 1994
†Subfamily Kararhynchinae Zherikhin and Gratshev, 1994
†Subfamily Obrieniinae Zherikhin and Gratshev, 1994
†Subfamily Pseudobrieniinae Legalov, 2012

Superfamily **Chrysomeloidea** Latreille, 1802
Family Oxypeltidae Lacordaire, 1868
Family Vesperidae Mulsant, 1839
Subfamily Philinae Thomson, 1861
Subfamily Vesperinae Mulsant, 1839
Subfamily Anoplodermatinae Guérin-Méneville, 1840
Family Disteniidae Thomson, 1861
Family Cerambycidae Latreille, 1802
Subfamily Dorcasominae Lacordaire, 1868
Subfamily Prioninae Latreille, 1802
Subfamily Cerambycinae Latreille, 1802
Subfamily Lepturinae Latreille, 1802
Subfamily Spondylidinae Audinet-Serville, 1832
Subfamily Lamiinae Latreille, 1825
Family Megalopodidae Latreille, 1802
Subfamily Megalopodinae Latreille, 1802
Subfamily Palophaginae Kuschel and May, 1990
Subfamily Zeugophorinae Böving and Craighead, 1931
Family Orsodacnidae Thomson, 1859
Subfamily Orsodacninae Thomson, 1859
Subfamily Aulacoscelidinae Chapuis, 1874
Family Chrysomelidae Latreille, 1802
Subfamily Chrysomelinae Latreille, 1802
Subfamily Galerucinae Latreille, 1802
Subfamily Synetinae LeConte and Horn, 1883
Subfamily Sagrinae Leach, 1815
Subfamily Bruchinae Latreille, 1802
Subfamily Donaciinae Kirby, 1837
Subfamily Criocerinae Latreille, 1804
Subfamily Cassidinae Gyllenhal, 1813
Subfamily Eumolpinae Hope, 1840
Subfamily Lamprosomatinae Lacordaire, 1848
Subfamily Cryptocephalinae Gyllenhal, 1813
Subfamily Spilopyrinae Chapuis, 1874
†Subfamily Protoscelidinae Medvedev, 1968

**INCERATE SEDIS**

Crown **COLEOPTERA** ***incerate sedis***

†Subfamily Avitortorinae Nikolaev 2007

Suborder **POLYPHAGA** Emery, 1886 ***incertae sedis***

†Family Lasiosynidae Kirejtshuk, Chang, Ren and Kun, 2010

†Family Peltosynidae Yan, Beutel and Ponomarenko, 2018

Family Jurodidae Ponomarenko, 1985

### Phylogenetic and age justifications for fossil calibrations

#### **Node 1:** (Megaloptera + Neuroptera) **–** Coleoptera: **298.75 Ma – 324 Ma.**

***Fossil taxon and specimen***: *Stephanastus polinae* Kirejtshuk and Nel, 2013 [holotype, MNHN.A.49011 (exoskeleton with legs and elytra): Palaeoentomological Laboratory, Muséum national d'Histoire naturelle, Paris, France] from the Commentry Shales Formation in Allier, France (Nel et al. 2013).
***Phylogenetic justification***: Megaloptera and Neuroptera, along with Raphidioptera, form the clade Neuropterida, which is strongly supported by both molecular and morphology-based phylogenies. As discussed in Nel et al. (2013), the extinct *Stephanastus* (family Stephanastidae) is sister to the stem coleopterids, which include the extant Coleoptera. Since the Neuropterida is a sister group to Stephanastidae + coleopterids, the fossil species *Stephanastus polinae* is a crown group of the clade ((Megaloptera, Neuroptera), Coleoptera).
***Minimum age***: 298.75 Ma.
***Soft maximum age***: 324 Ma.
***Age justification***: *Stephanastus polinae* occurs in the Commentry Shales Formation in Allier, France. Although no geochemical dating of the fossil deposit has been carried out, the formation has been regarded as Gzhelian (late Carboniferous) based on stratigraphic evidence. The top of Gzhelian has been dated to 298.9 ± 0.15, which provides a conservative minimum constraint (298.75 Ma). The soft maximum constraint is based on the earliest occurrence of Archaeorthoptera from the early Carboniferous, which also represents the earliest known Pterygota (including almost all insects (Ectognatha), but not the Archaeognatha or the Zygentoma, two primitively wingless insect orders). This earliest pterygote fossil was described from the Upper Silesian Basin in the Czech Republic (Prokop et al. 2005). The deposit has been dated as the early Carboniferous (Namurian A/E1, ca. 324 Ma) (Jirásek et al. 2013). Regardless the alternative interpretations of the Czech specimen (Dvořák et al. 2019), the other oldest uncontested candidate for the earliest pterygote, is the practically contemporaneous palaeopteran *Delitzschala bitterfeldensis* from near Bitterfeld, Germany (Brauckmann et al. 1994; Brauckmann and Schneider 1996).

#### **Node 2:** Crown Coleoptera: **251.878 Ma – 307.1 Ma.**

***Fossil taxon and specimen***: *Ponomarenkium belmonthensis* Yan, Lawrence, Beattie and Beutel, 2018 [holotype, 40278; paratype, 41618 (exoskeletons without legs): Australian Museum, Sydney] from the Tatarian insect beds, Newcastle Coal Measures at Belmont, north of Sydney, Australia (Yan et al. 2018a, b).
***Phylogenetic justification***: This clade comprises four extant beetle suborders Adephaga, Archostemata, Myxophaga, and Polyphaga, their last common ancestor and all of its descendants, but excludes putative stem coleopterans known collectively as Protocoleoptera (Crowson 1975). Yan et al. (2018a) did not provide a formal phylogenetic analysis of *Ponomarenkium* but compared it with related fossil taxa and representatives of four modern suborders. *Ponomarenkium* lacks the ancestral broad and apically truncated prosternal process and a broad prothoracic postcoxal bridge, which suggest it as a crown group Coleoptera (Yan et al. 2018a). Specifically, *Ponomarenkium* may belong to the stem group of either Polyphaga or Myxophaga, or more likely the stem group of a clade comprising more than one of the extant suborders (Yan et al. 2018a).
***Minimum age***: 251.878 Ma.
***Soft maximum age***: 307.1 Ma.
***Age justification***: *Ponomarenkium belmonthensis* was recovered from the late Permian Tatarian insect beds located within the Upper Newcastle Coal Measures of New South Wales, Australia (Yan et al. 2018d). The Belmont insect bed was considered to be Tatarian-aged (late Permian), but no precise dating of the fossil bed has been published. The minimum age for the fossil can be taken from the top of the Changhsingian (dated at 251.902 ± 0.024 Ma), providing a conservative minimum age for *Ponomarenkium*. The earliest definitive beetle, *Coleopsis archaica* is from the early Permian of Germany (Kirejtshuk et al. 2014; Kirejtshuk and Nel 2016). Like most Permian beetles, *C. archaica* belongs to the stem group of Coleoptera. Béthoux (2009) regarded *Adiphlebia lacoana* from the Pennsylvanian (Carboniferous) of Mazon Creek as the oldest beetle, but this hypothesis was rejected based on a critical re-analysis of the anatomy and taphonomy of the fossil.

The original interpretation of *Adiphlebia* as Coleoptera by Béthoux (2009) was based on clay artefacts attached to the wing membrane (Kukalová-Peck and Beutel, 2012). The Mazon Creek Lagerstätte is one of the best records of late Palaeozoic ecosystems, which has yielded diverse insects (11 orders and more than 210 species) preserved within siderite concretions (Clements et al. 2019). Despite the high palaeodiversity of the entomofauna, no beetles, one of most diverse animal group in today's terrestrial ecosystems, have been documented. Palynological and palaeobotanical evidence indicates that the age of the Mazon Creek Lagerstätte equates to the upper part of the Moscovian stage (Clements et al. 2019), the top of which has been dated at 307.0 ± 0.1. The age 307.1 Ma thus provides a conservative soft maximum constraint.

#### **Node 3:** Gyrinidae – Haliplidae: **251.878 Ma – 293.69 Ma.**

***Fossil taxon and specimen***: *Tunguskagyrus planus* Yan, Beutel and Lawrence, 2018 [holotype, PIN 5381/32 (exoskeleton without legs): Borissiak Paleontological Institute, Russian Academy of Sciences, Moscow, Russia] from the Lebedevskian Horizon at the Anakit locality, which has been correlated with the Kedrovskian layers of the Maltsevo Formation, Krasnoyarsk region, Russia (Yan et al. 2018c).
***Phylogenetic justification***: The placement of *Tunguskagyrus planus* in the extant family Gyrinidae is supported by numerous derived features of the head, but it bears many ancestral characters, indicating it as a stem group gyrinid (Yan et al. 2018c). *Tunguskagyrus* demonstrates that the specialised lifestyle of whirligig beetles probably changed little over the past 250 million. As such, *T. planus* is the oldest record of crown group of the Gyrinidae + Haliplidae clade.
***Minimum age***: 251.878 Ma.
***Soft maximum age***: 293.69 Ma.
***Age justification***: *Tunguskagyrus planus* occurs in the Lebedevskian Horizon. The age of this layer was identified as either Permian or Triassic. In a recent biostratigraphical study by Sadovnikov (2016), it was attributed to the Changhsingian of the uppermost Permian. The minimum age for the fossil can be taken at the top of the Changhsingian, which is currently dated at 251.902 ± 0.024 (International Chronostratigraphic Chart, v. 2018/08). Therefore, the minimum age constraint for this node is 251.878 Ma. The soft maximum constraint is based on the oldest occurrence of a stem group beetle from the early Permian of Germany, as in node 3.
The earliest known beetle, *Coleopsis archaica* (family Coleopsidae), has been reported from the early Permian of Germany (Kirejtshuk et al. 2014). As in many Permian species and some Mesozoic early beetle species, *C. archaica* has well sclerotised elytra with remains of primary venation, which is not developed in crown group beetles. Although *C. archaica* was placed in the extant suborder Archostemata by Kirejtshuk et al. (2014), Beutel et al. (2008) conducted a morphological phylogenetic analysis that supported Tshekardocoleidae as a stem Coleoptera, sister to the other remaining extinct and extant beetles sampled in the study. Permian beetles are largely represented by extinct stem group beetle families, such as Permocupedidae and Tshekardocoleidae (Kirejtshuk et al. 2014). Recently, an intriguing well-preserved fossil beetle, *Ponomarenkia belmonthensis* Yan et al. 2018 (family Ponomarenkiidae), was described from the mid-Permian of Australia that can be excluded from the earliest stem group coleopterans and the four extant suborders. As such, *P. belmonthensis* likely belongs to the stem group of one of the extant suborders or in the stem group of a clade comprising more than one of them (Yan et al. 2018a). Given the discussion above, the age of the oldest beetle, *Coleopsis archaica*, provides a maximum age constraint on the node. *Coleopsis archaica* was recovered from the top of the Meisenheim Formation at Grügelborn, Germany. Schneider and Werneburg (2006, 2012) proposed a Sakmarian age for this horizon based on the occurrence of the biostratigraphically important cockroach *Spiloblattina odernheimensis* Schneider. Therefore, we consider the bottom of the Sakmarian, 293.69 Ma (293.52 ± 0.17 Ma), as the soft maximum constraint.

#### **Node 4:** Dytiscidae – Noteridae: **155 Ma** – **251.878 Ma.**

***Fossil taxon and specimen***: *Palaeodytes gutta* Ponomarenko, 1987 [holotype, PIN2066/2654 (exoskeleton): Paleontological Institute, Russian Academy of Sciences, Moscow, Russia] from the Karabastau Formation of the Karatau Range near the village of Mikhailovka, Chimkent Region, Chayan District, southern Kazakhstan (Ponomarenko 1987).

***Phylogenetic justification***: The extinct genus *Palaeodytes*, currently classified as subfamily *incertae sedis* within Dytiscidae, includes three Mesozoic species: *P. baissensis*, *P. gutta*, and *P. sibiricus* (Prokin et al. 2013). Among them, the Jurassic *P. gutta* represents the oldest unequivocal record for the genus. The phylogenetic position of *Palaeodytes* within Dytiscidae remains untested, but *P. gutta* represents a crown group of Dytiscidae + Noteridae. Ponomarenko (1963) described a dytiscid species, *Angaragabus jurassicus*, from the Lower Jurassic of Irkutsk Oblast (Russia) based on a fossil larva. A re-examination of the fossil confirmed its placement in the superfamily Dytiscoidea, but not necessarily within the extant family Dytiscidae. Instead, *A. jurassicus* probably belongs to the extinct group Liadytidae (Prokin et al. 2013).
***Minimum age***: 155 Ma.
***Soft maximum age***: 251.878 Ma.
***Age justification***: *Palaeodytes gutta* is from the Karabastau Formation in Karatau, Kazakhstan. The precise age of the Karabastau Formation is uncertain within the Late Jurassic, but floras (Doludenko and Orlovskaya, 1976), insects (Rohdendorf, 1968) and stratigraphy (Kirichkova and Doludenko, 1996) suggest a mid–Late Jurassic date, around 155 Ma. Zheng et al. (2018) reported an unnamed beetle belonging to the superfamily Dytiscoidea from the mid-Triassic Tongchuan entomofauna (Shaanxi, central China), which was dated as approximately 237–238 Ma. This dytiscoid fossil, characterised by a streamlined body, smooth elytra, and second abdominal sternite divided by metacoxae, closely resembles some representatives of Noteridae. The soft maximum constraint (251.878 Ma) is based on the oldest occurrence of the suborder Adephaga (a stem group of Gyrinidae) from Late Permian deposits of the Anakit area in Middle Siberia as discussed in node 3.

#### **Node 5:** Cupedidae – Torridincolidae: **235 Ma – 293.69 Ma.**
***Fossil taxon and specimen***: *Kirghizocupes cellulosus* Ponomarenko, 1966 [holotype, PIN 2069/107 (exoskeleton): Borissiak Paleontological Institute, Russian Academy of Sciences, Moscow, Russia] from the Madygen Formation at the Dzhailyau-Cho locality, Lyailyakskii District, Batken Region, southwest Kyrgyzstan (Ponomarenko 1966).
***Phylogenetic justification***: *Kirghizocupes cellulosus* was originally placed in the extinct family Triadocupedidae (Ponomarenko 1966), which is currently regarded as stem group Coleoptera sister to all four extant suborders or as a member of Archostemata (Yan et al. 2018d). Kirejtshuk et al. (2016) attributed *Kirghizocupes* to the subfamily Cupedinae (= Cupedidae *s.s.*, here adopted, *viz* Classification of beetles) based on an extensive comparative study of fossil cupedids, representing the earliest known record of the group. Although the systematic position of *K. cellulosus* among Cupedidae *s.s.* remains untested, it represents a crown group of the Cupedidae + Torridincolidae clade.
***Minimum age***: 235 Ma.
***Soft maximum age***: 293.69 Ma.
***Age justification***: The earliest records of *Kirghizocupes* occur in the Madygen Formation at Dzhailyau-Cho in Kyrgyzstan. The Madygen Formation was dated as Ladinian–Carnian based on the evidence of megaflora (Dobruskina 1995), but the insect assemblage, namely the orders Blattodea, Orthoptera, and Hemiptera, were suggested to be more primitive than the Carnian-aged entomofaunas from South Africa and Australia, which suggests that a Ladinian age is more probable for the Madygen Formation (Shcherbakov 2008). Recent biostratigraphic and radioisotopic data indicated a mid-Triassic age (237 ± 2 Ma, late Ladinian to early Carnian) of the Madygen Formation (Voigt et al. 2017). Thus, the minimum age constraint on the node is 235 Ma. The soft maximum constraint is based on the oldest occurrence of a stem group beetle from the early Permian of Germany, as in node 3.

#### **Node 6:** Crown Polyphaga: **237 Ma – 293.69 Ma.**
***Fossil taxon and specimen***: undescribed taxon of the series Elateriformia [NIGP162053 (exoskeleton without legs): Nanjing Institute of Geology and Palaeontology, Chinese Academy of Sciences, Nanjing, China] from the Ladinian Tongchuan entomofauna at Hejiafang, Shaanxi, central China (Zheng et al. 2018).
***Phylogenetic justification***: Although the fossil was not formally named, it is well preserved and displays many features of Elateriformia (e.g., Dascilloidea and Elateroidea), including a robust body with a distinct prosternal intercoxal process and produced hind pronotal angles. The fossil clearly represents a crown group

polyphagan.
***Minimum age***: 237 Ma.
***Soft maximum age***: 293.69 Ma.
***Age justification***: The elateriform beetle originates from the Tongchuan Formation in central China**.** The insect-bearing layer of this formation was dated to 237 to 238 Ma, providing a minimum age of 237 Ma. The soft maximum constraint is based on the oldest occurrence of a stem group beetle from the early Permian of Germany, as in node 3.

#### **Node 7:** Derodontidae – (Clambidae + Eucinetidae): **165 Ma – 251.878 Ma.**

***Fossil taxon and specimen***: *Juropeltastica sinica* Cai, Lawrence and Huang, 2014 [holotype, NIGP157738 (exoskeleton without legs): Nanjing Institute of Geology and Palaeontology, Chinese Academy of Sciences, Nanjing, China] from the Haifanggou Formation at Daohugou, Ningcheng County, Inner Mongolia, northeastern China (Cai et al. 2014a).
***Phylogenetic justification***: *Juropeltastica sinica* from the Middle Jurassic represents the only Mesozoic fossil of the family Derodontidae. Derodontidae currently comprises three subfamilies: Peltasticinae (*Peltastica*), Derodontinae (*Derodontus*), and Laricobiinae (*Laricobius* and *Nothoderodontus*). *Juropeltastica* is placed in the extant subfamily Peltasticinae based on derived characters such as its explanate pronotum and carinate elytra. Additionally, it shares with *Derodontus* the relatively narrow pronotum and densely punctate postgena, prosternum, and meso- and metaventrite. Molecular-based phylogenetic analyses of three genera of Derodontidae by McKenna et al. (2015) indicated *Laricobius* as sister to *Derodontus + Nothoderodontus*. As such, *J. sinica* with toothed, explanate pronotum and carinate elytra is a crown derodontid.
***Minimum age***: 165 Ma.
***Soft maximum age***: 251.878 Ma.
***Age justification***: *Juropeltastica sinica* was recovered from the Daohugou beds of the Haifanggou Formation, Inner Mongolia, from northeastern China (Huang 2016). The precise age of the Daohugou beds has been controversial (He et al. 2004; Gao and Ren, 2006; Wang et al. 2005; Zhang 2015). Ar/Ar and SHRIMP U-Pb dating indicated that the age of the intermediate-acid volcanic rock overlaying the Daohugou fossiliferous beds is approximately 164–165 Ma, and so the age of the fossil deposit is older than or equal to 165 Ma, corresponding to the Callovian of the Middle Jurassic (Chen et al. 2004), an age widely adopted by palaeoentomologists (Gao and Ren, 2006). Indeed, the fossil insects from the Daohugou beds may be correlated to those from the slightly younger deposit in Karatau (Kazakhstan) (Zhang 2015), but there is strong evidence indicating that the age of the Daohugou beds may be older (Huang 2016). As such, the most conservative age of the deposit is 165 Ma, providing a minimum age for *Juropeltastica*. The soft maximum constraint, 251.878 Ma, is based on the oldest occurrence of the suborder Adephaga from Late Permian deposits of the Anakit area in Middle Siberia as discussed in node 4.

#### **Node 8:** Clambidae – Eucinetidae: **145 Ma – 235 Ma.**

***Fossil taxon and specimen***: *Mesocinetus aequalis* Kirejtshuk and Ponomarenko, 2010 [holotype, PIN 4270–48, with part and counterpart; paratypes, PIN 4270–1079; PIN 4270–1071, PIN 4270–1078: Borissiak Paleontological Institute, Russian Academy of Sciences, Moscow, Russia] from the Upper Jurassic of Shar-Teg, southwestern Mongolia (Kirejtshuk and Ponomarenko 2010).
***Phylogenetic justification***: The extinct genus *Mesocinetus*, along with four other extinct genera, was placed in the extinct family Mesocinetidae, which apparently has close affinity to the extant families Scirtidae and Eucinetidae (Kirejtshuk and Ponomarenko 2010). In particular, *Mesocinetus* strongly resembles extant genera of Eucinetidae, namely *Eucinetus*, in having a characteristic body form with a strongly declined head, and 5-segmented tarsi. The close morphological resemblance indicates that *Mesocinetus* is probably a member of the extant Eucinetidae, rather than Mesocinetidae. Although the phylogenetic position of *Mesocinetus* has not been formally tested, their apomorphies (e.g., declined head and long hind legs) indicate that the genus is a crown group of the Clambidae + Eucinetidae clade.
***Minimum age***: 145 Ma.
***Soft maximum age***: 235 Ma.

***Age justification***: *Mesocinetus aequalis* was discovered in the Shar Teg Lagerstätte in southwestern Mongolia. The deposit has not been dated geochemically but its age has been inferred from comparisons of insect assemblages among Middle-Late Jurassic entomofaunas, such as those from Karatau (Late Jurassic) and Daohugou (late Middle Jurassic) (Ponomarenko et al. 2014). The minimum age constraint on the node is thus given by the boundary between the Jurassic and the Cretaceous, *ca*. 145 Ma. The soft maximum age (*ca*. 235 Ma) is derived from the age of the Madygen Formation at Dzhailyau-Cho, Kyrgyzstan. The entomofauna of the Triassic Madygen fossil locality is dominated by beetles, which constitute one quarter of total insects collected from the Madygen Formation. Among beetles, about 90% of specimens belong to Archostemata, and polyphagans are rare, including the well-known stem group polyphagan beetles belonging to the genus *Peltosyne* Ponomarenko (Ponomarenko 1977; Yan et al. 2018d). *Peltosyne triassica* and *P. varyvrosa* are is the earliest-occurring beetles placed in the polyphagan stem group based on specific apomorphies and plesiomorphic features compared to extant lineages.

#### **Node 9:** Rhipiceridae – Dascillidae: **165 Ma – 235 Ma.**

***Fossil taxon and specimen***: *Parelateriformius communis* Yan and Wang, 2010 [holotype, NIGP, nos. 151537a, 151537b (part and counterpart of almost completely preserved beetle; legs missing, antennae partly missing): Nanjing Institute of Geology and Palaeontology, Chinese Academy of Sciences, Nanjing, China] from the Haifanggou Formation at Daohugou, Ningcheng County, Inner Mongolia, northeastern China (Yan and Wang 2010).
***Phylogenetic justification***: The genus *Parelateriformius* has been assigned to the extinct polyphagan family Lasiosynidae, a group that apparently shares similarities with Dascillidae. While the most obvious apomorphies of Dascillidae, namely the strongly reduced lacinia and male flabellate antennae (Jin et al. 2013a), are not preserved in the fossil, it shows other important similarities with *Dascillus* (Jin et al. 2013b): presence of a distinct frontoclypeal suture, head lacking a posterior constriction, eyes large and somewhat protuberant, exposed antennal insertions; antenna 11-segmented and serrate, pronotum with a complete lateral carina, prosternum in front of procoxae long, prosternal process complete between coxae, procoxal cavities transverse and with lateral notches exposing protrochantins, mesocoxal cavities narrowly separated, metaventrite with discrimen and katepisternal sutures almost complete; hind coxae transverse with large coxal plates, 5-segmented tarsi, simple claws lacking an empodium, and abdomen with 5 ventrites. This would place it within crown Dascilloidea.
***Minimum age***: 165 Ma.
***Soft maximum age***: 235 Ma.
***Age justification***: For the minimum age justification of the Daohugou biota, see node 7. The soft maximum age (*ca*. 235 Ma) is derived from the Madygen Formation at Dzhailyau-Cho, Kyrgyzstan. See node 8 for justification.

#### **Node 10:** Buprestidae – Byrrhidae: **165 Ma – 235 Ma.**

***Fossil taxon and specimen***: *Sinoparathyrea bimaculata* Pan, Chang and Ren, 2011 [holotype, CNU-COL-NN2010408 (exoskeleton with legs, elytra, and wings): Key Lab of Insect Evolution and Environmental Changes, Capital Normal University, Beijing, China] from Daohugou Village, Middle Jurassic Jiulongshan Formation, Ningcheng County, Inner Mongolia, China (Pan et al. 2011).
***Phylogenetic justification***: *Sinoparathyrea bimaculata* was described as a member of Buprestidae and placed into the extinct subfamily Parathyreinae (Pan et al. 2011). Cai et al. (2015a), however, pointed out that the characters used to define Parathyreinae (Alekseev 1993), namely the straight katepisternal suture, may be a plesiomorphy. In fact, *Sinoparathyrea* shares a number of characters with members of Schizopodidae, a group that was included as a subfamily within Buprestidae by some authors. These include the relatively short prosternum in front of the procoxae, short and broad prosternal process, and a relatively large radial cell of the hind wing (Cai et al. 2015a). This indicates that *S. bimaculate* belongs to the crown group of Buprestidae – Byrrhidae.
***Minimum age***: 165 Ma.

***Soft maximum age***: 235 Ma.
***Age justification***: See node 7 for the minimum age justification of the Daohugou locality. The soft maximum is from the Madygen Formation at the Dzhailyau-Cho locality, Lyailyakskii District, Batken Region, southwest Kyrgyzstan, Central Asia, as justified in node 7.

#### **Node 11:** Dryopoidea: **112.6 Ma – 165 Ma.**
***Fossil taxon and specimen***: undescribed taxon of the series family Dryopidae [AMNH SA43296: American Museum of Natural History, New York, USA] from the Lower Cretaceous (Aptian) Crato Formation, northeastern Brazil (Grimaldi and Engel 2005).
***Phylogenetic justification***: The specimen (Fig. 10.35 in Grimaldi and Engel 2005), together with other undescribed material from the deposit, can be placed into the family Dryopidae based on its characteristic habitus, 5-segmented tarsi without lobes, and peculiar antennae with the basal segments expanded.
***Minimum age***: 112.6 Ma.
***Soft maximum age***: 165 Ma.
***Age justification***: The oldest fossil dryopid was recovered from the Crato Formation in the Araripe Basin, northeastern Brazil. This unit is generally agreed to date back to around the Aptian/Albian border (Martill et al. 2007). The Crato Formation has been dated using palynomorphs (Pons et al. 1990) to the Aptian. The upper boundary of the Aptian, at 113.0 Ma ± 0.4 Ma, gives a minimum date of 112.6 Ma. The maximum age constraint is based on the Daohugou biota, which lacks representatives of the clade. See node 7 for justification.

#### **Node 12:** Crown Psephenidae: **98.17 Ma – 165 Ma.**
***Fossil taxon and specimen***: *Vetujinbrianax cretaceus* Cai and Huang, 2018 [holotype, NIGP168215 (amber inclusion, male): Nanjing Institute of Geology and Palaeontology, Chinese Academy of Sciences, Nanjing, China] lowermost Cenomanian, from an amber mine located near Noije Bum Village, Tanaing Town, Hukawng Valley, northern Myanmar (Cai and Huang 2018).
***Phylogenetic justification***: *Vetujinbrianax cretaceus* is known from a well-preserved adult from mid-Cretaceous Burmese amber. It can be placed into the family on the basis of the following characters: body apparently soft, antennae pectinate in the males, protrochantins exposed, maxillary palps relatively long, and tarsi simple and 5-segmented. *Vetujinbrianax* can be placed in the extant subfamily Eubrianacinae based on the presence of plesiomorphic characters such as short prosternal process and paired tibial apical spurs, similar to the extant Oriental genus *Jinbrianax* (Lee et al. 2007; Cai and Huang 2018). *V. cretaceus* represents the oldest fossil record of Eubrianacinae, as well as the first definitive adult fossil of Psephenidae from the Mesozoic. Further specimens of psephenid larvae, also belonging to the subfamily Eubrianacinae from Burmese amber were reported by Bao et al. (2018) and described as *Succinumanax birmaniasis*. They represent the oldest fossil record of Psephenidae, and show a high similarity in morphology with fossils from the Eocene Messel pit, Germany. A phylogenetic analysis demonstrated that *S. birmaniasis* and the Messel specimen can be placed in Psephenidae.
***Minimum age***: 98.17 Ma.
***Soft maximum age***: 165 Ma.
***Age justification***: The specimen is preserved in Burmese amber (burmite) mined in the Hukawng Valley, Kachin State, northern Myanmar. Volcanoclastic matrix from the amber-bearing horizon was dated radiometrically at 98.79 ± 0.62 Ma (Shi et al. 2012), although this estimate should be taken as a minimum age (Mao et al. 2018). A mid-Cretaceous age is consistent with paleontological evidence (Grimaldi et al. 2002). The discovery of an ammonite in the amber indicates that it is at most Albian in age (Yu et al., 2019). Here we use 98.17 Ma as the conservative minimum age of Burmese amber from northern Myanmar. The maximum age constraint is based on the Daohugou biota, which lacks representatives of the clade. See node 7 for justification.

#### **Node 13:** Elmidae – (Limnichidae + Heteroceridae): **105 Ma – 165 Ma.**
***Fossil taxon and specimen***: *Elmadulescens rugosus* Peris, Maier and Sánchez-García, 2015 [holotype, CES–

567 (amber inclusion): Institut de Recerca de la Biodiversitat (IRBio), Facultat de Geologia, Universitat de Barcelona, Martí i Franquès s/n, 08028 Barcelona, Spain] El Soplao amber outcrop located of the El Soplao Cave, Celis, Cantabria, Spain (Peris et al. 2015a).
***Phylogenetic justification***: *Elmadulescens rugosus* can be placed into Elmidae on the basis of its long, slender and widely inserted antennae, large and protuberant eyes, a pentagonal scutellum abruptly elevated anteriorly, long legs with elongate and simple 5-segmented tarsi, metacoxae with a posterior face, pronotum with complete lateral carinae, the last tarsomere longer than the other four combined, large pretarsal claws, and five abdominal ventrites (Peris et al. 2015a). The holotype is preserved in a transparent amber piece, but is dorsal-ventrally compressed and deformed, making the observation of some characters difficult.
***Minimum age***: 105 Ma
***Soft maximum age***: 165 Ma
***Age justification***: The amber piece was found in Early Albian strata of the Las Peñosas Formation, as determined by biostratigraphy (Najarro et al. 2009). The minimum age is derived from the upper Albian. The maximum age constraint is based on the Daohugou biota, which lacks representatives of the clade. See node 7 for justification.

#### **Node 14:** Crown Heteroceridae: **122.2 Ma – 165 Ma.**
***Fossil taxon and specimen***: *Heterocerites magnus* Prokin and Ren, 2011 [holotype, CNU-COL-LB-2010023 (impression of adult beetle with party preserved legs and mandibles): Key Lab of Insect Evolution and Environmental Changes, Capital Normal University, Beijing, China] from the Huanbanjigou of the Yixian Formation, Shangyuan, Beipiao city, Liaoning Province (Prokin and Ren 2011).
***Phylogenetic justification***: The fossil species from the Yixian Formation was placed in the formal genus *Heterocerites* erected by Ponomarenko (1986) for compression fossils of putative heterocerids. While the genus is probably not monophyletic (Li et al. 2020a), *H. magnus* can be placed into Heteroceridae on the basis of having a densely pubescent body, large head with remnants of prominent mandibles and labrum, characteristically modified antennae with the distal half forming a loosely serrated club, a series of spine at the outer edges of the tibiae, short mesosternum, and stridulate ridges on ventrite 1 (Clarke 1973; Vanin et al. 2016). The dark elytra moreover possess tree pairs of light spots, a pattern similar to those found in extant members of the family (Vanin et al. 2016).
***Minimum age***: 122.2 Ma.
***Soft maximum age***: 165 Ma.
***Age justification***: Diverse and abundant fossils of the terrestrial Jehol Biota have been discovered from the Yixian Formation and the overlying Jiufotang Formation in Inner Mongolia, Hebei Province and Liaoning Province, northeastern China. The Yixian Formation is Early Cretaceous in age. $^{40}Ar/^{39}Ar$ dating produced a mean age of 124.6 ± 0.1 Ma for sanidine from one tuff interbedded in the fossiliferous horizons of the lower Yixian Formation near Jianshangou village, and 40Ar/39Ar single- grain total fusion analyses provided an age of 124.6 ± 0.25 Ma for the same tuff from Sihetun village (Swisher et al. 1999). Chang et al. (2009) provided a high precision $^{40}Ar/^{39}Ar$ dating of the Yixian Formation: the lowest part of the main fossil bed was dated as 124.1± 0.3 Ma, while the uppermost part of the bed was dated as 122.9 ± 0.7 Ma. The latter age provides a conservative minimum constraint on the node, 122.2 Ma. The maximum age constraint is based on the Daohugou biota, which lacks representatives of the clade. See node 7 for justification.

#### **Node 15:** Artematopodidae – Omethidae: **165 Ma – 235 Ma.**
***Fossil taxon and specimen***: *Sinobrevipogon jurassicus* Cai, Lawrence, Ślipiński and Huang, 2015 [holotype, NIGP160704; paratypes, NIGP160705, NIGP160706 (holotype nearly completely preserved adult beetle, with both dorsal and ventral characters visible, including hindwings); paratype NIGP160705 (displays mainly ventral and some dorsal characters); paratype NIGP160706 (shows mainly dorsal and a few ventral characters such as abdominal ventrites): Nanjing Institute of Geology and Palaeontology, Chinese Academy of Sciences, Nanjing, China] from the Haifanggou Formation at Daohugou, Ningcheng County, Inner Mongolia, northeastern China (Cai et al. 2015b).
***Phylogenetic justification***: *Sinobrevipogon jurassicus* exhibits a number of defining features of

Artematopodidae, including the presence of paired carinae on the prosternum and an internal apical interlocking tongue on the ventral side of each elytron. However, it differs from modern members of Artematopodidae by having the mesocoxal cavities closed by the mesepimeron and the anterolateral edge of metanepisternum. A morphological phylogenetic analysis nonetheless recovered *Sinobrevipogon* within crown Artematopodidae (Cai et al. 2015b; 2020).
***Minimum age***: 165 Ma.
***Soft maximum age***: 235 Ma.
***Age justification***: See node 7 for minimum age justification. The soft maximum age (ca. 235 Ma) is derived from the Madygen Formation at Dzhailyau-Cho, Kyrgyzstan. See node 8 for justification.

### **Node 16:** Throscidae – Eucnemidae: **125 Ma – 235 Ma.**

***Fossil taxon and specimen***: *Potergosoma gratiosum* Kovalev and Kirejtshuk, 2013 [holotype, "750" (amber inclusion, probably male): Lebanese University, Faculty of Sciences II, Fanar, Lebanon] from the Early Cretaceous Lebanese amber originating from the outcrop Hammana – Mdeirij, Caza Baabda, Mouhafazet, Jabal Loubnan, central Lebanon (Kovalev et al. 2013; Li et al. 2020b).
***Phylogenetic justification***: *Potergosoma gratiosum* possesses clear diagnostic characters of the Throscidae, some regarded as apomorphies of the family. These are the developed metacoxal plates, exposed and heavily sclerotised labrum, antennae with distinctly enlarged apical antennomeres forming an elongate antennal club, characteristic grooves on the underside for receiving antennae and legs, wide prosternal process, and rather widely separated mesocoxae. The genus also possesses several plesiomorphies: complete lateral pronotal carinae, absence of tarsal grooves, simple tarsomere 4, simple tibiae, poorly developed pro-mesothoracic leg impressions, antennal fossae and distinct serial elytral punctures (Muona 2019). Morphological phylogenetic analyses recovered *Potergosoma* as a member of stem Throscidae (Muona 2019; Li et al. 2020b), making it suitable for calibrating the split between Throscidae and Eucnemidae.
***Minimum age***: 125 Ma.
***Soft maximum age***: 235 Ma.
***Age justification***: Krzemiski et al. (2014) indicated a Barremian age for the amber-bearing outcrop at Hammana – Mdeirij in central Lebanon. A recent biostratigraphic study of Lebanese amber by Maksoud et al. (2017) indicated that the pieces of amber with bioinclusions from the middle and upper intervals may have been reworked from the lower interval, so the age of the amber from "Grès du Liban" could be early Barremian or even older in age. As such, the minimum age for *E. rugidorsum* can be taken from the top of the Barremian, which is currently estimated at 125.0 Ma (International Chronostratigraphic Chart, v. 2018/08), representing the minimum age constraint for this node. The soft maximum age (*ca*. 235 Ma) is derived from the Madygen Formation at Dzhailyau-Cho, Kyrgyzstan. See node 8 for justification.

### **Node 17:** *Osslimus* (Elateridae) **–** Lycidae: **98.17 Ma – 165 Ma.**

***Fossil taxon and specimen***: *Burmolycus compactus* Bocak, Li and Ellenberger, 2019 [holotype, MB.I 7319 (amber inclusion, female): Museum für Naturkunde, Berlin, Germany] lowermost Cenomanian, from an amber mine located near Noije Bum Village, Tanaing Town, Hukawng Valley, northern Myanmar (Bocak et al. 2019).
***Phylogenetic justification***: *Burmolycus compactus* can be placed into the family Lycidae on the basis of having 11-segmented antennae, a ridged pronotum, a very short and transverse prosternum, 5-segmented tarsi, and tarsomeres 2–4 lobed (Bocak and Bocakova 2008, 2010). It can further be assigned to the extant subfamily Dexorinae based on the presence of a prolonged cranium, frontally inserted antennae, antennomere 2 relatively long compared to the antennomere 3, hypognathous head, rounded mouthpart cavity, reduced labium with 1 or 2-segmented labial palpi, and femora obliquely attached to trochanters (Bocak et al. 2019). Other lycids belonging to extant and extinct taxa are known from Burmese amber, making them likewise suitable candidates for calibration of the node (Tihelka et al. 2019; Kazantsev 2019).
***Minimum age***: 98.17 Ma.
***Soft maximum age***: 165 Ma.
***Age justification***: See node 12 for minimum age justification. The maximum age constraint is based on the

Daohugou biota, which lacks representatives of the clade. See node 7 for justification.

**Node 18:** Lampyridae – Phengodidae + Rhagophthalmidae: **98.17 Ma – 165 Ma.**
***Fossil taxon and specimen***: *Protoluciola albertalleni* Kazantsev, 2015 [holotype, ICM-BU-1687–1 (amber inclusion, male): Insect Centre, Moscow, Russia] lowermost Cenomanian, from an amber mine located near Noije Bum Village, Tanaing Town, Hukawng Valley, northern Myanmar (Kazantsev 2015).
***Phylogenetic justification***: *Protoluciola albertalleni* can be placed into the extant lampyrid subfamily Luciolinae based on the presence of six ventrites, genital segment apparently absent, the presence of a photic organ on the abdomen, 11-segmented antennae inserted anteriorly to the eyes, and five-segmented tarsi with simple claws (McDermott 1964).
***Minimum age***: 98.17 Ma.
***Soft maximum age***: 165 Ma.
***Age justification***: See node 12 for minimum age justification. The maximum age constraint is based on the Daohugou biota, which lacks representatives of the clade. See node 7 for justification.

**Node 19:** Crown Cantharidae: **105 Ma – 165 Ma.**
***Fossil taxon and specimen***: *Molliberus albae* Peris and Fanti, 2018 [holotype, CES-522 (amber inclusion): laboratory of the El Soplao Cave, Celis-Rábago, Cantabria, Spain) from Early Cretaceous Spanish amber originating from the El Soplao deposit, Celis municipality, Cantabria, Spain (Peris and Fanti 2018).
***Phylogenetic justification***: *Molliberus albae* can be placed into the extant subfamily Cantharinae based on the maxillary palpomeres different in length to the last securiform segment, long elytra, symmetrical last abdominal segments, prothorax without lateral processes, the presence of tibial spurs, and the fourth tarsomere undivided (Peris and Fanti 2018). As such, it represents the earliest member of crown Cantharidae.
***Minimum age***: 105 Ma.
***Soft maximum age***: 165 Ma.
***Age justification***: See node 13 for minimum age justification. The maximum age constraint is based on the Daohugou biota, which lacks representatives of the clade. See node 7 for justification.

**Node 20:** Georissidae – Hydrophilidae: **147.28 Ma – 235 Ma.**
***Fossil taxon and specimen***: *Protochares brevipalpis* Fikáček, Prokin, Yan, Yue, Wang, Ren and Beattie, 2014 [holotype, AMF109568 (exoskeleton preserved in dorsal aspect, with fragments of legs): Department of Earth Sciences, Australian Museum, Sydney, Australia] from Talbragar fossil fish bed, *ca*. 14 km NNW of Ulan, New South Wales, Australia (Fikáček et al. 2014).
***Phylogenetic justification***: *Protochares brevipalpis* can be assigned to Hydrophilidae based on the unique combination of the following characters: labrum well-sclerotised and largely exposed, clypeus large and with distinct frontoclypeal sutures reaching the lateral margin of the head closely before eyes, scutellum triangular, and head not strongly constricted behind the eyes (Fikáček et al. 2012, 2014).
***Minimum age***: 147.28 Ma.
***Soft maximum age***: 235 Ma.
***Age justification***: Dating the Jurassic Talbragar Lagerstätten fish beds has traditionally been contentious because no palynofloras have been recovered from the beds themselves as the rocks are highly oxidised. SHRIMP dating indicated an age of 151.55 ± 4.27 Ma (Bean 2006). This provides a conservative minimum age constraint on the node. The soft maximum age (*ca*. 235 Ma) is derived from the Madygen Formation at Dzhailyau-Cho, Kyrgyzstan. See node 8 for justification.

**Node 21:** Trogidae – Lucanidae: **165 Ma – 235 Ma.**
***Fossil taxon and specimen***: *Juraesalus atavus* Nikolajev, Wang, Liu and Zhang, 2011 [holotype, NIGP 152456 (exoskeleton with legs, preserved in ventral aspect): Nanjing Institute of Geology and Palaeontology, Chinese Academy of Sciences, Nanjing, China] from the Haifanggou Formation at Daohugou, Ningcheng County, Inner Mongolia, northeastern China (Nikolajev et al. 2011).

***Phylogenetic justification***: *Juraesalus atavus* can be placed into Lucanidae based on the presence of well-developed and somewhat projecting antennae, a complete prosternal process, concealed trochantins, procoxal cavities strongly transverse, closed externally, open internally, mesocoxal cavities narrowly separated and laterally open, 5-segmented tarsi, and a rather elongate and subglabrous body (Holloway 1960, 1969). Nikolajev et al. (2011) considered the fossil to represent a stem group to the extant subfamily Aesalinae based on the presence of a 3-segmented non-lamellate club, mandibles produced beyond apex of clypeus, eyes not divided by canthus, and abdomen with 5 ventrites. However, Kim and Farrell (2015) pointed out that these characters are not unique to Aesalinae and instead considered *J. atavus* to be the oldest member of Lucanidae *s.s.* In either way, the fossil would be suitable for calibrating the split between Trogidae and Lucanidae.
***Minimum age***: 165 Ma.
***Soft maximum age***: 235 Ma.
***Age justification***: For the minimum age justification of the Daohugou biota, see node 7. The soft maximum age (*ca*. 235 Ma) is derived from the Madygen Formation at Dzhailyau-Cho, Kyrgyzstan. See node 8 for justification.

**Node 22:** Passalidae – Geotrupidae: **122.2 Ma – 165 Ma.**
***Fossil taxon and specimen***: *Parageotrupes incanus* Nikolajev and Ren, 2010 [holotype, CNU-COL-LB2009751 (exoskeleton with partly preserved legs and antennae): Key Lab of Insect Evolution and Environmental Changes, Capital Normal University, Beijing, China] from the Huanbanjigou of the Yixian Formation, Shangyuan, Beipiao city, Liaoning Province (Nikolajev and Ren 2010).
***Phylogenetic justification***: *Parageotrupes incanus* can be placed into Geotrupidae on the basis of the size and the oval shape of the body, mandibles not hidden by a clypeus, eyes divided by canthus, pygidium hidden by elytra. The clypeus bears a tubercle and is clearly delimited from the vertex, suggesting that the species belongs to the extant subfamily Geotrupinae (Nikolajev and Ren 2010).
***Minimum age***: 122.2 Ma.
***Soft maximum age***: 165 Ma.
***Age justification***: For the minimum age justification of the Yixian Formation, see node 14. The maximum age constraint is based on the Daohugou biota, which lacks representatives of the clade. See node 7 for justification.

**Node 23:** Hybosoridae – Scarabaeidae: **155 Ma – 235 Ma.**
***Fossil taxon and specimen***: *Protohybosorus grandissimus* Nikolajev, 2010 [holotype, PIN no. 2335/25 (exoskeleton with fragments of legs and antennae): from Karatau–Mikhailovka, Karabastau Formation of the Karatau Range near the village of Mikhailovka, Chayan District, Chimkent Region, southern Kazakhstan (Nikolajev 2010)
***Phylogenetic justification:*** The genus *Protohybosorus*, containing three species, can be assigned to Hybosoridae based on the small size of the species, their developed eyes, labrum and mandibles produced beyond the apex of the clypeus, trapezoidal and convex pronotum with base wider than the elytral base, elytra convex and with well-defined striae, protibia with three teeth, mesotibia and metatibia somewhat dilated at the apex, mesotibia with 2 subequal spurs, mesotibia and metatibia without a transverse carina, and abdomen with five visible sternites (Jameson 2012; Ocampo 2006; Yan et al. 2012).
***Minimum age***: 155 Ma.
***Soft maximum age***: 235 Ma.
***Age justification***: For the minimum age justification of Karatau, see node 5. The soft maximum age (*ca*. 235 Ma) is derived from the Madygen Formation at Dzhailyau-Cho, Kyrgyzstan. See node 8 for justification.

**Node 24:** Ptiliidae – Hydraenidae: **182 Ma – 235 Ma.**
***Fossil taxon and specimen***: *Ochtebiites minor* Ponomarenko, 1985 [holotype, PIN 2245/278 (exoskeleton in dorsal and ventral aspect, with fragments of legs): Borissiak Paleontological Institute, Russian Academy of Sciences, Moscow, Russia] from trench 82an, Lyagush'ye locality, Kemerovo, Abasheva Formation, Kemerovo

Oblast, Russia (Ponomarenko 1985).
***Phylogenetic justification***: The oldest known representative of the family Hydraenidae is *Ochtebiites minor*. It can be placed in the family based on its small size, elongate body, elongate maxillary palpi, and the presence of an intercoxal sternite between the metacoxae (Jäch et al. 2016).
***Minimum age***: 182 Ma.
***Soft maximum age***: 235 Ma.
***Age justification***: The Abasheva Formation appears to be Pliensbachian in age, based on the fossil flora and fauna (Kovalev 1981; Mogutcheva 2009). The top of the Pliensbachian is 182.7 ± 0.7 (International Chronostratigraphic Chart, v. 2018/08), giving a minimum constraint of 182 Ma. The soft maximum age (*ca*. 235 Ma) is derived from the Madygen Formation at Dzhailyau-Cho, Kyrgyzstan. See node 8 for justification.

#### **Node 25:** Agyrtidae – Leiodidae: **165 Ma – 235 Ma.**
***Fossil taxon and specimen***: *Mesecanus communis* Ponomarenko, 1977 [holotype, PIN 3000/926 (exoskeleton with preserved appendages in dorsal and ventral aspects); paratypes, PIN 3000/915, 3000/913, 3000/909, 2744/12: Borissiak Paleontological Institute, Russian Academy of Sciences, Moscow, Russia], from Novospasskoye village, Ichetuy Formation, Buryat, Russia (Ponomarenko, 1977; Perkovsky 1999).
***Phylogenetic justification***: The systematic placement of *Mesecanus communis* has been historically quite unstable. Ponomarenko (1977) placed the species into Agyrtidae, while Perkovsky (1999) assigned the fossil to Leiodidae. It was treated as an agyrtid again by Newton (1997). It nonetheless meets all the diagnostic features of Agyrtidae, and we consider it as a member of this group. It has been compared to the extant genus *Ecanus* by Newton (1997)
***Minimum age***: 165 Ma.
***Soft maximum age***: 235 Ma.
***Age justification***: The Novospasskoye locality is usually considered Upper to Middle Jurassic based on biostratigraphy (Ponomarenko 1993). We thus use 166.1 ±1.2 as a conservative constraint on the node, representing the upper boundary of the Callovian, the last stage of the Middle Jurassic (International Chronostratigraphic Chart, v. 2018/08). The soft maximum age (*ca*. 235 Ma) is derived from the Madygen Formation at Dzhailyau-Cho, Kyrgyzstan. See node 8 for justification.

#### **Node 26:** Crown Staphylinidae: **165 Ma – 235 Ma.**
***Fossil taxon and specimen***: undescribed member of Silphidae [NIGP 156145a/b (exoskeleton with preserved appendages): Nanjing Institute of Geology and Palaeontology, Chinese Academy of Sciences, Nanjing, China] from the Haifanggou Formation at Daohugou, Ningcheng County, Inner Mongolia, northeastern China (Cai et al. 2014b).
***Phylogenetic justification***: The undescribed specimens can be placed into the former family Silphidae, treated here as a subfamily of Staphylinidae *sensu nov.*, based on their general habitus, clubbed antennae, large mesoscutellum, truncate elytra, and well-separated mesocoxae (Cai et al. 2014b). Given that it has been recently demonstrated that the enigmatic *Leehermania prorova* from the Late Triassic Cow Branch Formation on the Virginia-North Carolina border, USA is in fact not a staphylinid as assumed previously, these burying beetles are the oldest candidates for calibrating the node representing crown Staphylinidae (Chatzimanolis et al. 2012; Fikáček et al. 2020).
***Minimum age***: 165 Ma.
***Soft maximum age***: 235 Ma.
***Age justification***: For the minimum age justification of the Daohugou biota, see node 7. The soft maximum age (*ca*. 235 Ma) is derived from the Madygen Formation at Dzhailyau-Cho, Kyrgyzstan. See node 8 for justification.

#### **Node 27:** *Megalopaederus* (Paederinae) – Staphylininae: **122.2 Ma – 165 Ma.**
***Fossil taxon and specimen***: *Megolisthaerus minor* Cai and Huang, 2013 [holotype, NIGP 153697 (amber inclusion): Nanjing Institute of Geology and Palaeontology, Chinese Academy of Sciences, Nanjing, China] from the Huanbanjigou of the Yixian Formation, Shangyuan, Beipiao city, Liaoning Province (Cai and Huang

2013).
***Phylogenetic justification***: *Megolisthaerus minor* can be placed into Staphylininae on the basis of its relatively large and elongate body, antennae inserted along the anterior margin of the head, pronotal hypomeron lacking postcoxal processes; pro- and mesotarsi 5-segmented, elytron without epipleural keel, and two pairs of paratergites on abdominal segment III–VI (Cai and Huang 2013).
***Minimum age***: 122.2 Ma.
***Soft maximum age***: 165 Ma.
***Age justification***: For the minimum age justification of the Yixian Formation, see node 14. The maximum age constraint is based on the Daohugou biota, which lacks representatives of the clade. See node 7 for justification.

#### **Node 28:** Silphinae – Staphylinidae *partim*: **165 Ma – 235 Ma.**
***Fossil taxon and specimen***: As for node 25.
***Minimum age***: 165 Ma.
***Soft maximum age***: 235 Ma.
***Age justification***: As for node 25.

#### **Node 29:** Staphylinidae: Apateticinae – Osoriinae: **98.17 Ma – 165 Ma.**
***Fossil taxon and specimen***: *Mesallotrochus longiantennatus* Cai and Huang, 2015 [holotype, NIGP157737 (amber inclusion): Nanjing Institute of Geology and Palaeontology, Chinese Academy of Sciences, Nanjing, China] lowermost Cenomanian, from an amber mine located near Noije Bum Village, Tanaing Town, Hukawng Valley, northern Myanmar (Cai and Huang 2015)
***Phylogenetic justification***: *Mesallotrochus* is placed in the extant osoriine tribe Thoracophorini based on its general habitus, the protibia with the inner edge straight, missing ctenidium, exposed protrochantins, open procoxal cavities, and a more or less flattened body (Cai and Huang 2015). As such, *M. longiantennatus* represents the oldest member of Osoriinae and is suitable for calibrating the split between Apateticinae and Osoriinae.
***Minimum age***: 98.17 Ma.
***Soft maximum age***: 165 Ma.
***Age justification***: For the minimum age justification of Burmese amber from northern Myanmar, see node 12. The maximum age constraint is based on the Daohugou biota, which lacks representatives of the clade. See node 7 for justification.

#### **Node 30:** Crown Bostrichoidea: **165 Ma – 235 Ma.**
***Fossil taxon and specimen***: *Paradermestes jurassicus* Deng, Ślipiński, Ren and Pang, 2017 [CNU-COL-NN2017001 (exoskeleton) with legs and antennae: Key Lab of Insect Evolution and Environmental Changes, Capital Normal University, Beijing, China] from Daohugou Village, Middle Jurassic Jiulongshan Formation, Ningcheng County, Inner Mongolia, China (Deng et al. 2017a).
***Phylogenetic justification***: *Paradermestes* can be placed into Dermestidae on the basis of having a small deflexed head inserted into the prothorax, a developed antennal club, posterior angles of pronotum projecting posteriorly, excavate metacoxae, and simple tarsomeres. Given its age, it is a suitable calibration point for setting the minimum age of Bostrichoidea (Kirejshuk and Azar 2009).
***Minimum age***: 165 Ma.
***Soft maximum age***: 235 Ma.
***Age justification***: For the minimum age justification of the Daohugou biota, see node 7. The soft maximum age (*ca*. 235 Ma) is derived from the Madygen Formation at Dzhailyau-Cho, Kyrgyzstan. See node 8 for justification.

#### **Node 31:** Crown Bostrichidae: **98.17 Ma – 165 Ma.**
***Fossil taxon and specimen***: *Poinarinius burmaensis* Legalov, 2018 [holotype ISEA no. MA2018/3 (amber inclusion): Institute of Systematics and Ecology of Animals, Novosibirsk, Russia] lowermost Cenomanian, from

an amber mine located near Noije Bum Village, Tanaing Town, Hukawng Valley, northern Myanmar (Legalov 2018)
***Phylogenetic justification***: *Poinarinius* can be placed into Bostrichidae based on its non- pseudotetramerous tarsi, head lacking paired ocelli, mesocoxal cavities closed laterally, short and triangular metatrochanter, and the femur partially separated from the coxa. Based on the protibia with one apical spine and subequal tarsomeres 1 and 2, the fossil was assigned into the extant subfamily Dinoderinae (Legalov 2018).
***Minimum age***: 98.17 Ma.
***Soft maximum age***: 165 Ma.
***Age justification***: For the minimum age justification of Burmese amber from northern Myanmar, see node 12. The maximum age constraint is based on the Daohugou biota, which lacks representatives of the clade. See node 7 for justification.

**Node 32:** Crown Ptinidae: **105 Ma – 165 Ma.**
***Fossil taxon and specimen***: *Actenobius magneoculus* Peris, Philips, and Delclòs, 2015 [holotype, SJ-10-18, SJ (amber inclusion): San Just amber collection, Fundación Conjunto Paleontológico de Teruel-Dinópolis, Teruel, Spain] from the San Just outcrop, Utrillas, Regachuelo Member, Escucha Formation, Teruel Province, Spain (Peris et al. 2015b).
***Phylogenetic justification***: *Actenobius magneoculus* is placed in the family Ptinidae by its cylindrical body, deflexed head inserted into the prothorax and covered dorsally by the pronotum; antennae with 11 antennomeres and with the last three segments asymmetrically elongated, large compound eyes, reduced prosternum lengthwise and pronotum convex dorsally, ovoid pro- and mesocoxae, transverse and excavated hind coxae for reception of metafemora, slender femora and tibiae, tarsal formula 5-5-5, and tarsomere 1 longer than tarsomere 5 (Lawrence and Britton 1991; Philips and Bell 2010). It can be assigned to Anobiinae based on the metaventrite and first abdominal segment without grooves for the reception of hind legs, complete lateral border of pronotum, and hypognathous head with mandibles not near the metaventrite with the head retracted (Español 1992)
***Minimum age***: 105 Ma.
***Soft maximum age***: 165 Ma.
***Age justification***: The amber from San Just is early to middle Albian in age based on palynomorph correlation of the Escucha Formation (Peñalver and Delclòs 2010). We thus use 105 Ma as a conservative minimum on the age of the amber deposit. The maximum age constraint is based on the Daohugou biota, which lacks representatives of the clade. See node 7 for justification.

**Node 33:** Thanerocleridae **–** Cleridae: **165 Ma – 235 Ma.**
***Fossil taxon and specimen***: *Wangweiella calloviana* Kolibáč and Huang, 2016 [holotype, NIGP162533 (exoskeleton with fragments of legs and antennae): Nanjing Institute of Geology and Palaeontology, Chinese Academy of Sciences, Nanjing, China] from Daohugou Village, Middle Jurassic Jiulongshan Formation, Ningcheng County, Inner Mongolia, China (Kolibáč and Huang 2016).
***Phylogenetic justification***: *Wangweiella calloviana* can be placed into the clerid subfamily Epiclininae based on its elongate pedicel, absence of paracoxal sutures, basitarsus shortened but not covered from above with the base of tarsomere 2, and tegmen composed of two parts. A morphological phylogenetic analysis recovered the species as sister to the rest of Epiclininae (Kolibáč and Huang 2016), making it suitable for calibrating the split between Thanerocleridae and Cleridae.
***Minimum age***: 165 Ma.
***Soft maximum age***: 235 Ma.
***Age justification***: For the minimum age justification of the Daohugou biota, see node 7. The soft maximum age (*ca*. 235 Ma) is derived from the Madygen Formation at Dzhailyau-Cho, Kyrgyzstan. See node 8 for justification.

**Node 34:** Prionoceridae **–** Melyridae: **165 Ma – 235 Ma.**
***Fossil taxon and specimen***: *Idgiaites jurassicus* Liu, Ślipiński, Leschen, Ren and Pang, 2015 [holotype: CNU-

COL-NN2010238 (nearly complete exoskeleton in lateral position): Key Lab of Insect Evolution and Environmental Changes, Capital Normal University, Beijing, China] from Daohugou Village, Middle Jurassic Jiulongshan Formation, Ningcheng County, Inner Mongolia, China (Liu et al. 2015).
***Phylogenetic justification***: *Idgiaites jurassicus* belongs to Prionoceridae based on the general body shape and the 11-segmented antenna with three apical segments flattened and the apical segment enlarged (Liu et al. 2015).
***Minimum age***: 165 Ma.
***Soft maximum age***: 235 Ma.
***Age justification***: For the minimum age justification of the Daohugou biota, see node 7. The soft maximum age (*ca*. 235 Ma) is derived from the Madygen Formation at Dzhailyau-Cho, Kyrgyzstan. See node 8 for justification.

**Node 35:** Lymexylidae: **112.6 Ma – 235 Ma.**
***Fossil taxon and specimen***: *Cratoatractocerus grimaldii* Wolf, 2011 [holotype SMNS-66534 (exoskeleton with fragments of appendages): Staatliches Museum für Naturkunde Stuttgart, Stuttgart, Germany] from the vicinity of Nova Olinda, southern Ceará, Nova Olinda Member of the Crato Formation, north-east Brazil (Wolf 2011).
***Phylogenetic justification***: *Cratoatractocerus grimaldii* can be placed into Lymexylidae and the subfamily Atractocerinae based on its long and slender body form, large eyes that occupy almost the entire frons and are narrowly separated, absence of the first radial crossvein, and heavily sclerotised female genitalia (Kurosawa 1985; Wheeler 1986; Wolf 2011).
***Minimum age***: 112.6 Ma.
***Soft maximum age***: 235 Ma.
***Age justification***: For the minimum age justification of the Crato Formation, see node 11. The soft maximum age (*ca*. 235 Ma) is derived from the Madygen Formation at Dzhailyau-Cho, Kyrgyzstan. See node 8 for justification.

**Node 36:** Crown Tenebrionoidea: **165 Ma – 235 Ma.**
***Fossil taxon and specimen***: *Archaeoripiphorus nuwa* Hsiao, Yu and Deng, 2017 [holotype: CNU-C-NN-2006841 (exoskeleton with fragments of appendages): Key Lab of Insect Evolution and Environmental Changes, Capital Normal University, Beijing, China] from Daohugou Village, Middle Jurassic Jiulongshan Formation, Ningcheng County, Inner Mongolia, China (Hsiao et al. 2017).
***Phylogenetic justification***: *Archaeoripiphorus* was considered to represent a member of Rhipiphoridae by Hsiao et al. (2017) based mainly on overall morphological similarity. Batelka et al. (2018) reviewed the evidence for the classification of the genus and concluded that it cannot be placed into Rhipiphoridae with certainty. It however clearly is a member of Tenebrionoidea, as indicated by its wedge-shaped body and tarsal formula 5-5-4, and is herein used to calibrate the node representing crown Tenebrionoidea.
***Minimum age***: 165 Ma.
***Soft maximum age***: 235 Ma.
***Age justification***: For the minimum age justification of the Daohugou biota, see node 7. The soft maximum age (*ca.* 235 Ma) is derived from the Madygen Formation at Dzhailyau-Cho, Kyrgyzstan. See node 8 for justification.

**Node 37:** Rhipiphoridae – Mordellidae: **98.17 Ma – 235 Ma.**
***Fossil taxon and specimen***: *Protoripidius burmiticus* Cai, Yin and Huang, 2018 [holotype, NIGP166155 (amber inclusion): Nanjing Institute of Geology and Palaeontology, Chinese Academy of Sciences, Nanjing, China] lowermost Cenomanian, from an amber mine located near Noije Bum Village, Tanaing Town, Hukawng Valley, northern Myanmar (Cai et al. 2018a)
***Phylogenetic justification***: *Protoripidius burmiticus* can be assigned to the rhipiphorid subfamily Ripidiinae based on the combination of the following characters: abdominal sternite II visible externally and similar to

sternite III, eyes coarsely facetted, elytra shortened in the male and widely separated, hind wings not folded transversely, tibial spurs completely absent, claws simple, and body densely clothed with short setae (Falin and Engel 2010; Cai et al. 2018a).
***Minimum age***: 98.17 Ma.
***Soft maximum age***: 235 Ma.
***Age justification***: For the minimum age justification of Burmese amber from northern Myanmar, see node 12. The maximum age constraint is based on the Daohugou biota, which lacks representatives of the clade. See node 7 for justification.

**Node 38:** Ischaliidae – Aderidae: **125 Ma – 165 Ma.**
***Fossil taxon and specimen***: nine undescribed fossils [“JG 204/15”, “96”, “1282”, “AD4A”, “1521”, “1239”, “853-F”, “NBS-2F”, “822” (amber inclusions of various degrees of preservation): Lebanese University, Faculty of Sciences II, Fanar, Lebanon] from the Nabaa Es-Sukkar – Brissa, Hammana – Mdeirij, Ain Dara, Rihane, and Jouar Es-Souss (Bkassine) outcrops, Lebanon (Kirejtshuk and Azar 2013).
***Phylogenetic justification***: Kirejtshuk and Azar (2013) cite nine members of the family Aderidae from Cretaceous Lebanese amber. The presence of aderids in Lebanese amber was also noted by Grimaldi and Engel (2005).
***Minimum age***: 125 Ma.
***Soft maximum age***: 165 Ma.
***Age justification***: For the minimum age justification of Lebanese amber, see node 16. The maximum age constraint is based on the Daohugou biota, which lacks representatives of the clade. See node 7 for justification.

**Node 39:** Salpingidae – Pythidae: **53 Ma – 165 Ma.**
***Fossil taxon and specimen***: *Eopeplus stetzenkoi* Kirejtshuk and Nel, 2009 [holotype, PA 891/3 (amber inclusion): Laboratoire de Paléontologie, Muséum National d’Histoire Naturelle, Paris, France] from Eocene French amber obtained from the Farm Le Quesnoy locality, Chevrière, region of Creil, Oise department, northern France (Kirejtshuk and Nel 2009).
***Phylogenetic justification***: *Eopeplus stetzenkoi* can be placed into the salpingid subfamily Inopeplinae based on its flattened body, pronotum distinctly narrowing posteriorly, and shortened elytra. It moreover shares with recent representatives of the group the shape of the head, structure of antennae, and others (Kirejtshuk and Nel 2009).
***Minimum age***: 53 Ma.
***Soft maximum age***: 165 Ma.
***Age justification***: The age of Oise amber, approximately 53 Ma is indicated by the presence of mammalian fossils corresponding to the MP7 horizon (Kirejtshuk and Nel 2009). The maximum age constraint is based on the Daohugou biota, which lacks representatives of the clade. See node 7 for justification.

**Node 40:** Anthicidae – Meloidae: **125 Ma – 165 Ma.**
***Fossil taxon and specimen***: *Camelomorpha longicervix* Kirejtshuk, Azar and Telnov, 2008 [holotype, “846” (amber inclusion, sex undefinable): Lebanese University, Faculty of Sciences II, Fanar, Lebanon] from the Early Cretaceous Lebanese amber originating from the outcrop Hammana – Mdeirij, Caza Baabda, Mouhafazet, Jabal Loubnan, central Lebanon (Kirejtshuk and Azar 2008).
***Phylogenetic justification:*** *Camelomorpha longicervix* can be placed into the anthicid subfamily Macratriinae based on its elongate body, absence of a frontoclypeal suture, elytra with distinct longitudinal sulcus along the lateral margins and parallel to epipleura, mesosternum and mesepisterna fused, without an apparent carina at fusion line (Chandler 2010). It is thus suitable for calibrating the split between Anthicidae and Meloidae.
***Minimum age***: 125 Ma.
***Soft maximum age***: 165 Ma.
***Age justification***: For the minimum age justification of Lebanese amber, see node 16. The maximum age

constraint is based on the Daohugou biota, which lacks representatives of the clade. See node 7 for justification.

**Node 41:** Crown Zopheridae: **98.17 Ma – 165 Ma.**
***Fossil taxon and specimen***: *Paleoendeitoma antennata* Deng, Ślipiński, Ren and Pang, 2017 [holotype, CNU-COL-MA-0312; paratype, CNU-COL-MA-0313 (amber inclusion): Key Lab of Insect Evolution and Environmental Changes, Capital Normal University, Beijing, China] lowermost Cenomanian, from an amber mine located near Noije Bum Village, Tanaing Town, Hukawng Valley, northern Myanmar (Deng et al. 2017b).
***Phylogenetic justification***: *Paleoendeitoma* can be placed in Zopheridae, subfamily Colydiinae, based on the antennal insertions concealed by frontal ridges, antennae distinctly clubbed with antennomere 3 as broad as the pedicel, procoxal cavities lacking lateral extensions, procoxae rounded with concealed trochantins, mesocoxae rounded with their cavities externally closed by sterna and the mesotrochantins concealed, abdomen with five ventrites, legs with visible trochanters and 4-segmented tarsi (Deng et al. 2017b).
***Minimum age***: 98.17 Ma.
***Soft maximum age***: 165 Ma.
***Age justification***: For the minimum age justification of Burmese amber from northern Myanmar, see node 12. The maximum age constraint is based on the Daohugou biota, which lacks representatives of the clade. See node 7 for justification.

**Node 42:** Crown Tenebrionidae: **122.2 Ma – 165 Ma.**
***Fossil taxon and specimen***: *Alphitopsis initialis* Kirejtshuk, Nabozhenko and Nel, 2011 [holotype, "A 31966" (exoskeleton with legs and fragments of antennae): Henan Geological Museum, Zhengzhou, China] from the Huanbanjigou of the Yixian Formation, Shangyuan, Beipiao city, Liaoning Province, China (Kirejtshuk et al. 2011).
***Phylogenetic justification***: The species has been assigned to the subfamily Alleculinae and tribe Gonoderini based on its five visible abdominal ventrites, simple tarsomeres without membranous ventral lobes, serrate antennae, and shape of pronotum and the body (Kirejtshuk et al. 2011). It is thus suitable for calibrating the node representing Crown Tenebrionidae.
***Minimum age***: 122.2 Ma.
***Soft maximum age***: 165 Ma.
***Age justification***: For the minimum age justification of the Yixian Formation, see node 14. The maximum age constraint is based on the Daohugou biota, which lacks representatives of the clade. See node 7 for justification.

**Node 43:** Alexiidae – Latridiidae: **125 Ma – 165 Ma.**
***Fossil taxon and specimen***: *Tetrameropsis mesozoica* Kirejtshuk and Azar, 2008 [holotype, "474A" (amber inclusion, sex indeterminate) Lebanese University, Faculty of Sciences II, Fanar, Lebanon] from the Early Cretaceous Lebanese amber originating from the outcrop Hammana – Mdeirij, Caza Baabda, Mouhafazet, Jabal Loubnan, central Lebanon (Kirejtshuk and Azar 2008).
***Phylogenetic justification***: The habitus of the species corresponds to corticariine Latridiidae (Shockley and Alekseev 2014). It was initially placed into its own subfamily, which was later raised to family level by Shockley and Alekseev (2014) where it has been hypothesised to be related to Latridiidae or Latridiidae + Akalyptoischiidae. In either case, the fossil is suitable for calibrating the split between Alexiidae and Latridiidae.
***Minimum age***: 125 Ma.
***Soft maximum age***: 165 Ma.
***Age justification***: For the minimum age justification of Lebanese amber, see node 16. The maximum age constraint is based on the Daohugou biota, which lacks representatives of the clade. See node 7 for justification.

**Node 44:** Crown Endomychidae: **98.17 Ma – 165 Ma.**

***Fossil taxon and specimen***: *Palaeomycetes foveolatus* Tomaszewska, Ślipiński and Ren, 2018 [holotype, CNU008072 (amber inclusion): Key Lab of Insect Evolution and Environmental Changes, Capital Normal University, Beijing, China] lowermost Cenomanian, from an amber mine located near Noije Bum Village, Tanaing Town, Hukawng Valley, northern Myanmar (Tomaszewska et al. 2018).
***Phylogenetic justification***: *Palaeomycetes* can be placed into crown Endomychidae based on having a frontoclypeal suture, lacking antennal grooves, antenna 11-segmented with a club, pronotum with transverse basal and longitudinal paired lateral sulci, visible portion of procoxae globular, procoxae separated by prosternal process, mesocoxal cavities externally open, metaventral paired postcoxal openings usually present, elytral epipleura well-developed, tarsi 4-segmented, abdominal ventrite 1 at least as long as ventrites 2 and 3 combined, and abdominal postcoxal lines absent (Tomaszewska 2000; Robertson et al. 2015)
***Minimum age***: 98.17 Ma.
***Soft maximum age***: 165 Ma.
***Age justification***: For the minimum age justification of Burmese amber from northern Myanmar, see node 12. The maximum age constraint is based on the Daohugou biota, which lacks representatives of the clade. See node 7 for justification.

**Node 45:** Crown Coccinellidae: **53 Ma – 125 Ma.**
***Fossil taxon and specimen***: *Rhyzobius antiquus* Kirejtshuk and Nel, 2012 [holotype, MNHN PA 11862 (amber inclusion): Muséum National d'Histoire Naturelle, Paris, France] from Eocene French amber obtained from the Farm Le Quesnoy locality, Chevrière, region of Creil, Oise department, northern France (Kirejtshuk and Nel 2012).
***Phylogenetic justification***: Affinity to Coccinellidae is indicated a broadly rounded body with a convex dorsum and flattened ventral side, clubbed antennae, the presence of a postcoxal line on the first abdominal ventrite, and pseudotrimerous tarsi (Kirejtshuk and Nel 2012).
***Minimum age***: 53 Ma.
***Soft maximum age***: 98.17 Ma.
***Age justification***: For the minimum age justification of Eocene Oise amber, see node 38. The maximum age constraint is based on the Daohugou biota, which lacks representatives of the clade. See node 7 for justification.

**Node 46:** Boganiidae – Erotylidae: **165 Ma – 235 Ma.**
***Fossil taxon and specimen***: *Palaeoboganium jurassicum* Liu, Ślipiński, Lawrence, Ren and Pang, 2018 [holotype, CNU-COL-NN201601PC (exoskeleton with appendages): Key Lab of Insect Evolution and Environmental Changes, Capital Normal University, Beijing, China] from Daohugou Village, Middle Jurassic Jiulongshan Formation, Ningcheng County, Inner Mongolia, China (Liu et al. 2018).
***Phylogenetic justification***: *Palaeoboganium jurassicum* is the earliest fossil representative of the family Boganiidae, as indicated by the head with an arcuate frontoclypeal suture, labrum not visible externally, mandible with a dorsal tubercle fitting into a depression on lateral edge of clypeus, subantennal grooves absent, antennae short and weakly capitate, pro- and mesocoxae with exposed trochantins, metacoxae narrowly separated and, not extending laterally to meet elytra, and abdomen with five freely articulated ventrites. A phylogenetic analysis of adult external characters recovered *Palaeoboganium* as the sister group of *Paracucujus* and *Metacucujus* (Liu et al. 2018), making it suitable for calibrating the split between Boganiidae and Erotylidae.
***Minimum age***: 165 Ma.
***Soft maximum age***: 235 Ma.
***Age justification***: See node 7 for the minimum age justification of the Daohugou locality. The soft maximum is from the Madygen Formation at the Dzhailyau-Cho locality, Lyailyakskii District, Batken Region, southwest Kyrgyzstan, Central Asia, as justified in node 7.

**Node 47:** Helotidae – (Protocucujidae + Sphindidae): **125 Ma – 235 Ma.**

***Fossil taxon and specimen***: *Libanopsis poinari* Kirejtshuk, 2015 [holotype, "841 BC" (amber inclusion): Lebanese University, Faculty of Sciences II, Department of Natural Sciences, Beirut, Lebanon] from the Early Cretaceous Lebanese amber originating from the outcrop Hammana – Mdeirij, Caza Baabda, Mouhafazet, Jabal Loubnan, central Lebanon (Kirejtshuk et al. 2015a).
***Phylogenetic justification***: *Libanopsis poinari* can be placed within Sphindidae based on the sparse sculpture of dorsal integument, large coarsely faceted eyes, antennal morphology, undulate lateral pronotal carinas, and small serration in pretarsal claws (Kirejtshuk et al. 2015a).
***Minimum age***: 125 Ma.
***Soft maximum age***: 235 Ma.
***Age justification***: For the minimum age justification of Lebanese amber, see node 16. The maximum age constraint is based on the Daohugou biota, which lacks representatives of the clade. See node 7 for justification.

**Node 48:** Monotomidae – (Kateretidae + Nitidulidae): **165 Ma – 235 Ma.**
***Fossil taxon and specimen***: *Jurorhizophagus alienus* Cai, Ślipiński and Huang, 2015 [holotype, NIGP 161706 (exoskeleton with antennae and partly preserved legs): Nanjing Institute of Geology and Palaeontology, Chinese Academy of Sciences, Nanjing, China] from the Haifanggou Formation at Daohugou, Ningcheng County, Inner Mongolia, northeastern China (Cai et al. 2015c).
***Phylogenetic justification***: *Jurorhizophagus alienus* can be placed among monotomids based on its narrowly elongate body, prognathous head with lateral finely faceted eyes, procoxal cavities closed externally, elytra truncate apically and exposing two tergites, and 5-segmented abdomen with relatively long ventrites 1 and 5 (Sen Gupta 1988; Cai et al. 2015c).
***Minimum age***: 165 Ma.
***Soft maximum age***: 235 Ma.
***Age justification***: For the minimum age justification of the Daohugou biota, see node 7. The soft maximum age (*ca*. 235 Ma) is derived from the Madygen Formation at Dzhailyau-Cho, Kyrgyzstan. See node 8 for justification.

**Node 49:** Crown Nitidulidae: **119 Ma – 165 Ma.**
***Fossil taxon and specimen***: *Crepuraea archaica* Kirejtshuk, 1990 [holotype, PIN 3064/7189 (exoskeleton): Borissiak Paleontological Institute, Russian Academy of Sciences, Moscow, Russia] Baissa locality situated on the left bank of the Vitim River, 8 km below the mouth of the Baissa River, Buryat Republic, west Transbaikalia, Russia.
***Phylogenetic justification***: As discussed in Kirejtshuk and Ponomarenko (1990) the genus *Crepuraea* has a close affinity to the extant *Epuraea*, which is placed in the subfamily Epuraeinae and tribe Epuraeini (Kirejtshuk 2008). Based on molecular phylogenies of Nitidulidae (Cline et al. 2014), this would place *Crepuraea* well within crown Nitidulidae, making it a suitable candidate for calibrating this node.
***Minimum age***: 119 Ma.
***Soft maximum age***: 165 Ma.
***Age justification***: The exact age of the lacustrine sediments of the Zaza Formation at Baissa is disputable. While early estimates suggested an Early Cretaceous (Neocomian–Aptian) age, most palaeoentomologists date them as Valanginian–Hauterivian (Zherikhin et al. 1999). The co-occurrence of *Coptoclava longipoda, Ephemeropsis trisetalis* and *Lycoptera* in the Zaza Formation as well as the Jiufotang Formation in western Liaoning, China suggests that the two may be contemporaneous. Given that the latter has been dated to 120 ± 1.0 Ma (unpubl. data; Kirejtshuk and Ponomarenko 1990), we use 119 Ma as the minimum constraint on the clade representing crown Nitidulidae. The maximum age constraint is based on the Daohugou biota, which lacks representatives of the clade. See node 7 for justification.

**Node 50:** Crown Silvanidae: **98.17 Ma – 165 Ma.**
***Fossil taxon and specimen***: *Protoliota antennatus* Liu, Ślipiński, Wang and Pang, 2019 [holotype, No. CNU008072 (amber inclusion): Key Lab of Insect Evolution and Environmental Changes, Capital Normal

University, Beijing, China] lowermost Cenomanian, from an amber mine located near Noije Bum Village, Tanaing Town, Hukawng Valley, northern Myanmar (Liu et al. 2019).
***Phylogenetic justification*:** The placement of *Protoliota* within Silvanidae is supported by its overall body form, long filiform antennae with long scape and without club, 5-5-5 tarsi, and narrowly open procoxal cavities (Liu et al. 2019). The distinctly lobed tarsomeres and prominent sexual dimorphism manifesting in the presence of male mandibular horns and antennal length (Thomas 2004; Cai and Huang 2019) suggest a placement in Brontinae.
***Minimum age*:** 98.17 Ma.
***Soft maximum age*:** 165 Ma.
***Age justification*:** For the minimum age justification of Burmese amber from northern Myanmar, see node 12. The maximum age constraint is based on the Daohugou biota, which lacks representatives of the clade. See node 7 for justification.

**Node 51:** Nemonychidae – Anthribidae: **155 Ma – 235 Ma.**
***Fossil taxon and specimen*:** *Protoscelis longicornis* (Medvedev, 1968) [holotype, PIN, no. 2066/3241 (exoskeleton): Borissiak Paleontological Institute, Russian Academy of Sciences, Moscow, Russia] from the Karabastau Formation of the Karatau Range near the village of Mikhailovka, Chimkent Region, Chayan District, southern Kazakhstan (Medvedev 1968; Legalov 2013).
***Phylogenetic justification*:** The presence of a reduced rostrum, a three-segmented and rather compact antennal club, developed subbasal line on pronotum, and sexually dimorphic antennae place the fossil into Anthribidae (Legalov 2013).
***Minimum age*:** 155 Ma.
***Soft maximum age*:** 235 Ma.
***Age justification*:** For the minimum age justification of the Karatau Range, see node 5. The soft maximum age (*ca*. 235 Ma) is derived from the Madygen Formation at Dzhailyau-Cho, Kyrgyzstan. See node 8 for justification.

**Node 52:** Belidae – (Attelabidae, (Brentidae, Curculionidae)): **165 Ma – 235 Ma.**
***Fossil taxon and specimen*:** *Sinoeuglypheus daohugouensis* Yu, Davis and Shih, 2019 [holotype, CNU-COL-NN2011122 (exoskeleton with appendages preserved in lateral aspect, sex unknown): Key Lab of Insect Evolution and Environmental Changes, Capital Normal University, Beijing, China] from the Haifanggou Formation at Daohugou, Ningcheng County, Inner Mongolia, northeastern China (Yu et al. 2019).
***Phylogenetic justification*:** The species can be assigned to Belidae on the basis of the irregularly arranged punctures on the elytra, elongated first flagellomere of the antennae, outer longitudinal carina and an apical setal brush on the fore tibiae, wide prothoracic cavity for reception of the procoxa, form of the meso- and metathoracic episterna and epimera, small flange along the elytral basal margin, and lateral carinae on the abdominal ventrites (Yu et al. 2019).
***Minimum age*:** 165 Ma.
***Soft maximum age*:** 235 Ma.
***Age justification*:** For the minimum age justification of the Daohugou biota, see node 7. The soft maximum age (*ca*. 235 Ma) is derived from the Madygen Formation at Dzhailyau-Cho, Kyrgyzstan. See node 8 for justification.

**Node 53:** Brentidae **–** Curculionidae: **112.6 Ma – 165 Ma.**
***Fossil taxon and specimen*:** *Axelrodiellus ruptus* Zherikhin and Gratshev, 2004 [holotype, AMNH 43312 (exoskeleton lacking legs, with fragments of antennae): American Museum of Natural History, New York City, USA] from the Lower Cretaceous (Aptian) Crato Formation, northeastern Brazil (Zherikhin and Gratshev 2004).
***Phylogenetic justification:*** The species is placed in Brentidae on the basis of having an unpaired gular suture, narrow and loose antennal club, enlarged trochanters, and fused and strongly enlarged first two abdominal sternites. Assignment to Eurhynchinae is supported by the sloping trochanters and the bases of the femora

reaching the coxae, complete and deep suture between the first two ventrites, gular suture running from the base of the rostrum to the eyes, and striated gular region (Zherikhin and Gratshev 2004).
***Minimum age***: 112.6 Ma.
***Soft maximum age***: 165 Ma.
***Age justification***: For the minimum age justification of the Crato Formation, see node 11. The maximum age constraint is based on the Daohugou biota, which lacks representatives of the clade. See node 7 for justification.

#### **Node 54:** Crown Chrysomelidae: **122.2 Ma – 165 Ma.**
***Fossil taxon and specimen***: *Mesolpinus antennatus* Kirejtshuk, Moseyko and Ren, 2015 [holotype, CNU-COL-LB-2009840 (exoskeleton with fragments of appendages preserved in ventral aspect): Key Lab of Insect Evolution and Environmental Changes, Capital Normal University, Beijing, China] from the Huanbanjigou of the Yixian Formation, Shangyuan, Beipiao city, Liaoning Province, China (Kirejtshuk et al. 2015b).
***Phylogenetic justification***: The genus can be referred to the subfamily Chrysomelinae based on the hypognathous head not narrowed at the base, antennal cavities located in front of eyes and widely separated, frontoclypeus subquadrangular, behind the mandibles and separated from the posterior part of the frons by a wide V-shaped suture, the pronotum laterally margined, all coxae moderately separated, the prosternal process apparently lacking lateral structures for the reception of the procoxae, procoxae distinctly transverse, mesoventrite lacking projections for reception of the mesocoxae, metepisterna simple and lacking projections, the abdomen with ventrite 1 longest and other ventrites subequal in length, not fused, and tarsomere 3 bilobed (Kirejtshuk et al. 2015b).
***Minimum age***: 122.2 Ma.
***Soft maximum age***: 165 Ma.
***Age justification***: For the minimum age justification of the Yixian Formation, see node 14. The maximum age constraint is based on the Daohugou biota, which lacks representatives of the clade. See node 7 for justification.

#### **Node 55:** Crown Cerambycidae: **122.2 Ma – 165 Ma.**
***Fossil taxon and specimen***: *Cretoprionus liutiaogouensis* Wang, Ma, McKenna, Yan, Zhang and Jarzembowski, 2014 [holotype, NIGP154953a, b (exoskeleton with appendages): Nanjing Institute of Geology and Palaeontology, Chinese Academy of Sciences, Nanjing, China] from the Huanbanjigou of the Yixian Formation, Shangyuan, Beipiao city, Liaoning Province, China (Wang et al. 2014).
***Phylogenetic justification***: *Cretoprionus liutiaogouensis* can be assigned to Cerambycidae based on its large and robust body, simple mandibular apex, pseudotetramerous tarsi, raised antennal insertions, and long antennae. It can furthermore be placed in Prioninae due to the very large and robust body, absence of a stridulatory plate on the mesonotum, pseudotetramerous tarsi, mouthparts projecting anteriorly, and lateral carinae on the prothorax (Turnbow and Thomas 2002; Wang et al. 2014; Vitali, 2019).
***Minimum age***: 122.2 Ma.
***Soft maximum age***: 165 Ma.
***Age justification***: For the minimum age justification of the Yixian Formation, see node 14. The maximum age constraint is based on the Daohugou biota, which lacks representatives of the clade. See node 7 for justification.

#### **Node 56:** Sagrinae – Bruchinae: **78 Ma – 155 Ma.**
***Fossil taxon and specimen***: *Mesopachymerus antiqua* Poinar, 2005 [holotype, TMP 91.148.771 (amber inclusion): Royal Tyrrell Museum in Drumheller, Albert Canada] from Cretaceous Canadian amber, obtained from the Grassy Lake amber deposit, Foremost Formation, Alberta, Canada (Poinar 2005).
***Phylogenetic justification***: The widened protarsus 1 reliably places *Mesopachymerus antiqua* among bruchines, making it a suitable candidate for calibrating the split between Sagrinae and Bruchinae (Poinar 2005).

***Minimum age***: 78 Ma.
***Soft maximum age***: 155 Ma.
***Age justification***: The age of Canadian amber has been estimated as approximately 78 – 79 Ma based on biostratigraphic correlation (McKellar et al. 2013). We conservatively use 78 Ma as the minimum age constraint. The soft maximum age is derived from the Karabastau Formation of the Karatau Range, see node 5 for justification.

**Node 57:** Galerucinae – Chrysomelinae: **83.4 Ma – 165 Ma.**
***Fossil taxon and specimen***: *Taimyraltica calcarata* Nadein, 2018 [holotype, PIN 3311/1203 (amber inclusion): Borissiak Paleontological Institute, Russian Academy of Sciences, Moscow, Russia] from Cretaceous Taimyr amber collected from the right bank of the Maimecha River, Taimyr Peninsula, Kheta Formation, Krasnoyarskiy Krai, Russia (Nadein and Perkovsky 2018).
***Phylogenetic justification***: *Taimyraltica* can be attributed to Galerucinae based on its elongate body, hypognathous head without a constriction behind the eyes, antennal insertions situated between eyes at a distance from anterior margin of frons, eyes without emarginations, pronotum with a lateral margin and not constricted, pronotal base with a transverse furrow, procoxae transverse, elytra regularly punctured, and pygidium not vertical (Nadein and Beždek 2014; Nadein and Perkovsky 2018).
***Minimum age***: 83.4 Ma.
***Soft maximum age***: 165 Ma.
***Age justification***: *Taimyraltica calcarata* occurs in the Late Cretaceous Taimyr amber from the Kheta Formation at the Yantardakh locality, Taimyr Peninsula, Russia. The age of the Kheta Formation is Coniacian-Santonian according to Saks et al. (1959). All amber from the Yantardakh locality has been found in the upper horizons of the formation, and is therefore thought to be Santonian (Zherikhin, 1978) or even upper Santonian based on the presence of marine bivalves of the genus *Inoceramus* in the overlying Mutino Formation (Perkovsky and Makarkin, 2015). As such, the age of Taimyr amber from Yantardakh is late Santonian, the upper boundary of which is dated to 83.6 ± 0.2 Ma (International Chronostratigraphic Chart, v. 2018/08). The minimum age constraint for this node is 83.4 Ma. The maximum age constraint is based on the Daohugou biota, which lacks representatives of the clade. See node 7 for justification.

### Beetle phylogeny and evolutionary timescale

**Figure S1.** Test of model-depenmdent phylogenetic signal on a reduced 138 taxon dataset analysed with the LG model in IQ-TREE.

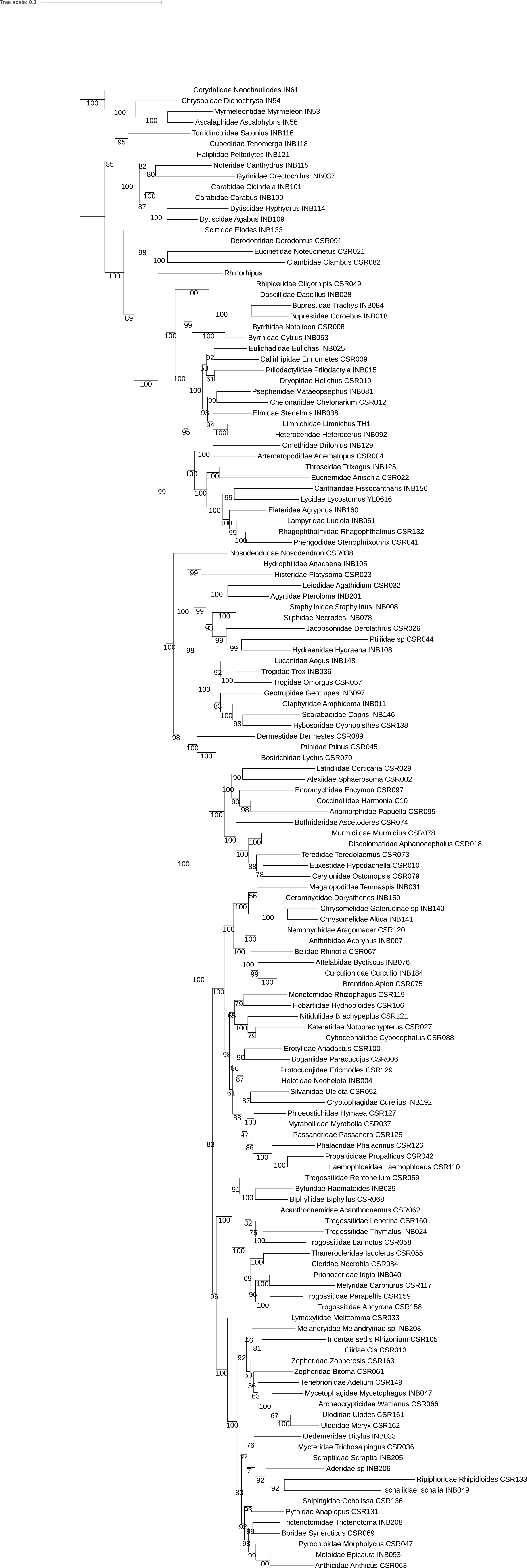

Tree scale: 0.1
Corydalidae Neochauliodes IN61
Chrysopidae Dichochrysa IN54
Myrmeleontidae Myrmeleon IN53
Ascalaphidae Ascalohybris IN56
Torridincolidae Satonius INB116
Cupedidae Tenomerga INB118
Haliplidae Peltodytes INB121
Noteridae Canthydrus INB115
Gyrinidae Orectochilus INB037
Carabidae Cicindela INB101
Carabidae Carabus INB100
Dytiscidae Hyphydrus INB114
Dytiscidae Agabus INB109
Scirtidae Elodes INB133
Derodontidae Derodontus CSR091
Eucinetidae Noteucinetus CSR021
Clambidae Clambus CSR082
Rhinorhipus
Rhipiceridae Oligorhipis CSR049
Dascillidae Dascillus INB028
Buprestidae Trachys INB084
Buprestidae Coroebus INB018
Byrrhidae Notolioon CSR008
Byrrhidae Cytilus INB053
Eulichadidae Eulichas INB025
Callirhipidae Ennometes CSR009
Ptilodactylidae Ptilodactyla INB015
Dryopidae Helichus CSR019
Psephenidae Mataeopsephus INB081
Chelonariidae Chelonarium CSR012
Elmidae Stenelmis INB038
Limnichidae Limnichus TH1
Heteroceridae Heterocerus INB092
Omethidae Drilonius INB129
Artematopodidae Artematopus CSR004
Throscidae Trixagus INB125
Eucnemidae Anischia CSR022
Cantharidae Fissocantharis INB156
Lycidae Lycostomus YL0616
Elateridae Agrypnus INB160
Lampyridae Luciola INB061
Rhagophthalmidae Rhagophthalmus CSR132
Phengodidae Stenophrixothrix CSR041
Nosodendridae Nosodendron CSR038
Hydrophilidae Anacaena INB105
Histeridae Platysoma CSR023
Leiodidae Agathidium CSR032
Agyrtidae Pteroloma INB201
Staphylinidae Staphylinus INB008
Silphidae Necrodes INB078
Jacobsoniidae Derolathrus CSR026
Ptiliidae sp CSR044
Hydraenidae Hydraena INB108
Lucanidae Aegus INB148
Trogidae Trox INB036
Trogidae Omorgus CSR057
Geotrupidae Geotrupes INB097
Glaphyridae Amphicoma INB011
Scarabaeidae Copris INB146
Hybosoridae Cyphopisthes CSR138
Dermestidae Dermestes CSR089
Ptinidae Ptinus CSR045
Bostrichidae Lyctus CSR070
Latridiidae Corticaria CSR029
Alexiidae Sphaerosoma CSR002
Endomychidae Encymon CSR097
Coccinellidae Harmonia C10
Anamorphidae Papuella CSR095
Bothrideridae Ascetoderes CSR074
Murmidiidae Murmidius CSR078
Discolomatidae Aphanocephalus CSR018
Teredidae Teredolaemus CSR073
Euxestidae Hypodacnella CSR010
Cerylonidae Ostomopsis CSR079
Megalopodidae Temnaspis INB031
Cerambycidae Dorysthenes INB150
Chrysomelidae Galerucinae sp INB140
Chrysomelidae Altica INB141
Nemonychidae Aragomacer CSR120
Anthribidae Acorynus INB007
Belidae Rhinotia CSR067
Attelabidae Byctiscus INB076
Curculionidae Curculio INB184
Brentidae Apion CSR075
Monotomidae Rhizophagus CSR119
Hobartiidae Hydnobioides CSR106
Nitidulidae Brachypeplus CSR121
Kateretidae Notobrachypterus CSR027
Cybocephalidae Cybocephalus CSR088
Erotylidae Anadastus CSR100
Boganiidae Paracucujus CSR006
Protocucujidae Ericmodes CSR129
Helotidae Neohelota INB004
Silvanidae Uleiota CSR052
Cryptophagidae Curelius INB192
Phloeostichidae Hymaea CSR127
Myraboliidae Myrabolia CSR037
Passandridae Passandra CSR125
Phalacridae Phalacrinus CSR126
Propalticidae Propalticus CSR042
Laemophloeidae Laemophloeus CSR110
Trogossitidae Rentonellum CSR059
Byturidae Haematoides INB039
Biphyllidae Biphyllus CSR068
Acanthocnemidae Acanthocnemus CSR062
Trogossitidae Leperina CSR160
Trogossitidae Thymalus INB024
Trogossitidae Larinotus CSR058
Thanerocleridae Isoclerus CSR055
Cleridae Necrobia CSR084
Prionoceridae Idgia INB040
Melyridae Carphurus CSR117
Trogossitidae Parapeltis CSR159
Trogossitidae Ancyrona CSR158
Lymexylidae Melittomma CSR033
Melandryidae Melandryinae sp INB203
Incertae sedis Rhizonium CSR105
Ciidae Cis CSR013
Zopheridae Zopherosis CSR163
Zopheridae Bitoma CSR061
Tenebrionidae Adelium CSR149
Mycetophagidae Mycetophagus INB047
Archeocrypticidae Wattianus CSR066
Ulodidae Ulodes CSR161
Ulodidae Meryx CSR162
Oedemeridae Ditylus INB033
Mycteridae Trichosalpingus CSR036
Scraptiidae Scraptia INB205
Aderidae sp INB206
Ripiphoridae Rhipidioides CSR133
Ischaliidae Ischalia INB049
Salpingidae Ocholissa CSR136
Pythidae Anaplopus CSR131
Trictenotomidae Trictenotoma INB208
Boridae Synercticus CSR069
Pyrochroidae Morpholycus CSR047
Meloidae Epicauta INB093
Anthicidae Anthicus CSR063

**Figure S2.** Test of model-depenmdent phylogenetic signal on a reduced 138 taxon dataset analysed with the WAG model in IQ-TREE.

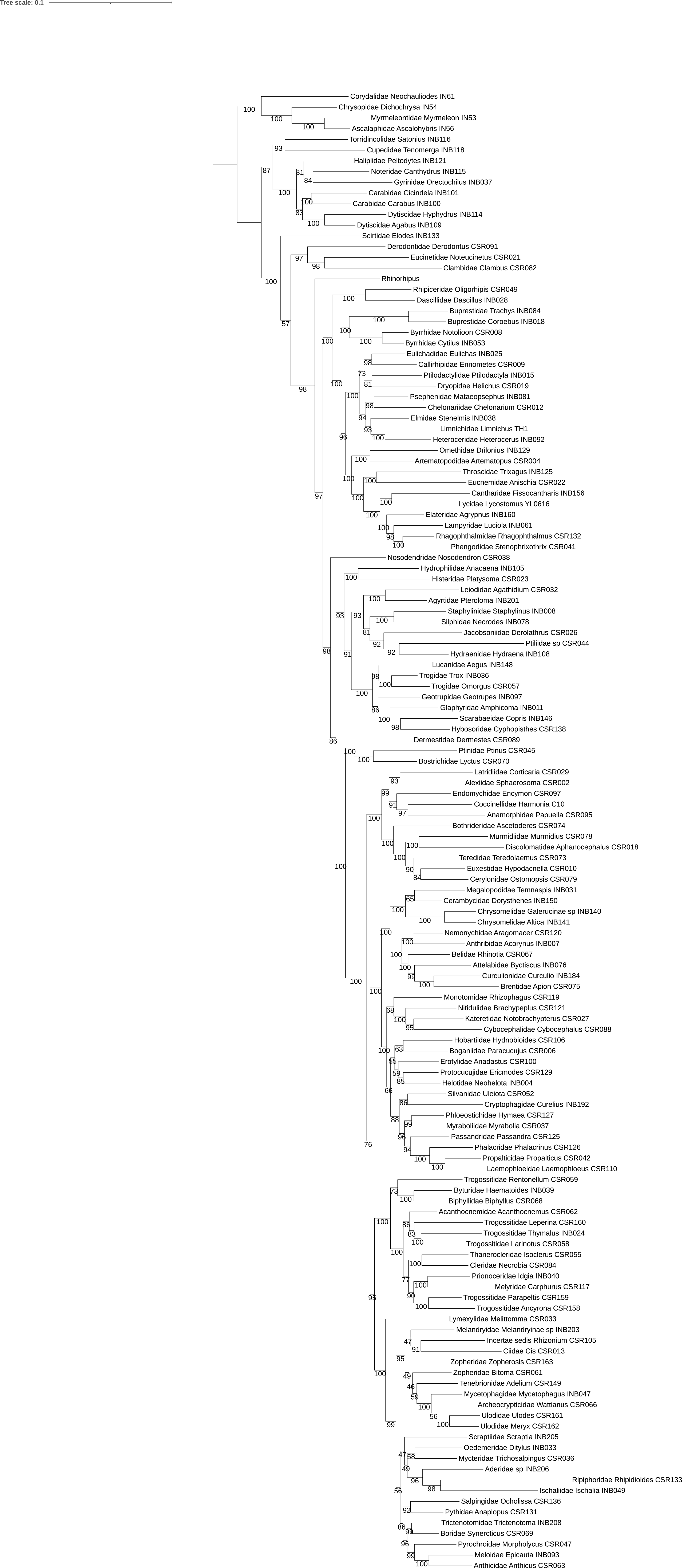
Tree scale: 0.1
Corydalidae Neochauliodes IN61
Chrysopidae Dichochrysa IN54
Myrmeleontidae Myrmeleon IN53
Ascalaphidae Ascalohybris IN56
Torridincolidae Satonius INB116
Cupedidae Tenomerga INB118
Haliplidae Peltodytes INB121
Noteridae Canthydrus INB115
Gyrinidae Orectochilus INB037
Carabidae Cicindela INB101
Carabidae Carabus INB100
Dytiscidae Hyphydrus INB114
Dytiscidae Agabus INB109
Scirtidae Elodes INB133
Derodontidae Derodontus CSR091
Eucinetidae Noteucinetus CSR021
Clambidae Clambus CSR082
Rhinorhipus
Rhipiceridae Oligorhipis CSR049
Dascillidae Dascillus INB028
Buprestidae Trachys INB084
Buprestidae Coroebus INB018
Byrrhidae Notolioon CSR008
Byrrhidae Cytilus INB053
Eulichadidae Eulichas INB025
Callirhipidae Ennometes CSR009
Ptilodactylidae Ptilodactyla INB015
Dryopidae Helichus CSR019
Psephenidae Mataeopsephus INB081
Chelonariidae Chelonarium CSR012
Elmidae Stenelmis INB038
Limnichidae Limnichus TH1
Heteroceridae Heterocerus INB092
Omethidae Drilonius INB129
Artematopodidae Artematopus CSR004
Throscidae Trixagus INB125
Eucnemidae Anischia CSR022
Cantharidae Fissocantharis INB156
Lycidae Lycostomus YL0616
Elateridae Agrypnus INB160
Lampyridae Luciola INB061
Rhagophthalmidae Rhagophthalmus CSR132
Phengodidae Stenophrixothrix CSR041
Nosodendridae Nosodendron CSR038
Hydrophilidae Anacaena INB105
Histeridae Platysoma CSR023
Leiodidae Agathidium CSR032
Agyrtidae Pteroloma INB201
Staphylinidae Staphylinus INB008
Silphidae Necrodes INB078
Jacobsoniidae Derolathrus CSR026
Ptiliidae sp CSR044
Hydraenidae Hydraena INB108
Lucanidae Aegus INB148
Trogidae Trox INB036
Trogidae Omorgus CSR057
Geotrupidae Geotrupes INB097
Glaphyridae Amphicoma INB011
Scarabaeidae Copris INB146
Hybosoridae Cyphopisthes CSR138
Dermestidae Dermestes CSR089
Ptinidae Ptinus CSR045
Bostrichidae Lyctus CSR070
Latridiidae Corticaria CSR029
Alexiidae Sphaerosoma CSR002
Endomychidae Encymon CSR097
Coccinellidae Harmonia C10
Anamorphidae Papuella CSR095
Bothrideridae Ascetoderes CSR074
Murmidiidae Murmidius CSR078
Discolomatidae Aphanocephalus CSR018
Teredidae Teredolaemus CSR073
Euxestidae Hypodacnella CSR010
Cerylonidae Ostomopsis CSR079
Megalopodidae Temnaspis INB031
Cerambycidae Dorysthenes INB150
Chrysomelidae Galerucinae sp INB140
Chrysomelidae Altica INB141
Nemonychidae Aragomacer CSR120
Anthribidae Acorynus INB007
Belidae Rhinotia CSR067
Attelabidae Byctiscus INB076
Curculionidae Curculio INB184
Brentidae Apion CSR075
Monotomidae Rhizophagus CSR119
Nitidulidae Brachypeplus CSR121
Kateretidae Notobrachypterus CSR027
Cybocephalidae Cybocephalus CSR088
Hobartiidae Hydnobioides CSR106
Boganiidae Paracucujus CSR006
Erotylidae Anadastus CSR100
Protocucujidae Ericmodes CSR129
Helotidae Neohelota INB004
Silvanidae Uleiota CSR052
Cryptophagidae Curelius INB192
Phloeostichidae Hymaea CSR127
Myraboliidae Myrabolia CSR037
Passandridae Passandra CSR125
Phalacridae Phalacrinus CSR126
Propalticidae Propalticus CSR042
Laemophloeidae Laemophloeus CSR110
Trogossitidae Rentonellum CSR059
Byturidae Haematoides INB039
Biphyllidae Biphyllus CSR068
Acanthocnemidae Acanthocnemus CSR062
Trogossitidae Leperina CSR160
Trogossitidae Thymalus INB024
Trogossitidae Larinotus CSR058
Thanerocleridae Isoclerus CSR055
Cleridae Necrobia CSR084
Prionoceridae Idgia INB040
Melyridae Carphurus CSR117
Trogossitidae Parapeltis CSR159
Trogossitidae Ancyrona CSR158
Lymexylidae Melittomma CSR033
Melandryidae Melandryinae sp INB203
Incertae sedis Rhizonium CSR105
Ciidae Cis CSR013
Zopheridae Zopherosis CSR163
Zopheridae Bitoma CSR061
Tenebrionidae Adelium CSR149
Mycetophagidae Mycetophagus INB047
Archeocrypticidae Wattianus CSR066
Ulodidae Ulodes CSR161
Ulodidae Meryx CSR162
Scraptiidae Scraptia INB205
Oedemeridae Ditylus INB033
Mycteridae Trichosalpingus CSR036
Aderidae sp INB206
Ripiphoridae Rhipidioides CSR133
Ischaliidae Ischalia INB049
Salpingidae Ocholissa CSR136
Pythidae Anaplopus CSR131
Trictenotomidae Trictenotoma INB208
Boridae Synercticus CSR069
Pyrochroidae Morpholycus CSR047
Meloidae Epicauta INB093
Anthicidae Anthicus CSR063

**Figure S3.** Test of model-depenmdent phylogenetic signal on a reduced 138 taxon dataset analysed with the JTT model in IQ-TREE.

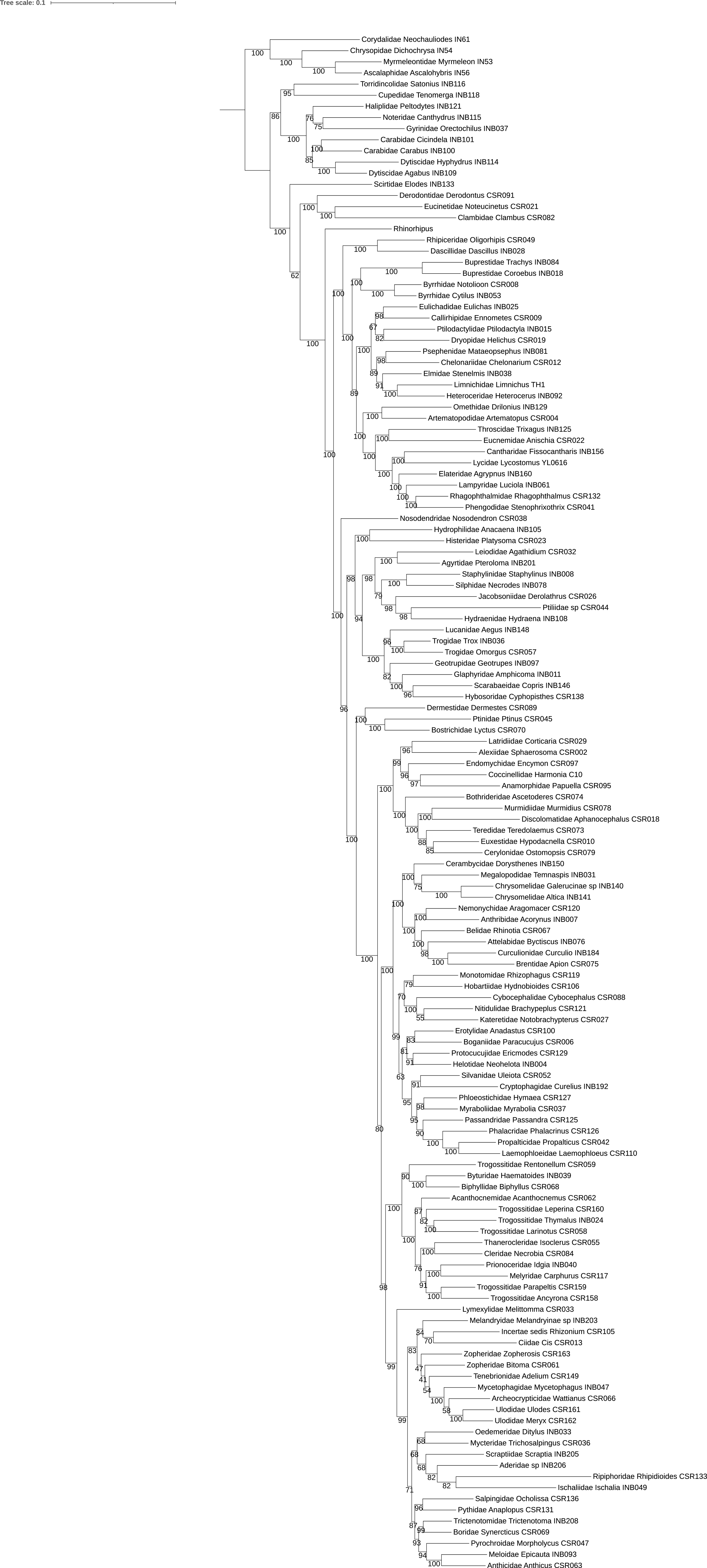
Tree scale: 0.1
Corydalidae Neochauliodes IN61
Chrysopidae Dichochrysa IN54
Myrmeleontidae Myrmeleon IN53
Ascalaphidae Ascalohybris IN56
Torridincolidae Satonius INB116
Cupedidae Tenomerga INB118
Haliplidae Peltodytes INB121
Noteridae Canthydrus INB115
Gyrinidae Orectochilus INB037
Carabidae Cicindela INB101
Carabidae Carabus INB100
Dytiscidae Hyphydrus INB114
Dytiscidae Agabus INB109
Scirtidae Elodes INB133
Derodontidae Derodontus CSR091
Eucinetidae Noteucinetus CSR021
Clambidae Clambus CSR082
Rhinorhipus
Rhipiceridae Oligorhipis CSR049
Dascillidae Dascillus INB028
Buprestidae Trachys INB084
Buprestidae Coroebus INB018
Byrrhidae Notolioon CSR008
Byrrhidae Cytilus INB053
Eulichadidae Eulichas INB025
Callirhipidae Ennometes CSR009
Ptilodactylidae Ptilodactyla INB015
Dryopidae Helichus CSR019
Psephenidae Mataeopsephus INB081
Chelonariidae Chelonarium CSR012
Elmidae Stenelmis INB038
Limnichidae Limnichus TH1
Heteroceridae Heterocerus INB092
Omethidae Drilonius INB129
Artematopodidae Artematopus CSR004
Throscidae Trixagus INB125
Eucnemidae Anischia CSR022
Cantharidae Fissocantharis INB156
Lycidae Lycostomus YL0616
Elateridae Agrypnus INB160
Lampyridae Luciola INB061
Rhagophthalmidae Rhagophthalmus CSR132
Phengodidae Stenophrixothrix CSR041
Nosodendridae Nosodendron CSR038
Hydrophilidae Anacaena INB105
Histeridae Platysoma CSR023
Leiodidae Agathidium CSR032
Agyrtidae Pteroloma INB201
Staphylinidae Staphylinus INB008
Silphidae Necrodes INB078
Jacobsoniidae Derolathrus CSR026
Ptiliidae sp CSR044
Hydraenidae Hydraena INB108
Lucanidae Aegus INB148
Trogidae Trox INB036
Trogidae Omorgus CSR057
Geotrupidae Geotrupes INB097
Glaphyridae Amphicoma INB011
Scarabaeidae Copris INB146
Hybosoridae Cyphopisthes CSR138
Dermestidae Dermestes CSR089
Ptinidae Ptinus CSR045
Bostrichidae Lyctus CSR070
Latridiidae Corticaria CSR029
Alexiidae Sphaerosoma CSR002
Endomychidae Encymon CSR097
Coccinellidae Harmonia C10
Anamorphidae Papuella CSR095
Bothrideridae Ascetoderes CSR074
Murmidiidae Murmidius CSR078
Discolomatidae Aphanocephalus CSR018
Teredidae Teredolaemus CSR073
Euxestidae Hypodacnella CSR010
Cerylonidae Ostomopsis CSR079
Cerambycidae Dorysthenes INB150
Megalopodidae Temnaspis INB031
Chrysomelidae Galerucinae sp INB140
Chrysomelidae Altica INB141
Nemonychidae Aragomacer CSR120
Anthribidae Acorynus INB007
Belidae Rhinotia CSR067
Attelabidae Byctiscus INB076
Curculionidae Curculio INB184
Brentidae Apion CSR075
Monotomidae Rhizophagus CSR119
Hobartiidae Hydnobioides CSR106
Cybocephalidae Cybocephalus CSR088
Nitidulidae Brachypeplus CSR121
Kateretidae Notobrachypterus CSR027
Erotylidae Anadastus CSR100
Boganiidae Paracucujus CSR006
Protocucujidae Ericmodes CSR129
Helotidae Neohelota INB004
Silvanidae Uleiota CSR052
Cryptophagidae Curelius INB192
Phloeostichidae Hymaea CSR127
Myraboliidae Myrabolia CSR037
Passandridae Passandra CSR125
Phalacridae Phalacrinus CSR126
Propalticidae Propalticus CSR042
Laemophloeidae Laemophloeus CSR110
Trogossitidae Rentonellum CSR059
Byturidae Haematoides INB039
Biphyllidae Biphyllus CSR068
Acanthocnemidae Acanthocnemus CSR062
Trogossitidae Leperina CSR160
Trogossitidae Thymalus INB024
Trogossitidae Larinotus CSR058
Thanerocleridae Isoclerus CSR055
Cleridae Necrobia CSR084
Prionoceridae Idgia INB040
Melyridae Carphurus CSR117
Trogossitidae Parapeltis CSR159
Trogossitidae Ancyrona CSR158
Lymexylidae Melittomma CSR033
Melandryidae Melandryinae sp INB203
Incertae sedis Rhizonium CSR105
Ciidae Cis CSR013
Zopheridae Zopherosis CSR163
Zopheridae Bitoma CSR061
Tenebrionidae Adelium CSR149
Mycetophagidae Mycetophagus INB047
Archeocrypticidae Wattianus CSR066
Ulodidae Ulodes CSR161
Ulodidae Meryx CSR162
Oedemeridae Ditylus INB033
Mycteridae Trichosalpingus CSR036
Scraptiidae Scraptia INB205
Aderidae sp INB206
Ripiphoridae Rhipidioides CSR133
Ischaliidae Ischalia INB049
Salpingidae Ocholissa CSR136
Pythidae Anaplopus CSR131
Trictenotomidae Trictenotoma INB208
Boridae Synercticus CSR069
Pyrochroidae Morpholycus CSR047
Meloidae Epicauta INB093
Anthicidae Anthicus CSR063

**Figure S4.** Test of model-depenmdent phylogenetic signal on a reduced 138 taxon dataset analysed with the GTR20 model in IQ-TREE.

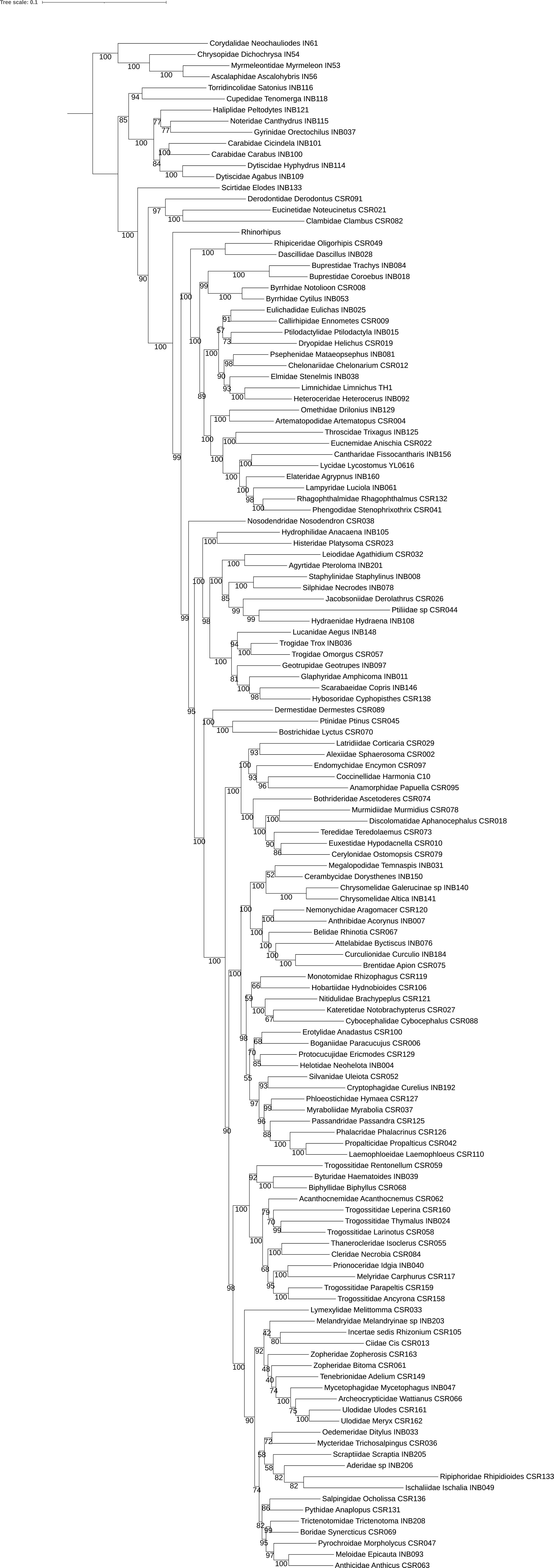

Tree scale: 0.1
Corydalidae Neochauliodes IN61
Chrysopidae Dichochrysa IN54
Myrmeleontidae Myrmeleon IN53
Ascalaphidae Ascalohybris IN56
Torridincolidae Satonius INB116
Cupedidae Tenomerga INB118
Haliplidae Peltodytes INB121
Noteridae Canthydrus INB115
Gyrinidae Orectochilus INB037
Carabidae Cicindela INB101
Carabidae Carabus INB100
Dytiscidae Hyphydrus INB114
Dytiscidae Agabus INB109
Scirtidae Elodes INB133
Derodontidae Derodontus CSR091
Eucinetidae Noteucinetus CSR021
Clambidae Clambus CSR082
Rhinorhipus
Rhipiceridae Oligorhipis CSR049
Dascillidae Dascillus INB028
Buprestidae Trachys INB084
Buprestidae Coroebus INB018
Byrrhidae Notolioon CSR008
Byrrhidae Cytilus INB053
Eulichadidae Eulichas INB025
Callirhipidae Ennometes CSR009
Ptilodactylidae Ptilodactyla INB015
Dryopidae Helichus CSR019
Psephenidae Mataeopsephus INB081
Chelonariidae Chelonarium CSR012
Elmidae Stenelmis INB038
Limnichidae Limnichus TH1
Heteroceridae Heterocerus INB092
Omethidae Drilonius INB129
Artematopodidae Artematopus CSR004
Throscidae Trixagus INB125
Eucnemidae Anischia CSR022
Cantharidae Fissocantharis INB156
Lycidae Lycostomus YL0616
Elateridae Agrypnus INB160
Lampyridae Luciola INB061
Rhagophthalmidae Rhagophthalmus CSR132
Phengodidae Stenophrixothrix CSR041
Nosodendridae Nosodendron CSR038
Hydrophilidae Anacaena INB105
Histeridae Platysoma CSR023
Leiodidae Agathidium CSR032
Agyrtidae Pteroloma INB201
Staphylinidae Staphylinus INB008
Silphidae Necrodes INB078
Jacobsoniidae Derolathrus CSR026
Ptiliidae sp CSR044
Hydraenidae Hydraena INB108
Lucanidae Aegus INB148
Trogidae Trox INB036
Trogidae Omorgus CSR057
Geotrupidae Geotrupes INB097
Glaphyridae Amphicoma INB011
Scarabaeidae Copris INB146
Hybosoridae Cyphopisthes CSR138
Dermestidae Dermestes CSR089
Ptinidae Ptinus CSR045
Bostrichidae Lyctus CSR070
Latridiidae Corticaria CSR029
Alexiidae Sphaerosoma CSR002
Endomychidae Encymon CSR097
Coccinellidae Harmonia C10
Anamorphidae Papuella CSR095
Bothrideridae Ascetoderes CSR074
Murmidiidae Murmidius CSR078
Discolomatidae Aphanocephalus CSR018
Teredidae Teredolaemus CSR073
Euxestidae Hypodacnella CSR010
Cerylonidae Ostomopsis CSR079
Megalopodidae Temnaspis INB031
Cerambycidae Dorysthenes INB150
Chrysomelidae Galerucinae sp INB140
Chrysomelidae Altica INB141
Nemonychidae Aragomacer CSR120
Anthribidae Acorynus INB007
Belidae Rhinotia CSR067
Attelabidae Byctiscus INB076
Curculionidae Curculio INB184
Brentidae Apion CSR075
Monotomidae Rhizophagus CSR119
Hobartiidae Hydnobioides CSR106
Nitidulidae Brachypeplus CSR121
Kateretidae Notobrachypterus CSR027
Cybocephalidae Cybocephalus CSR088
Erotylidae Anadastus CSR100
Boganiidae Paracucujus CSR006
Protocucujidae Ericmodes CSR129
Helotidae Neohelota INB004
Silvanidae Uleiota CSR052
Cryptophagidae Curelius INB192
Phloeostichidae Hymaea CSR127
Myraboliidae Myrabolia CSR037
Passandridae Passandra CSR125
Phalacridae Phalacrinus CSR126
Propalticidae Propalticus CSR042
Laemophloeidae Laemophloeus CSR110
Trogossitidae Rentonellum CSR059
Byturidae Haematoides INB039
Biphyllidae Biphyllus CSR068
Acanthocnemidae Acanthocnemus CSR062
Trogossitidae Leperina CSR160
Trogossitidae Thymalus INB024
Trogossitidae Larinotus CSR058
Thanerocleridae Isoclerus CSR055
Cleridae Necrobia CSR084
Prionoceridae Idgia INB040
Melyridae Carphurus CSR117
Trogossitidae Parapeltis CSR159
Trogossitidae Ancyrona CSR158
Lymexylidae Melittomma CSR033
Melandryidae Melandryinae sp INB203
Incertae sedis Rhizonium CSR105
Ciidae Cis CSR013
Zopheridae Zopherosis CSR163
Zopheridae Bitoma CSR061
Tenebrionidae Adelium CSR149
Mycetophagidae Mycetophagus INB047
Archeocrypticidae Wattianus CSR066
Ulodidae Ulodes CSR161
Ulodidae Meryx CSR162
Oedemeridae Ditylus INB033
Mycteridae Trichosalpingus CSR036
Scraptiidae Scraptia INB205
Aderidae sp INB206
Ripiphoridae Rhipidioides CSR133
Ischaliidae Ischalia INB049
Salpingidae Ocholissa CSR136
Pythidae Anaplopus CSR131
Trictenotomidae Trictenotoma INB208
Boridae Synercticus CSR069
Pyrochroidae Morpholycus CSR047
Meloidae Epicauta INB093
Anthicidae Anthicus CSR063

**Figure S5.** Test of model-depenmdent phylogenetic signal on a reduced 138 taxon dataset analysed with the C10 model in IQ-TREE.

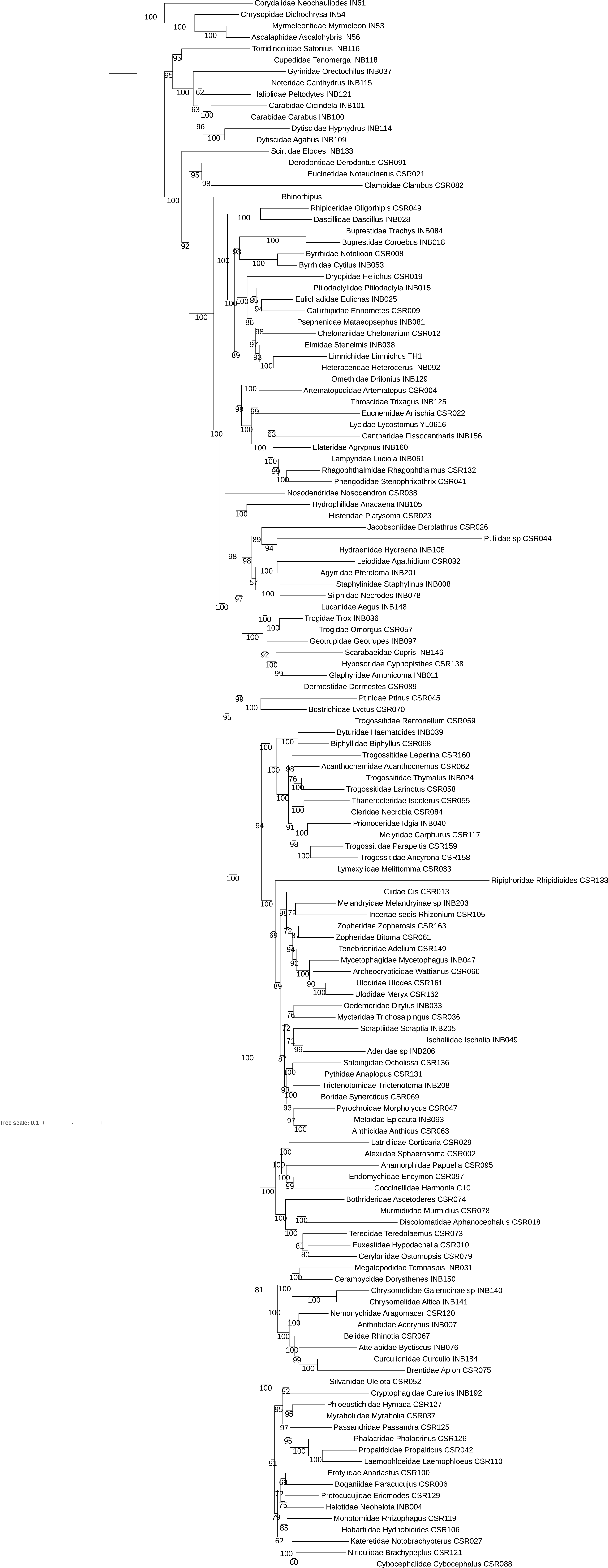
Corydalidae Neochauliodes IN61
Chrysopidae Dichochrysa IN54
Myrmeleontidae Myrmeleon IN53
Ascalaphidae Ascalohybris IN56
Torridincolidae Satonius INB116
Cupedidae Tenomerga INB118
Gyrinidae Orectochilus INB037
Noteridae Canthydrus INB115
Haliplidae Peltodytes INB121
Carabidae Cicindela INB101
Carabidae Carabus INB100
Dytiscidae Hyphydrus INB114
Dytiscidae Agabus INB109
Scirtidae Elodes INB133
Derodontidae Derodontus CSR091
Eucinetidae Noteucinetus CSR021
Clambidae Clambus CSR082
Rhinorhipus
Rhipiceridae Oligorhipis CSR049
Dascillidae Dascillus INB028
Buprestidae Trachys INB084
Buprestidae Coroebus INB018
Byrrhidae Notolioon CSR008
Byrrhidae Cytilus INB053
Dryopidae Helichus CSR019
Ptilodactylidae Ptilodactyla INB015
Eulichadidae Eulichas INB025
Callirhipidae Ennometes CSR009
Psephenidae Mataeopsephus INB081
Chelonariidae Chelonarium CSR012
Elmidae Stenelmis INB038
Limnichidae Limnichus TH1
Heteroceridae Heterocerus INB092
Omethidae Drilonius INB129
Artematopodidae Artematopus CSR004
Throscidae Trixagus INB125
Eucnemidae Anischia CSR022
Lycidae Lycostomus YL0616
Cantharidae Fissocantharis INB156
Elateridae Agrypnus INB160
Lampyridae Luciola INB061
Rhagophthalmidae Rhagophthalmus CSR132
Phengodidae Stenophrixothrix CSR041
Nosodendridae Nosodendron CSR038
Hydrophilidae Anacaena INB105
Histeridae Platysoma CSR023
Jacobsoniidae Derolathrus CSR026
Ptiliidae sp CSR044
Hydraenidae Hydraena INB108
Leiodidae Agathidium CSR032
Agyrtidae Pteroloma INB201
Staphylinidae Staphylinus INB008
Silphidae Necrodes INB078
Lucanidae Aegus INB148
Trogidae Trox INB036
Trogidae Omorgus CSR057
Geotrupidae Geotrupes INB097
Scarabaeidae Copris INB146
Hybosoridae Cyphopisthes CSR138
Glaphyridae Amphicoma INB011
Dermestidae Dermestes CSR089
Ptinidae Ptinus CSR045
Bostrichidae Lyctus CSR070
Trogossitidae Rentonellum CSR059
Byturidae Haematoides INB039
Biphyllidae Biphyllus CSR068
Trogossitidae Leperina CSR160
Acanthocnemidae Acanthocnemus CSR062
Trogossitidae Thymalus INB024
Trogossitidae Larinotus CSR058
Thanerocleridae Isoclerus CSR055
Cleridae Necrobia CSR084
Prionoceridae Idgia INB040
Melyridae Carphurus CSR117
Trogossitidae Parapeltis CSR159
Trogossitidae Ancyrona CSR158
Lymexylidae Melittomma CSR033
Ripiphoridae Rhipidioides CSR133
Ciidae Cis CSR013
Melandryidae Melandryinae sp INB203
Incertae sedis Rhizonium CSR105
Zopheridae Zopherosis CSR163
Zopheridae Bitoma CSR061
Tenebrionidae Adelium CSR149
Mycetophagidae Mycetophagus INB047
Archeocrypticidae Wattianus CSR066
Ulodidae Ulodes CSR161
Ulodidae Meryx CSR162
Oedemeridae Ditylus INB033
Mycteridae Trichosalpingus CSR036
Scraptiidae Scraptia INB205
Ischaliidae Ischalia INB049
Aderidae sp INB206
Salpingidae Ocholissa CSR136
Pythidae Anaplopus CSR131
Trictenotomidae Trictenotoma INB208
Boridae Synercticus CSR069
Pyrochroidae Morpholycus CSR047
Meloidae Epicauta INB093
Anthicidae Anthicus CSR063
Tree scale: 0.1
Latridiidae Corticaria CSR029
Alexiidae Sphaerosoma CSR002
Anamorphidae Papuella CSR095
Endomychidae Encymon CSR097
Coccinellidae Harmonia C10
Bothrideridae Ascetoderes CSR074
Murmidiidae Murmidius CSR078
Discolomatidae Aphanocephalus CSR018
Teredidae Teredolaemus CSR073
Euxestidae Hypodacnella CSR010
Cerylonidae Ostomopsis CSR079
Megalopodidae Temnaspis INB031
Cerambycidae Dorysthenes INB150
Chrysomelidae Galerucinae sp INB140
Chrysomelidae Altica INB141
Nemonychidae Aragomacer CSR120
Anthribidae Acorynus INB007
Belidae Rhinotia CSR067
Attelabidae Byctiscus INB076
Curculionidae Curculio INB184
Brentidae Apion CSR075
Silvanidae Uleiota CSR052
Cryptophagidae Curelius INB192
Phloeostichidae Hymaea CSR127
Myraboliidae Myrabolia CSR037
Passandridae Passandra CSR125
Phalacridae Phalacrinus CSR126
Propalticidae Propalticus CSR042
Laemophloeidae Laemophloeus CSR110
Erotylidae Anadastus CSR100
Boganiidae Paracucujus CSR006
Protocucujidae Ericmodes CSR129
Helotidae Neohelota INB004
Monotomidae Rhizophagus CSR119
Hobartiidae Hydnobioides CSR106
Kateretidae Notobrachypterus CSR027
Nitidulidae Brachypeplus CSR121
Cybocephalidae Cybocephalus CSR088

**Figure S6.** Phylogeny of Coleoptera inferred with the site-heterogeneous Bayesian model CAT-GTR+G4 implemented in PhyloBayes.

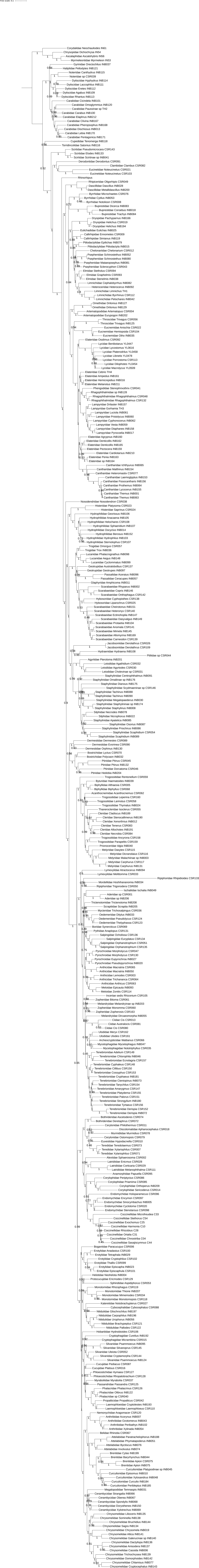

**Figure S7.** Phylogeny of Coleoptera inferred with the site-homogeneous ML model GTR20 implemented in IQ-TREE.

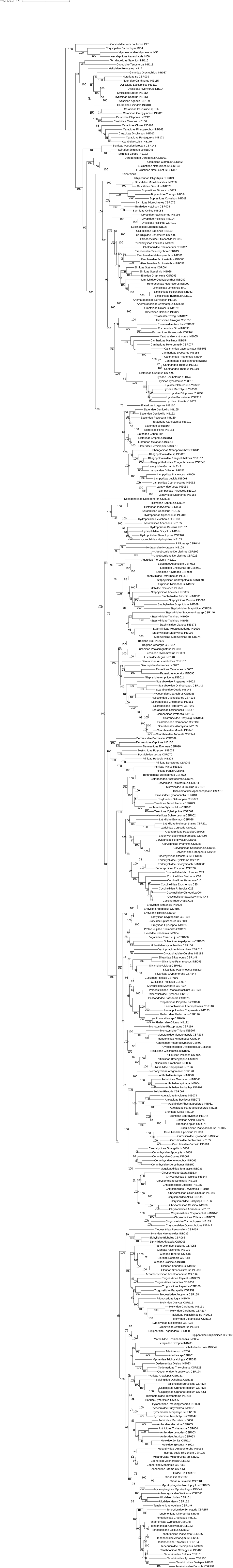
Tree scale: 0.1

**Figure S8.** Phylogeny of Coleoptera inferred with the site-homogeneous ML model LG+C20 implemented in IQ-TREE.

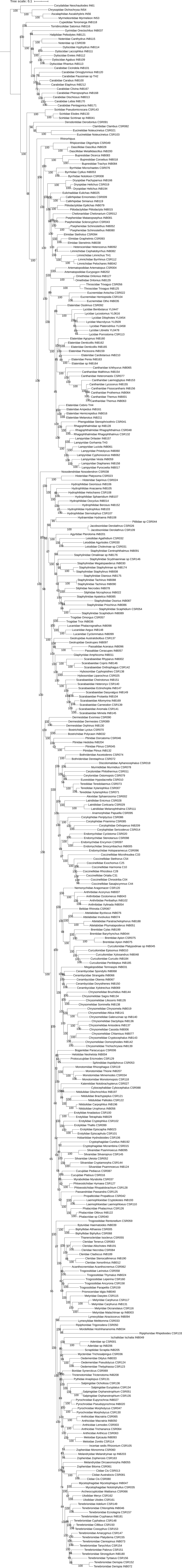
Tree scale: 0.1

**Figure S9.** Dated phylogenetic tree of Coleoptera; uniform priors, autocorrelated rate clock.

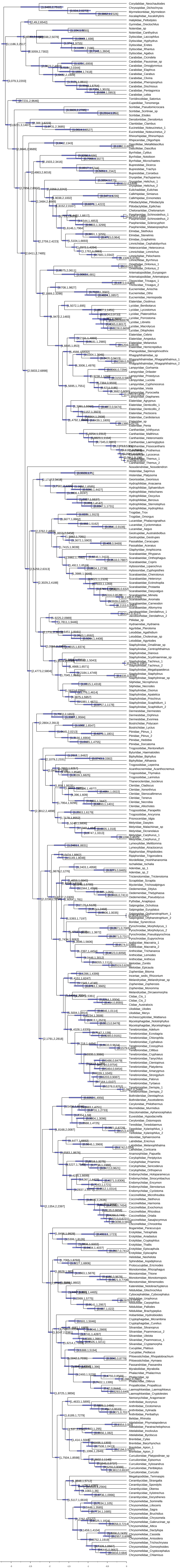

**Figure S10.** Dated phylogenetic tree of Coleoptera; uniform priors, independent rate clock.

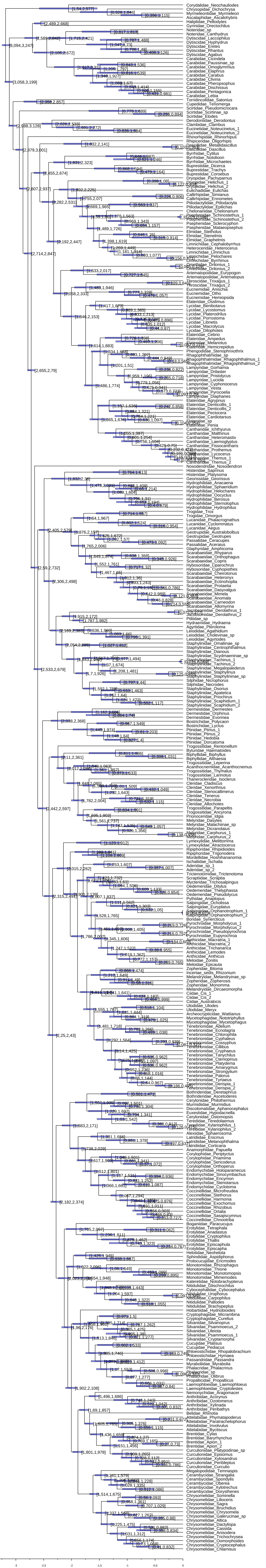

**Figure S11. P**hylogenetic tree of Coleoptera displaying the effective priors.

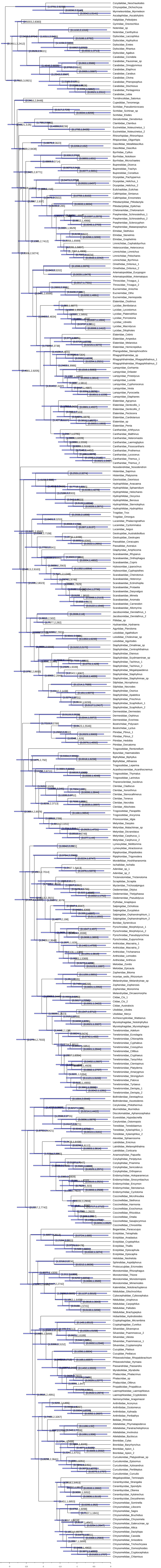

**Table S1.** Fit of maximum likelihood (ML) models to the reduced coleopteran dataset for model sensitivity testing, analysed in ModelFinder. Abbreviations: AIC, Akaike information criterion scores.

| Model | AIC |
|---|---|
| C10 | 1035697.227 |
| GTR20 | 1210678.781 |
| LG | 1211046.717 |
| JTT | 1215087.892 |
| WAG | 1220238.554 |

**Table S2.** Overview of fossil calibrations for molecular clock analyses.

| Node | Clade | Fossil constraint | Minimum age (Ma) | Maximum age (Ma) |
|---|---|---|---|---|
| 1 | (Megaloptera + Neuroptera) – Coleoptera | *Stephanastus polinae* Kirejtshuk and Nel, 2013 | 298.75 | 324 |
| 2 | Crown Coleoptera | *Ponomarenkium belmonthensis* Yan, Lawrence, Beattie and Beutel, 2018 | 251.878 | 307.1 |
| 3 | Gyrinidae – Haliplidae | *Tunguskagyrus planus* Yan, Beutel and Lawrence, 2018 | 251.878 | 293.69 |
| 4 | Dytiscidae – Noteridae | *Palaeodytes gutta* Ponomarenko, 1987 | 155 | 251.878 |
| 5 | Cupedidae – Torridincolidae | *Kirghizocupes cellulosus* Ponomarenko, 1966 | 235 | 293.69 |
| 6 | Crown Polyphaga | undescribed Elateriformia, Ladinian of Tongchuan, China | 237 | 293.69 |
| 7 | Derodontidae – (Clambidae + Eucinetidae) | *Juropeltastica sinica* Cai, Lawrence and Huang, 2014 | 165 | 251.878 |
| 8 | Clambidae – Eucinetidae | *Mesocinetus aequalis* Kirejtshuk and Ponomarenko, 2010 | 145 | 235 |
| 9 | Rhipiceridae – Dascillidae | *Parelateriformius communis Yan and Wang, 2010* | 165 | 235 |
| 10 | Buprestidae – Byrrhidae | *Sinoparathyrea bimaculata* Pan, Chang and Ren, 2011 | 165 | 235 |
| 11 | Dryopoidea – Byrrhoidea + Buprestoidea | undescribed Dryopidae, Aptian Crato Formation, Brazil | 112.6 | 165 |
| 12 | Crown Psephenidae | *Vetujinbrianax cretaceus* Cai and Huang, 2018 | 98.17 | 165 |

| | | | | |
|---|---|---|---|---|
| 13 | Elmidae – (Limnichidae + Heteroceridae) | *Elmadulescens rugosus* Peris, Maier and Sánchez-García, 2015 | 105 | 165 |
| 14 | Crown Heteroceridae | Heterocerites magnus Prokin and Ren, 2011 | 122.2 | 165 |
| 15 | Artematopodidae – Omethidae | *Sinobrevipogon jurassicus* Cai, Lawrence, Ślipiński and Huang, 2015 | 165 | 235 |
| 16 | Throscidae – Eucnemidae | *Potergosoma gratiosum* Kovalev and Kirejtshuk, 2013 | 125 | 235 |
| 17 | *Osslimus* (Elateridae) – Lycidae | *Burmolycus compactus* Bocak, Li and Ellenberger, 2019 | 98.17 | 165 |
| 18 | Lampyridae – Phengodidae + Rhagophthalmidae | *Protoluciola albertalleni* Kazantsev, 2015 | 98.17 | 165 |
| 19 | Crown Cantharidae | *Molliberus albae* Peris and Fanti, 2018 | 105 | 165 |
| 20 | Georissidae – Hydrophilidae | *Protochares brevipalpis* Fikáček, Prokin, Yan, Yue, Wang, Ren and Beattie, 2014 | 147.28 | 235 |
| 21 | Trogidae – Lucanidae | *Juraesalus atavus* Nikolajev, Wang, Liu and Zhang, 2011 | 165 | 235 |
| 22 | Passalidae – Geotrupidae | *Parageotrupes incanus* Nikolajev and Ren, 2010 | 122.2 | 165 |
| 23 | Hybosoridae – Scarabaeidae | *Protohybosorus grandissimus* Nikolajev, 2010 | 155 | 235 |
| 24 | Ptiliidae – Hydraenidae | *Ochtebiites minor* Ponomarenko, 1985 | 182 | 235 |
| 25 | Agyrtidae – Leiodidae | *Mesecanus communis* Ponomarenko, 1977 | 165 | 235 |

| | | | | |
|---|---|---|---|---|
| 26 | Crown Staphylinidae | undescribed Silphidae, Haifanggou Formation, China | 165 | 235 |
| 27 | *Megalopaederus* (Paederinae) – Staphylininae | *Megolisthaerus minor* Cai and Huang, 2013 | 122.2 | 165 |
| 28 | Silphinae – Staphylinidae *partim* | *Mesecanus communis* Ponomarenko, 1977 | 165 | 235 |
| 29 | Staphylinidae: Apateticinae – Osoriinae | *Mesallotrochus longiantennatus* Cai and Huang, 2015 | 98.17 | 165 |
| 30 | Crown Bostrichoidea | *Paradermestes jurassicus* Deng, Ślipiński, Ren and Pang, 2017 | 165 | 235 |
| 31 | Crown Bostrichidae | *Poinarinius burmaensis* Legalov, 2018 | 98.17 | 165 |
| 32 | Crown Ptinidae | *Actenobius magneoculus* Peris, Philips, and Delclòs, 2015 | 105 | 165 |
| 33 | Thanerocleridae – Cleridae | *Wangweiella calloviana* Kolibáč and Huang, 2016 | 165 | 235 |
| 34 | Prionoceridae – Melyridae | *Idgiaites jurassicus* Liu, Ślipiński, Leschen, Ren and Pang, 2015 | 165 | 235 |
| 35 | Lymexylidae | *Cratoatractocerus grimaldii* Wolf, 2011 | 112.6 | 235 |
| 36 | Crown Tenebrionoidea | *Archaeoripiphorus nuwa* Hsiao, Yu and Deng, 2017 | 165 | 235 |
| 37 | Rhipiphoridae – Mordellidae | *Protoripidius burmiticus* Cai, Yin and Huang, 2018 | 98.17 | 235 |
| 38 | Ischaliidae – Aderidae | undescribed Aderidae from Cretaceous Lebanese amber | 125 | 165 |

| | | | | |
|---|---|---|---|---|
| 39 | Salpingidae – Pythidae | *Eopeplus stetzenkoi* Kirejtshuk and Nel, 2009 | 53 | 165 |
| 40 | Anthicidae – Meloidae | *Camelomorpha longicervix* Kirejtshuk, Azar and Telnov, 2008 | 125 | 165 |
| 41 | Crown Zopheridae | *Paleoendeitoma antennata* Deng, Ślipiński, Ren and Pang, 2017 | 98.17 | 165 |
| 42 | Crown Tenebrionidae | *Alphitopsis initialis* Kirejtshuk, Nabozhenko and Nel, 2011 | 122.2 | 165 |
| 43 | Alexiidae – Latridiidae | *Tetrameropsis mesozoica* Kirejtshuk and Azar, 2008 | 125 | 165 |
| 44 | Crown Endomychidae | *Palaeomycetes foveolatus* Tomaszewska, Ślipiński and Ren, 2018 | 98.17 | 165 |
| 45 | Crown Coccinellidae | *Rhyzobius antiquus* Kirejtshuk and Nel, 2012 | 53 | 125 |
| 46 | Boganiidae – Erotylidae | *Palaeoboganium jurassicum* Liu, Ślipiński, Lawrence, Ren and Pang, 2018 | 165 | 235 |
| 47 | Helotidae – (Protocucujidae + Sphindidae) | *Libanopsis poinari* Kirejtshuk, 2015 | 125 | 235 |
| 48 | Monotomidae – (Kateretidae + Nitidulidae) | *Jurorhizophagus alienus* Cai, Ślipiński and Huang, 2015 | 165 | 235 |
| 49 | Crown Nitidulidae | *Crepuraea archaica* Kirejtshuk, 1990 | 119 | 165 |
| 50 | Crown Silvanidae | *Protoliota antennatus* Liu, Ślipiński, Wang and Pang, 2019 | 98.17 | 165 |
| 51 | Nemonychidae – Anthribidae | *Protoscelis longicornis* (Medvedev, 1968) | 155 | 235 |

| | | | | |
|---|---|---|---|---|
| 52 | Belidae – (Attelabidae, (Brentidae, Curculionidae)) | *Sinoeuglypheus daohugouensis* Yu, Davis and Shih, 2019 | 165 | 235 |
| 53 | Brentidae – Curculionidae | *Axelrodiellus ruptus* Zherikhin and Gratshev, 2004 | 112.6 | 165 |
| 54 | Crown Chrysomelidae | *Mesolpinus antennatus* Kirejtshuk, Moseyko and Ren, 2015 | 122.2 | 165 |
| 55 | Crown Cerambycidae | *Cretoprionus liutiaogouensis* Wang, Ma, McKenna, Yan, Zhang and Jarzembowski, 2014 | 122.2 | 165 |
| 56 | Sagrinae – Bruchinae | *Mesopachymerus antiqua* Poinar, 2005 | 78 | 155 |
| 57 | Galerucinae – Chrysomelinae | *Taimyraltica calcarata* Nadein, 2018 | 83.4 | 165 |

Cai, C. and Huang, D. First fossil Coloninae from Upper Cretaceous Burmese amber (Coleoptera:

Staphylinoidea: Leiodidae). *Cretaceous Res.* 77: 69-74 (2017).
Cai, C. and Huang, D. First Mesozoic water-penny beetles from Upper Cretaceous Burmese amber (Coleoptera: Psephenidae). *Cretaceous Res*. 91, 202–207 (2018).
Cai, C., Yin, Z, and Huang, D. A new ripiphorid beetle from Upper Cretaceous Burmese amber sheds light on early evolution of the extant subfamily Ripidiinae (Coleoptera: Ripiphoridae). *Compt. Rend. Palevol* 17, 351–356 (2018a).
Cai, C. *et al.* Beetle pollination of cycads in the Mesozoic. *Curr. Biol.* 28, 2806–2812 (2018b).
Cai, C. and Huang, D. Sexual dimorphism in mid-Cretaceous silvanid beetles from northern Myanmar (Coleoptera, Silvanidae, Brontinae). *Palaeoentomol*. 2, 289–296 (2019).
Cai, C., Fu, Y. and Huang, D. A large artematopodid beetle (Coleoptera: Elateroidea: Artematopodidae) from the Early Cretaceous of China and its systematic position. *Cretaceous Res*. 105, 103986 (2020).
Caterino, M. S., Hunt, T. and Vogler, A. P. On the constitution and phylogeny of Staphyliniformia (Insecta: Coleoptera). *Mol. Phylogenet. Evol*. 34, 655–672 (2005).
Chandler D. S. Anthicidae Latreille, 1819. in *Handbook of Zoology, Arthropoda: Insecta, Coleoptera, Beetles, Vol. 2: Morphology and Systematics (Elateroidea, Bostrichiformia, Cucujiformia partim)* (eds. Leschen, R. A. B., Beutel, R. G. and Lawrence, J. F.) 729–741 (De Gruyter, Berlin and Boston, MA, 2010).
Chang, S., Zhang, H., Renne, P. R. and Fang, Y. High-precision 40Ar/39Ar age constraints on the basal Lanqi Formation and its implications for the origin of angiosperm plants. *Earth Planet. Sci. Lett*. 279, 212–221 (2009).
Chatzimanolis, S., Grimaldi, D. A., Engel, M. S. and Fraser, N. C. *Leehermania prorova*, the earliest staphyliniform beetle, from the Late Triassic of Virginia (Coleoptera: Staphylinidae). *Am. Mus. Novit.* 3761, 1–28 (2012).
Chen, W. *et al*. Isotope geochronology of the fossil-bearing beds in the Daohugou area, Ningcheng, Inner Mongolia. *Geol. Bull. Chin*. 23, 1165–1169 (2004).
Clarke, R. O. S. *Handbooks for the Identification of British Insects, Vol. 5, Part 2c: Coleoptera. Heteroceridae*. (Royal Entomological Society of London, London, 1973).
Clements, T., Purnell, M. and Gabbott, S. The Mazon Creek Lagerstätte: a diverse late Paleozoic ecosystem entombed within siderite concretions. *J. Geol. Soc.* 176, 1–11 (2019).
Cline, A. R. *et al*. Molecular phylogeny of Nitidulidae: assessment of subfamilial and tribal classification and formalization of the family Cybocephalidae (Coleoptera: Cucujoidea). *Syst. Entomol.* 39, 758–772 (2014).
Costa, C., Vanin, S. A. and Ide, S. Systematics and bionomics of Cneoglossidae with a cladistic analysis of Byrrhoidea *sensu* Lawrence and Newton (1995) (Coleoptera, Elateriformia). *Arq. Zool. São Paulo* 35, 231–300 (1999).
Crowson, R. A. *The Natural Classification of the Families of Coleoptera* (Nathaniel Lloyd, London, (1955).
Crowson, R. A. Studies on the Dermestoidea (Coleoptera), with special reference to the New Zealand fauna. *Trans. R. Entomol. Soc. London* 111, 81–94 (1959).
Crowson, R. A. The phylogeny of Coleoptera. *Annu. Rev. Entomol*. 5, 111–134 (1960).
Crowson, R. A. On a new superfamily Artematopoidea of polyphagan beetles, with the definition of two new fossil genera from the Baltic amber. *J. Nat. Hist.* 7, 225–238 (1973).
Crowson, R. A. The evolutionary history of Coleoptera, as documented by fossil and comparative evidence. *Atti del X Congr- Nazion. Ital. Entomol.* 1974, 47–90 (1975).
Crowson, R. A. Problems of phylogenetic relationships in Dryopoidea (Coleoptera). *Entomol. Germ.* 4, 250–257 (1978).

Crowson, R. A. On the dryopoid affinities of Buprestidae. *Coleopt. Bull.* 36, 22–25 (1982).
Crowson, R. A. A new genus of Boganiidae (Coleoptera) from Australia, with observations on glandular openings, cycad associations and geographical distribution of the family. *Austr. J. Entomol.* 29, 91–99 (1990).
Crowson, R.A. The relations of Coleoptera to Cycadales. in *Advances in Coleopterology* (ed. Zunino, M., Bellés, X. and Blas, M.) 13–28 (Asociación Europea de Coleopterologia, 1991).
Crowson, R. A. Some interesting evolutionary parallels in Coleoptera. in *Biology, Phylogeny, and Classification of Coleoptera: Papers Celebrating the 80th Birthday of Roy A. Crowson* (eds. Pakaluk, J. and Ślipiński, S. A.) 63–85 (Muzeum i Instytut Zoologii PAN, 1995).
Deng, C., Ślipiński, A., Ren, D. and Pang, H. The oldest dermestid beetle from the Middle Jurassic of China (Coleoptera: Dermestidae). *Ann. Zool*. 67, 109–112 (2017a).
Deng, C., Ślipiński, A., Ren, D. and Pang, H. The first Mesozoic colydiid beetles (Coleoptera: Zopheridae: Colydiinae) from the Upper Cretaceous amber of Myanmar. *Cretaceous Res*. 78, 71–77 (2017b).
Dobruskina, I. A. Keuper (Triassic) flora from Middle Asia (Madygen, Southern Fergana). *New Mexico Mus. Nat. Hist. Sci Bull.* 5, 1–49 (1995).
Doludenko, M. P. and Orlovskaya, E. R. Jurassic floras of the Karatau Range, southern Kazakhstan. *Palaeontol*. 19, 627–640 (1976).
Drilling, K., Dettner, K. and Klass, K. D. The distribution and evolution of exocrine compound glands in Erotylinae (Insecta: Coleoptera: Erotylidae). *Ann. Soc. Entomol. Fr.* 49, 36–52 (2013).
Dvořák, T., Pecharová, M., Krzemiński, W. and Prokop, J. New archaeorthopteran insects from the Carboniferous of Poland: Insights into tangled taxonomy. *Acta Palaeontol. Pol*. 64, 787–796 (2019).
Endrödy-Younga, S. Nosodendridae, a new beetle family for the African continent (Coleoptera: Byrrhoidea). *Ann. Transvaal Mus.* 35, 61–66 (1989).
Escalona, H. E., Lawrence, J. F., Wanat, M. and Ślipiński, A. Phylogeny and placement of Boganiidae (Coleoptera, Cucujoidea) with a review of Australian and New Caledonian taxa. *Syst. Entomol.* 40, 628–651 (2015).
Español, F. *Fauna Ibérica Coleoptera Anobiidae, Volume 2* (Museo Nacional de Ciencias Naturales, Madrid, 1992).
Falin, Z. H. and Engel, M. S. Notes on Cretaceous Ripidiini and revised diagnoses of the Ripidiinae, Ripidiini, and Eorhipidiini (Coleoptera: Ripiphoridae). *Alavesia* 3, 35–42 (2010).
Fikáček, M. *et al.* Phylogeny and the fossil record of the Helophoridae reveal Jurassic origin of extant hydrophiloid lineages (Coleoptera: Polyphaga). *Syst. Entomol.* 37, 420–447 (2012).
Fikáček, M. *et al.* Modern hydrophilid clades present and widespread in the Late Jurassic and Early Cretaceous (Coleoptera: Hydrophiloidea: Hydrophilidae). *Zool. J. Linn. Soc*. 170, 710–734 (2014).
Fikáček, M. *et al.* Reliable placement of beetle fossils via phylogenetic analyses–Triassic *Leehermania* as a case study (Staphylinidae or Myxophaga?). *Syst. Entomol.* 45, 175–187 (2020).
Forbes, W. T. M. The wing folding patterns of the Coleoptera. *J. N. Y. Entomol. Soc*. 34, 42–68, 91–138 (1926).
Forbes, W.T.M. The Protocoleoptera. *Psyche* 35, 32–35 (1928).
Forbes, W. T. M. The wing of Schizopini (Coleoptera: Dascillidae). *Entomol. News* 53, 101–102 (1942).
Friedrich, F. and Beutel, R. G. The pterothoracic skeletomuscular system of Scirtoidea (Coleoptera: Polyphaga) and its implications for the high-level phylogeny of beetles. J. *Zool. Syst. and Evol.*

*Res*. 44, 290–315 (2006).
Gao, K. and Ren, D. Radiometric dating of ignimbrite from Inner Mongolia provides no indication of a post-middle Jurassic age for the Daohugou beds. *Acta Geol. Sin*. 80, 42–45 (2006).
Grebennikov, V. V. and Scholtz, C. H. The basal phylogeny of Scarabaeoidea (Insecta: Coleoptera) inferred from larval morphology. *Invert. Syst*. 18, 321–348 (2004).
Grebennikov, V. V. and Newton, A. F. Good-bye Scydmaenidae, or why the ant-like stone beetles should become megadiverse Staphylinidae sensu latissimo (Coleoptera). *Eur. J. Entomol.* 106, 275–301 (2009).
Grebennikov, V. V. and Newton, A. F. Detecting the basal dichotomies in the monophylum of carrion and rove beetles (Insecta: Coleoptera: Silphidae and Staphylinidae) with emphasis on the Oxyteline group of subfamilies. *Arthropod Syst. Phylogenet.* 70, 133-165 (2012).
Grimaldi, D. A., Engel, M. S. and Nascimbene, P. C. Fossiliferous Cretaceous amber from Myanmar (Burma): its rediscovery, biotic diversity, and paleontological significance. *Am. Mus. Novit.* 3361, 1–71 (2002).
Grimaldi, D. and Engel, M. S. *Evolution of the Insects* (Cambridge University, 2005).
Hansen, M. Phylogeny and classification of the staphyliniform beetle families (Coleoptera). *Biol. Skrifter. Dan. Vid. Selsk.* 48, 1–339 (1997).
Hatch, M. H. Studies on the carrion beetles of Minnesota, including new species. *Tech. Bull. Univ. Minn. Agr. Exp. Stat*. 48, 1–19 (1927).
Hayes, W. P. and Chu, H. F. The larvae of the genus *Nosodendron* Latr. (Coleoptera, Nosodendridae). *Ann. Entomol. Soc. Am.* 39, 69–79 (1946).
He, H. Y. *et al*. 40Ar/39Ar dating of ignimbrite from Inner Mongolia, northeastern China, indicates a post Middle Jurassic age for the overlying Daohugou Bed. *Geophys. Res. Lett*. 31, L20609 (2004).
Hinton, H. E. An inquiry into the natural classification of the Dryopoidea, based partly on a study of their internal anatomy (Col.). *Trans. Roy. Entomol. Soc.* 89, 133–184 (1939).
Holloway, B. A. Taxonomy and phylogeny in the Lucanidae (Insecta: Coleoptera) *Rec. Dom. Museum* 3, 321–365 (1960).
Holloway, B. A. Further studies on generic relationships in Lucanidae (Insecta: Coleoptera) with special reference to the ocular canthus *New Zeal. J. Sci.* 12, 958–977 (1969).
Hong, Y. C. A new Early Cretaceous beetle family—Magnocoleidae fam. nov. (Insecta: Coleoptera) in Hebei Province. *Geosci. J. Grad. Sch., China Univ. Geosci.* 12, 40–48 (1998).
Hsiao, Y., Yu, Y., Deng, C. and Pang, H. The first fossil wedge-shaped beetle (Coleoptera, Ripiphoridae) from the middle Jurassic of China. *Eur. J. Taxon.* 277, 1–13 (2017).
Huang, D. *The Daohugou Biota*. (Shanghai Science and Technology, 2016) [in Chinese].
Hunt, T. *et al*. A comprehensive phylogeny of beetles reveals the evolutionary origins of a superradiation. *Science* 318, 1913–1916 (2007).
Jäch, M. A., Beutel, R. G., Delgado, J. A. and Díaz, J. A. *Hydraenidae Mulsant, 1844*. in *Handbook of Zoology, Arthropoda: Insecta, Coleoptera, Beetles, Vol. 1: Morphology and Systematics (Archostemata, Adephaga, Myxophaga, Polyphaga partim), 2nd ed.* (eds. Beutel, R. G., Leschen, R. A. B.) 316–345 (Walter de Gruyter, Berlin and Boston, MA, 2016).
Jameson, M. L. Hybosoridae Erichson 1847. in *American Beetles. Volume 2* (eds. Arnett, R. H., Thomas, M. C., Skelley, P. E. and Frank J. H.) 32–33 (CRC Press, Boca Raton, 2002).
Jeannel, R. Monographie des Catopidae (Insectes Coléoptères). *Mem. Mus. Natl. Hist. Natur. (N.S.)* 1, 1–433 (1936).
Jeannel, R. and Paulian, R. Morphologic abdominale des Coléoptères et systematique de l’ordre. *Rev. Fr. Entomol.* 11, 65–109 (1944).

Jin, Z., Escalona, H. E., Ślipiński, A. and Pang, H. Phylogeny and classification of Rhipicerinae (Coleoptera: Rhipiceridae) with a review of the Australian taxa. *Ann. Zool.* 63, 275–317 (2013a).

Jin, Z., Ślipiński, A., Pang, H. and Ren, D. A new Mesozoic species of soft-bodied plant beetle (Coleoptera: Dascillidae) from the Early Cretaceous of Inner Mongolia, China with a review of fossil Dascillidae. *Ann. Zool.* 63, 501–509 (2013b).

Jirásek, J. *et al.* The Main Ostrava Whetstone: composition, sedimentary processes, palaeogeography and geochronology of a major Mississippian volcaniclastic unit of the Upper Silesian Basin (Poland and Czech Republic). *Int. J. Earth Sci.* 102, 989–1006 (2013).

John, H. Eine ubersicht uber die familie Propalticidae (Col.). *Pacific Insects* 2, 149–170 (1960).

Nadein, K. S. and Beždek, J. Galerucinae Latreille 1802. in *Handbook of Zoology, Arthropoda: Insecta, Coleoptera, beetles, Vol. 3: Morphology and Systematics (Phytophaga)* (eds. Leschen R.A.B. and Beutel, R. G.) 251–259 (De Gruyter, Berlin and Boston, MA, 2014).

Kasap, H. and Crowson, R. A. A comparative anatomical study of Elateriformia and Dascilloidea (Coleoptera). *Trans. Roy. Entomol. Soc. London* 126, 441–495 (1975).

Kazantsev, S. V. *Protoluciola albertalleni* gen.n., sp.n., a new Luciolinae firefly (Insecta: Coleoptera: Lampyridae) from Burmite amber. *Russ. Entomol. J.* 24, 281–283 (2015).

Kazantsev, S. V. *Protolycus gedaniensis* gen. et sp. nov., the first Baltic amber representative of Lycini (Coleoptera: Lycidae: Lycinae). *Palaeoentomol*. 2, 327–332 (2019).

Kim, S. I. and Farrell, B. D. Phylogeny of world stag beetles (Coleoptera: Lucanidae) reveals a Gondwanan origin of Darwin's stag beetle. *Mol. Phylogenet. Evol.* 86, 35–48 (2015).

King, J. G., Starr, J. R. and Lago, P. K. Molecular data resolves relationships within Heteroceridae (Coleoptera: Dryopoidea). *Syst. Entomol.* 36, 435–445 (2011).

Kirejtshuk, A. G. (1994) Parandrexidae Fam. n., Jurassic beetles of the Infraorder Cucujiformia (Coleoptera, Polyphaga). *Palaeontol. J.* 28, 69–78.

Kirejtshuk, A. G. A current generic classification of sap beetles (Coleoptera, Nitidulidae). *Zoosyst. Ross.* 17, 107–122 (2008).

Kirejtshuk, A. G. and Ponomarenko, A. G. Fossil beetles of the families Peltidae and Nitidulidae. *Paleontol. Zhur*. 2, 78–88 (1990).

Kirejtshuk A. G. and Azar D. New beetles of Polyphaga (Coleoptera, Polyphaga) from Lower Cretaceous Lebanese amber. *Denisia* 26, 119–130 (2009).

Kirejtshuk, A. G. and Nel, A. New genera and species of Cucujiformia (Coleoptera, Polyphaga) from lowermost Eocene French amber. *Denisia* 26, 103–118 (2009).

Kirejtshuk, A. G. and Ponomarenko, A. G. A new coleopterous family Mesocinetidae fam. nov. (Coleoptera: Scirtoidea) from Late Mesozoic and notes on fossil remains from Shar-Teg (Upper Jurassic, South-Western Mongolia). *Zoosyst. Ross.* 19, 301–325 (2010).

Kirejtshuk, A. G., Nabozhenko, M. V. and Nel, A. First Mesozoic representative of the subfamily Tenebrioninae (Coleoptera, Tenebrionidae) from the lower cretaceous of Yixian (China, Liaoning). *Entomol. Obozr.* 90, 548-552 (2011, in Russian), *Entomol. Rev*. 92, 97–100 (2012). (English translation).

Kirejtshuk, A. G and Nel, A. The oldest representatives of the family Coccinellidae (Coleoptera: Polyphaga) from the Lowermost Eocene Oise amber (France). *Zoosyst. Ross*. 21, 131–144 (2012).

Kirejtshuk, A. G. and Azar, D. Current knowledge of Coleoptera (Insecta) from the Lower Cretaceous Lebanese amber and taxonomical notes for some Mesozoic groups. *Terrest. Arthropod Rev.* 6, 103–134 (2013).

Kirejtshuk, A. G., Poschmann, M., Prokop, J., Garrouste, R. and Nel, A. Evolution of the elytral venation and structural adaptations in the oldest Palaeozoic beetles (Insecta: Coleoptera:

Tshekardocoleidae). *J. Syst. Palaeontol.* 12, 575–600 (2014).
Kirejtshuk, A. G. and Mantič, M. On systematics of the subfamily Cybocephalinae (Coleoptera: Nitidulidae) with description of new species and generic taxa. *Proceedings of the Zoological Institute RAS* 319, 196–214 (2015).
Kirejtshuk, A. G., Chetverikov, P. E. and Azar, D. Libanopsinae, new subfamily of the family Sphindidae (Coleoptera, Cucujoidea) from Lower Cretaceous Lebanese amber, with remarks on using confocal microscopy for the study of amber inclusions. *Cretaceous Res.* 52, 461–479 (2015a).
Kirejtshuk, A. G., Moseyko, A. G. and Ren, D. Mesozoic leaf beetles of the tribe Mesolpinini trib. nov. (Coleoptera, Chrysomelidae) from the Lower Cretaceous. *Paleontol. J.* 49, 1424–1435 (2015b).
Kirejtshuk, A. G. and Nel, A. Origin of the Coleoptera and significance of the fossil record. *Euroas. Entomol. J.* 15, 66–73 (2016).
Kirejtshuk, A. G., Nel, A. and Kirejtshuk, P. A. Taxonomy of the reticulate beetles of the subfamily Cupedinae (Coleoptera, Archostemata), with a review of the historical development. *Invert. Zool*. 13, 61–190 (2016).
Kirichkova, A. I. and Doludenko M. P. New Data on phytostratigraphy of Jurassic Deposits of Kazakhstan. *Stratigr. Geol. Korrelyatsiya* 4, 35–52 (1996).
Kolbe, H. Mein system der coleopteren. *Z. wiss. Insectbiol*. 4, 116–123, 153–162, 219–226, 246–251, 286–294, 389–400 (1908).
Kolibáč, J. Trogossitidae: A review of the beetle family, with a catalogue and keys. *ZooKeys* 366, 1–194 (2013).
Kolibáč, J. and Huang, D. The oldest known clerid fossils from the Middle Jurassic of China, with a review of Cleridae systematics (Coleoptera). *Syst. Entomol.* 41, 808–823 (2016).
Kovalev, V. G. The oldest representatives of the Diptera with short antennae from the Jurassic in Siberia. *Paleontol. J*. 15, 84–100 (1981).
Kovalev, A. V., Kirejtshuk, A. G. and Azar, D. The oldest representatives of the family Throscidae (Coleoptera: Elateriformia) from the Lower Cretaceous Lebanese amber. *Cretaceous Res*. 44, 157–165 (2013).
Krzemiński, W., Kania, I. and Azar, D. The Early Cretaceous evidence of rapid evolution of the genus *Helius* Lepeletier and Serville, 1828 (Limoniidae, Diptera). *Cretaceous Res*. 48, 96–101 (2014).
Kukalová, J. Protelytroptera from the Upper Permian of Australia, with a discussion of the Protocoleoptera and Paracoleoptera. *Psyche* 73, 89–111 (1966).
Kukalová-Peck, J. and Lawrence, J. F. Evolution of the hind wing in Coleoptera. *Can. Entomol.* 125, 181–258 (1993).
Kukalová-Peck, J. and Lawrence, J. F. Relationships among coleopteran suborders and major endoneopteran lineages: evidence from hind wing characters. *Eur. J. Entomol.* 101, 95–144 (2004).
Kukalová-Peck, J. and Beutel, R. G. Is the Carboniferous *Adiphlebia lacoana* really the "oldest beetle"? Critical reassessment and description of a new Permian beetle family. *Eur. J. Entomol.* 109, 633–645 (2012).
Kundrata, R., Bocakova, M., Bocak, L. The comprehensive phylogeny of the superfamily Elateroidea (Coleoptera: Elateriformia). *Mol. Phylogenet. Evol.* 76, 162–171 (2014).
Kundrata, R., Jäch, M. A., Bocak, L. Molecular phylogeny of the Byrrhoidea–Buprestoidea complex (Coleoptera, Elateriformia). *Zool. Script.* 46, 150–164 (2017).
Kurosawa, Y. Revisional notes on the family Lymexylonidae (Coleoptera) in eastern and southeastern Asia. *Bull. Natl. Sci. Mus., Tokyo, Ser. A* 11, 109–119 (1985).
Kusy, D. *et al.* Genome sequencing of *Rhinorhipus* Lawrence exposes an early branch of the

Coleotera. *Front. Zool.* 15, 21 (2018).
Kusy, D., Motyka, M., Bocek, M., Vogler, A. P. and and Bocak, L. Genome sequences identify three families of Coleoptera as morphologically derived click beetles (Elateridae). *Sci. Rep.* 8, 17084 (2018).
Lacordaire, T. *Monographie des érotylicus: famille de Pordre des coléoptères* (Roret, 1842).
Larsén, O. The skeleton and muscles of the thorax in the Coleoptera. *Opusc. Entomol*. 30, 1–241 (1966).
Lawrence, J. F. Rhinorhipidae, a new beetle family from Australia, with comments on the phylogeny of the Elateriformia. *Invert. Syst*. 2, 1–53 (1988).
Lawrence, J. F. Derodontidae (Derodontoidea), Nosodendridae (Derodontoidea) and Jacobsoniidae (Derodontoidea). in *Immature Insects, Volume 2.* (ed. Stehr, F. W.) 431–434 (Kendall/Hunt Publishing, 1991).
Lawrence, J. F. The Australian Ommatidae (Coleoptera: Archostemata): new species, larva and discussion of relationships. *Invert. Systemat.* 13, 369–390 (1999).
Lawrence, J. F. A new genus of Valdivian Scirtidae (Coleoptera) with comments on Scirtoidea and the beetle suborders. *Sukunahikona*. *Spec. Publ. Coleopterol. Soc., Osaka* 1, 351–361 (2001).
Lawrence, J. F. Classification (families and subfamilies). in *Handbook of Zoology, Arthropoda: Insecta, Coleoptera, Beetles, Vol. 1: Morphology and Systematics (Archostemata, Adephaga, Myxophaga, Polyphaga partim), 2nd ed.* (eds. Beutel, R. G., Leschen, R. A. B.) 13–22 (Walter de Gruyter, Berlin and Boston, MA, 2016).
Lawrence, J. F. Superfamily Scirtoidea Fleming, 1821. in *Australian Beetles Volume 2: Archostemata, Myxophaga, Adephaga, Polyphaga (part)* (eds. Lawrence, J. F. and Ślipiński, A.) 219–220 (CSIRO, 2019a).
Lawrence, J. F. New species of *Eucinetus* and *Noteucinetus* from Australia (Coleoptera: Scirtoidea: Eucinetidae). *Zootaxa* 4668, 151–182. (2019b).
Lawrence, J. F. A new genus and species of Cybocephalidae (Coleoptera) from Australia. *Australian Entomologist* 46, 75–84 (2019c).
Lawrence, J. F. and Leschen, R. A. B. Review of Lamingtoniidae (Coleoptera: Cucujoidea) with descriptions of two new species. in *Systematics of Coleoptera: Papers Celebrating the Retirement of Ivan Löbl* (ed. Cuccodoro, G.). *Mem. Entomol. Int.* 17, 905–919 (2003).
Lawrence, J. F., and Leschen, R. A. B. (2010). Lamingtoniidae Sen Gupta and Crowson, 1969. in *Handbook of Zoology, Arthropoda: Insecta, Coleoptera, Beetles, Vol. 2: Morphology and Systematics (Elateroidea, Bostrichiformia, Cucujiformia partim)* (eds. Leschen, R. A. B., Beutel, R. G. and Lawrence, J. F.) 360–363 (De Gruyter, Berlin and Boston, MA, 2010).
Lawrence, J. F. and Newton, A. F. Evolution and classification of beetles. *Annu. Rev. Ecol. Syst.* 13, 261–290 (1982).
Lawrence, J. F. and Britton E. B. Coleoptera. in *The Insects of Australia, Volume 2* (ed. Mackerras, I. M.) 543–683 (Melbourne University Press and Cornell University Press, Melbourne, 1991).
Lawrence, J. F. and Newton, F. Families and subfamilies of Coleoptera. in *Biology, Phylogeny, and Classification of Coleoptera: Papers Celebrating the 80th Birthday of Roy A. Crowson* (eds. Pakaluk, J. and Ślipiński, S. A.) 779–1006 (Muzeum i Instytut Zoologii PAN, 1995).
Lawrence, J. F., Ślipiński, S. A. and Pakaluk, J. From Latreille to Crowson: a history of the higher-level classification of beetles. in *Biology, Phylogeny, and Classification of Coleoptera: Papers Celebrating the 80th Birthday of Roy A. Crowson* (eds. Pakaluk, J. and Ślipiński, S. A.) 87–154 (Muzeum i Instytut Zoologii PAN, 1995a).
Lawrence, J. F., Nikitsky, N. B. and Kirejtshuk, A. G. Phylogenetic position of Decliniidae and comments on the classification of Elateriformia (*sensu lato*). in *Biology, Phylogeny, and*

*Classification of Coleoptera: Papers Celebrating the 80th Birthday of Roy A. Crowson* (eds. Pakaluk, J. and Ślipiński, S. A.) 375–410 (Muzeum i Instytut Zoologii PAN, 1995b).

Lawrence, J. F., Beutel, R. G. and Leschen, R. A. B. Derodontiformia. in *Handbook of Zoology, Arthropoda: Insecta, Coleoptera, Beetles, Vol. 2: Morphology and Systematics (Elateroidea, Bostrichiformia, Cucujiformia partim)* (eds. Leschen, R. A. B., Beutel, R. G. and Lawrence, J. F.) 179–180 (De Gruyter, Berlin and Boston, MA, 2010).

Lawrence, J. F. and Leschen, R. A. B. Jacobsoniidae Heller, 1926. in *Handbook of Zoology, Arthropoda: Insecta, Coleoptera, Beetles, Vol. 2: Morphology and Systematics (Elateroidea, Bostrichiformia, Cucujiformia partim)* (eds. Leschen, R. A. B., Beutel, R. G. and Lawrence, J. F.) 190–195 (De Gruyter, Berlin and Boston, MA, 2010).

Lawrence, J. F. and Ślipiński, S. A. *Australian Beetles, Volume 1: Morphology, Classification and Keys* (CSIRO, 2013).

Le Conte, J. L. Classification of the Coleoptera of North America. Part I. *Smithsonian Misc. Coli*. (1861).

Lee, C. F., Sato, M., Shepard, W. D. and Jaech, M. A. Phylogeny of Psephenidae (Coleoptera: Byrrhoidea) based on larval, pupal and adult characters. *Syst. Entomol.* 32, 502–538 (2007).

Legalov, A. A. Review of the family Anthribidae (Coleoptera) from the Jurassic of Karatau: subfamily Protoscelinae. Genus *Protoscelis* Medvedev. *Paleontol. J.* 47, 292–302 (2013).

Leschen, R. A. B. Phylogeny and revision of the genera of Cryptophagidae (Coleoptea: Cucujoidea). *Univ. Kansas Sci. Bull*. 55, 594–634 (1996).

Leschen, R. A. B. Erotylidae (Insecta: Coleoptera: Cucujoidea): phylogeny and review. *Fauna of New Zealand* 47, 1–108 (2003).

Leschen, R. A. B., Lawrence, J. F. and Ślipiński, S. A. Classification of basal Cucujoidea (Coleoptera: Polyphaga): cladistic analysis, keys and review of new families. *Invert. Syst.* 19, 17–73 (2005).

Leschen, R. A. B. and Buckley, T. R. Multistate characters and diet shifts: evolution of Erotylidae (Coleoptera). *Syst. Biol*. 56(1), 97–112 (2007).

Leschen, R. A. B. and Beutel R. G. Nosodendridae Erichson, 1846. in *Handbook of Zoology, Arthropoda: Insecta, Coleoptera, Beetles, Vol. 2: Morphology and Systematics (Elateroidea, Bostrichiformia, Cucujiformia partim)* (eds. Leschen, R. A. B., Beutel, R. G. and Lawrence, J. F.) 185–190 (De Gruyter, Berlin and Boston, MA, 2010).

Leschen, R. A. B., Beutel, R. G. and Lawrence, J. F. *Handbook of Zoology, Arthropoda: Insecta, Coleoptera, Beetles, Vol. 2: Morphology and Systematics (Elateroidea, BostrichiformiaCucujiformia partim)* (De Gruyter, Berlin and Boston, MA, 2010).

Leschen, R. A. B. and Slipinski, S. A. (2010). Cucujoidea, Latreille, 1802. in *Handbook of Zoology, Arthropoda: Insecta, Coleoptera, Beetles, Vol. 2: Morphology and Systematics (Elateroidea, Bostrichiformia, Cucujiformia partim)* (eds. Leschen, R. A. B., Beutel, R. G. and Lawrence, J. F.) 281–282 (De Gruyter, Berlin and Boston, MA, 2010).

Li, Y., Tihelka, E., Huang, D. and Cai, C. Specialized variegated mud-loving beetles from mid-Cretaceous Burmese amber (Coleoptera: Heteroceridae). *Palaeoentomol*. 3, 59–67 (2020a).

Li, Y., Zhao, Y., Huang, D. and Cai, C. A new throscid from mid-Cretaceous Burmese amber (Coleoptera: Elateroidea: Throscidae). *Histor. Biol.,* in press, 1–5 (2020b). https://doi.org/10.1080/08912963.2020.1741573

Liu, Z., Ślipiński, A., Leschen, R. A. B., Ren, D. and Pang, H. The oldest Prionoceridae (Coleoptera: Cleroidea) from the Middle Jurassic of China. *Ann. Zool*. 65, 41–52 (2015).

Liu, Z., Ślipiński, A., Lawrence, J. F., Ren, D. and Pang, H. *Palaeoboganium* gen. nov. from the Middle Jurassic of China (Coleoptera: Cucujoidea: Boganiidae): the first cycad pollinators? *J. Syst. Palaeontol.* 16, 351–360 (2018).

Liu, Z., Ślipiński, A., Wang, B. and Pang, H. The oldest Silvanid beetles from the Upper Cretaceous Burmese amber (Coleoptera, Silvanidae, Brontinae). *Cretaceous Res.* 98, 1–8 (2019).

Löbl, I. and Burckhardt, D. Revision der Gattung *Sarothrias* mit Bemerkungen zur Familie Jacobsoniidae (Coleoptera). *Stutt. Beit. Naturk., Ser. A* 422, 1–23 (1988).

Lü, L., Cai, C., Zhang, X., Newton, A. F., Thayer, M. K. and Zhou, H. Linking evolutionary mode to palaeoclimate change reveals rapid radiations of staphylinoid beetles in low-energy conditions. *Curr. Zool.* 66, 435–444 (2020).

Lubkin, S. *Taxonomic Studies on Paleozoic and Mesozoic Beetles, with Phylogenetic Interpretation Based on Elytral Venation and Structure*. PhD thesis, Cornell University, 154 pp (2007).

Maksoud, S., Azar, D., Granier, B. and Gèze, R. New data on the age of the Lower Cretaceous amber outcrops of Lebanon. *Palaeoworld* 26, 331–338 (2017).

Mao, Y. Y. *et al*. Various amberground marine animals on Burmese amber with discussions on its age. *Palaeoentomol*. 1, 91–93 (2018).

Martill, D. M., Bechly, G. and Loveridge, R. F. *The Crato Fossil Beds of Brazil: Window into an Ancient World* (Cambridge University Press, 2007).

Marvaldi, A. E., Duckett, C. N., Kjer, K. M. and Gillespie, J. J. Structural alignment of 18S and 28S rDNA sequences provides insights into phylogeny of Phytophaga (Coleoptera: Curculionoidea and Chrysomeloidea). *Zool. Script.* 38, 63–77 (2009).

McDermott, F. A. The taxonomy of the Lampyridae (Coleoptera). *Trans. Am. Entomol. Soc.* 90, 1–72 (1964).

McHugh, J. V. A revision of *Eurysphindus* LeConte (Coleoptera: Cucujoidea: Sphindidae) and a review of sphindid classification and phylogeny. *Syst. Entomol.* 18, 57–92 (1993).

McHugh, J. V., Marshall, C. J. and Fawcett, F. L. A study of adult morphology in *Megalodacne heros* (Say) (Coleoptera: Erotylidae). *Trans. Am. Entomol. Soc.* 123 167-223. (1997).

McKellar, R. C., Glasier, J. R. and Engel, M. S. New ants (Hymenoptera: Formicidae: Dolichoderinae) from Canadian Late Cretaceous amber. *Bull. Geosci*. 88, 583–594 (2013).

McKenna, D. D. *et al*. Phylogeny and evolution of Staphyliniformia and Scarabaeiformia: forest litter as a stepping stone for diversification of nonphytophagous beetles. *Syst. Entomol*. 40, 35–60 (2015a).

McKenna, D. D. *et al*. The beetle tree of life reveals that Coleoptera survived end-Permian mass extinction to diversify during the Cretaceous terrestrial revolution. *Syst. Entomol*. 40, 835–880 (2015b).

McKenna, D. D. *et al.* The evolution and genomic basis of beetle diversity. *Proc. Natl. Acad. Sci*. 116, 24729–24737 (2019).

Medvedev, L. N., Leafbeetles of the Jurassic of the Karatau (Coleoptera, Chrysomelidae). in *Yurskie nasekomye Karatau* (ed. Rohdendorf, B.B) 155–165 (Nauka, Moscow, 1968).

Mogutcheva, N. K. Problems of phytostratigraphy and the correlation of the Lower Jurassic continental sediments in West Siberia and Kuznetsk and Kansk-Achinsk basins. *Strat. Geol. Corr.* 17, 283–290 (2009).

Muona, J. Throscidae (Coleoptera) relationships, with descriptions of new fossil genera and species. *Zootaxa*, 4576, 521–543 (2019).

Nadein, K. S. and Perkovsky, E. E. A new tribe of Galerucinae leaf beetle (Insecta: Coleoptera: Chrysomelidae) from the Upper Cretaceous Taimyr amber. *Cretaceous Res*. 84, 97–106 (2018).

Najarro, M. *et al.* Unusual concentration of Early Albian arthropod-bearing amber in the Basque-Cantabrian Basin (El Soplao, Cantabria, Northern Spain): palaeoenvironmental and palaeobiological implications. *Geol. Acta* 7, 363–387 (2009).

Nel, A. *et al*. The earliest known holometabolous insects. *Nature* 503, 257–261 (2013).

Newton, A. F. Review of Agyrtidae (Coleoptera), with a new genus and species from New Zealand. *Ann. Zool.* 47, 111–156 (1997).

Newton, A. F. Phylogenetic problems, current classification and generic catalog of world Leiodidae (including Cholevidae). in *Phylogeny and Evolution of Subterranean and Endogean Cholevidae (=Leiodidae Cholevinae)* (eds. Giachino, P. M. and Peck, S. B.) 41–178 (Atti Museo Regionale di Scienze Naturali, Torino 1998).

Newton, A. F. Potential phylogenetic significance of the number of functional abdominal spiracles in beetle pupae, with focus on Staphylinoidea (Coleoptera). *Pap. Avulsos de Zool.* 60, e202060(s.i.).36 (2020).

Nikolajev G. V. On the Mesozoic taxa of scarabaeoid beetles of the Family Hybosoridae (Coleoptera: Scarabaeoidea). *Paleontol. J.* 44, 649–653 (2010).

Nikolajev G. V. and Ren D. New genus of the subfamily Geotrupinae (Coleoptera: Scarabaeoidea: Geotrupindae) from the Jehol Biota. *Acta Geol. Sin*. 84, 673–675 (2010).

Nikolajev, G. V., Wang, B., Liu, Y. and Zhang, H. C. Stag beetles from the Mesozoic of Inner Mongolia, China (Scarabaeoidea: Lucanidae). *Acta Palaeontol. Sin*. 50, 41–47 (2011).

Ocampo, F. C. Phylogenetic analysis of the scarab family Hybosoridae and monographic revision of the New World subfamily Anaidinae (Coleoptera: Scarabaeoidea). 1. Introduction to the scarab family Hybosoridae (Coleoptera: Scarabaeoidea). *Bull. Univ. Nebr. St. Mus.* 19, 3–6 (2006).

Pan, X., Chang, H., Ren, D. and Shih, C. The first fossil buprestids from the Middle Jurassic Jiulongshan formation of China (Coleoptera: Buprestidae). *Zootaxa* 2745, 53–62 (2011).

Peñalver, E. and Delclòs, X. Spanish amber. in D Biodiversity of fossils in amber from the major world deposits (ed. Penney, D.) 236–271 (Siri Scientific Press, Manchester, 2010).

Peris, D., Maier, C. A., Sánchez-García, A. and Delclòs, X. The oldest known riffle beetle (Coleoptera: Elmidae) from Early Cretaceous Spanish amber. *Compt. Rend. Palevol*, 14(3), 181–186 (2015a).

Peris, D., Philips, T. K. and Delclòs, X. Ptinid beetles from the Cretaceous gymnosperm-dominated forests. *Cretaceous Res.* 52, 440–445 (2015b).

Peris, D., and Fanti, F. *Molliberus albae* gen. et sp. nov., the oldest Laurasian soldier beetle (Coleoptera: Cantharidae) from the Lower Cretaceous Spanish amber. *Cretaceous Res*. 91, 263–268 (2018).

Perkovsky, E. E. Significance of antennal characters, particularly the presence of Hamann's organ on the first segment of a club, for the systematics of the agyrtid-colonid group of Staphylinoidea (Coleoptera: Agyrtidae, Leiodidae, Colonidae). *Dopov. Nac. akad. nauk Ukr.* 1997 (11), 167–174 (1997).

Perkovsky, E. E. Evolutionary development of the specific antennal structure in leiodid beetles and systematic position of Jurassic *Mesecanus communis* and *Polysitum elongatum* (Coleoptera: Staphylinoidea, Leiodidae). in *Proceedings of the First Palaeoentomological Conference, Moscow 1998*, 111–116 (AMBA projekty, Bratislava, 1999).

Perkovsky, E. E., and Makarkin, V. N. First confirmation of spongillaflies (Neuroptera: Sisyridae) from the Cretaceous. *Cretaceous Res.* 56, 363–371 (2015).

Peyerimhoff, P. Les larves des Coléoptères d'aprés A. G. Böving et F. C. Craighead, et les grandes critériums de l'Ordre. *Ann. Soc. Entomol. Fr.* 102, 77–106 (1933):

Philips, T. K. and Bell K. L. *Ptinidae Latreille 1802.* in *Handbook of Zoology, Arthropoda: Insecta, Coleoptera, Beetles, Vol. 2: Morphology and Systematics (Elateroidea, Bostrichiformia, Cucujiformia partim)* (eds. Leschen, R. A. B., Beutel, R. G. and Lawrence, J. F.) 217–225 (De Gruyter, Berlin and Boston, MA, 2010b).

Poinar Jr, G. Cretaceous palm bruchid, *Mesopachymerus antiqua*, n. gen., n. sp. (Coleoptera: Bruchidae: Pachymerini) and biogeographical implications. *Proc. Entomol. Soc. Wash.* 107, 392–

397 (2005).
Ponomarenko, A. G. Ranneyurskie zhuki-plavuntsy s r. Angary. *Paleontol. Zhur*. 1963, 128–131 (1963).
Ponomarenko A. G. Zhuki semeystva Cupedidae iz nizhnego Triasa sredney Azii. *Paleontol. Zhur.* 1966, 47–68 (1966).
Ponomarenko A. G. Composition and ecological characteristics of Mesozoic coleopterans. in *Mesozoic Coleoptera. Proceedings of the Paleontological Institute of the Academy of Sciences of the USSR, Vol. 161* (eds. Arnoldy, L. V., Zherikhin, V. V., Nikritin, L. M. and Ponomarenko A. G) 1–117 (Nauka, Moscow, 1977).
Ponomarenko, A. G. Zhestkokrylye iz Yury Sibiri i zapadnoy Mongolii. in *Yurskie nasekomye Sibiri i Mongolii. Trudy Paleontologicheskogo Instituta Akademiya Nauk SSSR 211* (ed. Rasnitsyn, A. P.) 47–87 (Nauka, Moscow, 1985).
Ponomarenko A. G. Beetles. Scarabaeida (=Coleoptera). in *Nasekomye v rannemelovykh ekosistemakh zapadnoy Mongolii. Trudy Sovmestnoj Sovetsko-Mongol'skoj Paleontologicheskoj Ekspeditsii* (ed. Rasnitsyn, A. P.) 84–105 (Nauka, Moscow, 1986).
Ponomarenko, A. G. New Mesozoic water beetles (Insecta, Coleoptera) from Asia. *Paleontol. J.* 21, 79–92 (1987).
Ponomarenko, A. G. Two new species of Mesozoic dytiscoid beetles from Asia. *Paleontol. J.* 27, 9–34 (1993).
Ponomarenko, A. G. *et al.* Upper Jurassic Lagerstätte Shar Teg, southwestern Mongolia. *Paleontol. J*. 48, 1573–1682 (2014).
Pons, D., Berthou, P.-Y. and Campos, D. de A. Quelques observations sur la palynologie de l’Aptien sup´erieur et de l’Albien du Bassin d’Araripe. in *Atas do I Simposio sobre a Bacia do Araripe e bacias interiores do nordeste, Crato, 14 a 16 de junho de 1990* (eds. Campos, D. de A., Viana, M. S. S., Brito, P. M. and Beurlen, G.) 241–252 (Crato, 1990).
Prado, A. The cycad herbivores. *Bull. Soc. Entomol. Quebec* 18, 3–6 (2011).
Prokin A. A. and Ren, D. New species of variegated mud-loving beetles (Coleoptera: Heteroceridae) from Mesozoic deposits of China. *Paleontol. J.* 45, 284–286 (2011).
Prokin, A. A., Petrov, P. N., Wang, B. and Ponomarenko, A. G. New fossil taxa and notes on the Mesozoic evolution of Liadytidae and Dytiscidae (Coleoptera). *Zootaxa* 3666, 137–159 (2013).
Rasnitsyn, A. P. and Zherikhin, V. V. First fossil chewing louse from the lower Cretaceous of Baissa, Transbaikalia (Insecta, Pediculida= Phthiriaptera, Saurodectidae fam. n.). *Rus. Entomol. J.* 8, 253–255 (1999).
Robertson, J. A., McHugh, J. V. and Whiting, M. F. A molecular phylogenetic analysis of the pleasing fungus beetles (Coleoptera: Erotylidae): evolution of colour patterns, gregariousness and mycophagy. *Syst. Entomol.* 29, 173–187 (2004).
Robertson J.A. *et al.* Phylogeny and classification of Cucujoidea and the recognition of a new superfamily Coccinelloidea (Coleoptera: Cucujiformia) *Syst. Entomol.* 40, 745–778 (2015).
Rohdendorf, B. B. *Jurassic Insects of Karatau* (Nauka, 1968).
Sadovnikov, G. N. Evolution of the biome of the middle Siberian Trappean plateau. *Palaeontol. J.* 50, 518–532 (2016).
Saks *et al.* The Mesozoic deposits of the Khatanga depression. *Trudy Nauch.-issledovatel. Instit. Geol. Arkt.* 99, 1–246 (1959).
Schneider, J. W. and Werneburg, R. Insect biostratigraphy of the European Late Carboniferous and Early Permian. in *Non-marine Permian Biostratigraphy and Biochronology* (eds. Lucas, S. G., Cassinis G. and Schneider J. W.) 325–336 (Geological Society of London, Special Publication, 2006).

Schneider, J. W. and Werneburg, R. Biostratigraphie des Rotliegend mit insekten und amphibien. *Schrift*. *Dtsch. Gesellsch. Geowiss*. 61, 110–142 (2012).
Schütz, C. and Dettner, K. Cantharidin-secretion by elytral notches of male Anthicid species (Coleoptera: Anthicidae). *Z. Naturforsch. C* 47, 290–299 (1992).
Selander, R. B. A classification of the genera and higher taxa of the meloid subfamily Eleticinae (Coleoptera). *Can. Entomol.* 98, 449–481 (1966).
Selander, R. B. Meloidae (Tenebrionoidea). in *Immature Insects, Volume 2.* (ed. Stehr, F. W.) 530–534 (Kendall/Hunt Publishing, 1991).
Sen Gupta, T. A new subfamily of Merophysiidae (Clavicornia: Coleoptera) and descriptions of two new species of Gomya Dajoz and its larva. *Rev. Suisse Zool.* 86, 691–698 (1979).
Sen Gupta, T. Review of the genera of the family Rhizophagidae (Clavicornia: Coleoptera) of the world. *Mem. Zool. Surv. Ind.* 17, 1–58 (1988).
Sen Gupta, T. S. and Crowson, R. A. Further observations on the family Boganiidae, with definition of two new families Cavognathidae and Phloeostichidae. *J. Nat. Hist.* 3, 571–590 (1969).
Sen Gupta, T. and Crowson, R. A. The coleopteran family Sphindidae. *Entomol. Month. Mag*. 113, 177–191. (1979).
Shcherbakov, D. E. Madygen, Triassic Lagerstätte number one, before and after Sharov. *Alavesia* 2, 113–124 (2008).
Shi, G. *et al.* Age constraint on Burmese amber based on U–Pb dating of zircons. *Cretaceous Res*. 37, 155–163 (2012).
Shockley, F. W. and Alekseev, V. I. *Glesirhanis bercioi*, a new genus and species from Baltic amber (Coleoptera: Endomychidae: Leiestinae) with a checklist and nomenclatural notes regarding fossil Endomychidae. *Zootaxa* 3755, 391–400 (2014).
Sikes, D. S. Silphidae Latreille, 1807. in Handbook of Zoology, Arthropoda: Insecta, Coleoptera, Beetles, Vol. 1: Morphology and Systematics (Archostemata, Adephaga, Myxophaga, Polyphaga partim), 2nd ed. (eds. Beutel, R. G., Leschen, R. A. B.) 386–394 (Walter de Gruyter, Berlin and Boston, MA, 2016).
Ślipiński, S. A. Revision and phylogeny of Protocucujidae [Coleoptera: Cucujoidea]. *Ann. Zool.* 48, 295–316 (1998).
Song, H., Sheffield, N. C., Cameron, S. L., Miller, K. B. and Whiting, M. F. When phylogenetic assumptions are violated: base compositional heterogeneity and among-site rate variation in beetle mitochondrial phylogenomics. *Syst. Entomol*. 35 429–448 (2010).
Song, N., Zhai, Q. and Zhang, Y. Higher-level phylogenetic relationships of rove beetles (Coleoptera, Staphylinidae) inferred from mitochondrial genome sequences. *Mitochondrial DNA Part A*, in press (2021).
Suinyuy, T. N., Donaldson, J. S. and Johnson, S. D. Insect pollination in the African cycad *Encephalartos friderici-guilielmi* Lehm. *S. Afr. J. Bot.* 75, 682–688 (2009).
Swisher, C. C., Wang, Y. Q., Wang, X. L., Xu, X. and Wang, Y. Cretaceous age for the feathered dinosaurs of Liaoning, China. *Nature* 400, 58–61 (1999).
Thayer, M. K. Staphylinidae. in *Handbook of Zoology, Arthropoda: Insecta, Coleoptera, Beetles, Vol. 1: Morphology and Systematics (Archostemata, Adephaga, Myxophaga, Polyphaga partim), 2nd ed.* (eds. Beutel, R. G., Leschen, R. A. B.) 394–442 (Walter de Gruyter, Berlin and Boston, MA, 2016).
Thomas, M. C. A new species of apterous *Telephanus* (Coleoptera: Silvanidae) with a discussion of phylogenetic relationships of the Silvanidae. *Coleopt. Bull*. 43–55 (1984a).
Thomas, M. C. A new Neotropical genus and species of rostrate Laemophloeinae (Coleoptera: Cucujidae) with discussion of the systematic position of the subfamily. *Coleopt. Bull.* 67–83

(1984b).
Thomas, M. C. The Brontini of the world: A generic review of the tribe (Coleoptera: Silvanidae: Brontinae). *Insecta Mundi* 17, 1–31 (2003).
Tihelka, E., Huang, D. and Cai, C. A new genus and tribe of Cretaceous net-winged beetles from Burmese amber (Coleoptera: Elateroidea: Lycidae). *Palaeoentomol.* 2, 262–270 (2019).
Timmermans, M. J. and Vogler, A. P. Phylogenetically informative rearrangements in mitochondrial genomes of Coleoptera, and monophyly of aquatic elateriform beetles (Dryopoidea). *Mol. Phylogenet. Evol.* 63, 299–304 (2012).
Timmermans, M. J., Barton, C., Haran, J., Ahrens, D., Culverwell, C. L., Ollikainen, A., Dodsworth, S., Foster, P. G., Bocak, L., Vogler, A. P. Family-level sampling of mitochondrial genomes in Coleoptera: compositional heterogeneity and phylogenetics. *Genome Biol. Evol.* 8, 161–175. (2016).
Tomaszewska, W. Morphology, phylogeny and classification of adult Endomychidae (Coleoptera: Cucujoidea) *Ann. Zool.* 50, 449–558 (2000).
Tomaszewska, W., Ślipiński, A., Bai, M., Zhang, W. and Ren, D. The oldest representatives of Endomychidae (Coleoptera: Coccinelloidea) from the Upper Cretaceous Burmese amber. *Cretaceous Res.* 91, 287–298 (2018).
Toussaint, E. F. *et al*. The peril of dating beetles. *Syst. Entomol.* 42, 1–10 (2017).
Turnbow, R. H. Jr and Thomas, M. C. Cerambycidae Leach 1815. in *American Beetles. Vol. 2. Polyphaga: Scarabaeoidea through Curculionoidea* (eds. Arnett, R. H., Thomas, M. C., Skelley, P. E. and Frank J. H.) 568–601 (CRC Press, Boca Raton, 2002).
Vanin, S. A., Costa C., Ide, S. and Beutel, R. G. Heteroceridae MacLeay, 1825. in *Handbook of Zoology, Arthropoda: Insecta, Coleoptera, Beetles, Vol. 1: Morphology and Systematics (Archostemata, Adephaga, Myxophaga, Polyphaga partim), 2nd ed.* (eds. Beutel, R. G., Leschen, R. A. B.) 612–615 (Walter de Gruyter, Berlin and Boston, MA, 2016).
Vitali, F. Systematic notes on the Cerambycidae (Insecta: Coleoptera) described from Burmese amber. *Palaeoentomol*. 2, 215–218 (2019).
Voigt, S., Buchwitz, M., Fischer, J., Kogan, I., Moisan, P., Schneider, J. W., Spindler, F., Brosig, A., Preusse, M., Scholze, F. and Linnemann, U. Triassic life in an inland lake basin of the warm-temperate biome – the Madygen Lagerstätte (Southwest Kyrgyzstan, Central Asia). *in*: *Terrestrial Conservation Lagerstätten: Windows into the Evolution of Life on Land* (eds. Fraser, N. C. and Sues, H.-D.) 65–104 (Dunedin Academic Press, Edinburgh, 2017).
Wang, B., *et al.* The earliest known longhorn beetle (Cerambycidae: Prioninae) and implications for the early evolution of Chrysomeloidea. *J. System. Palaeontol.* 12, 565–574 (2014).
Wang, X. *et al*. Discussion on the relationship and era of the Daohugou fossils in Ningcheng of Inner Mongolia. *Chinese Sci. Bull*. 50, 2127–2135 (2005).
Wegrzynowicz, P. Morphology, phylogeny and classification of the family Erotylidae based on adult characters (Coleoptera: Cucujoidea). *Genus* 13, 435–504 (2002).
Wolf-Schwenninger, K. The oldest fossil record of Lymexylidae (Insecta: Coleoptera) from the Lower Cretaceous Crato Formation of Brazil. *Insect Syst. Evol*. 42, 205–212 (2011).
Yamamoto, S. Tachyporinae revisited: phylogeny, evolution, and higher classification based on morphology, with recognition of a new rove beetle subfamily (Coleoptera: Staphylinidae). *Biology* 10, 323 (2021).
Yamamoto, S. and Takahashi, Y. First discovery of Coloninae fossil in Cretaceous Burmese amber (Coleoptera, Staphylinoidea, Leiodidae). *PalZ* 92, 195–201 (2018).
Yamamoto, S., Takahashi, Y. and Parker, J. Evolutionary stasis in enigmatic jacobsoniid beetles. *Gondwana Res*. 45, 275–281 (2017).

Yan, E. V. and Wang, B. A new genus of elateriform beetles (Coleoptera, Polyphaga) from the Jurassic of Daohugou, China. *Paleontol. J*. 44, 297–302 (2010).
Yan, E. V., Lawrence, J. F., Beattie, R. and Beutel, R. G. At the dawn of the great rise: †*Ponomarenkia belmonthensis* (Insecta: Coleoptera), a remarkable new Late Permian beetle from the Southern Hemisphere. *J. Syst. Palaeontol.* 16, 611–619 (2018a).
Yan, E. V., Beutel, R. G., Beattie, R. and Lawrence, J. F. *Ponomarenkium* gen. nov., a replacement name for the stem group beetle *Ponomarenkia* Yan et al. 2017 (Insecta: Coleoptera). *Paleontol. J.* 52, 220–220 (2018b).
Yan, E. V., Beutel, R. G. and Lawrence, J. F. Whirling in the late Permian: ancestral Gyrinidae show early radiation of beetles before Permian-Triassic mass extinction. *BMC Evol. Biol*. 18, 33 (2018c).
Yan, E. V., Beutel, R. G. and Ponomarenko, A. G. †Peltosynidae, a new beetle family from the Middle–Late Triassic of Kyrgyzstan: its affinities with Polyphaga (Insecta, Coleoptera) and the groundplan of this megadiverse suborder. *J. Syst. Palaeontol*. 16, 515–530 (2018d).
Yan, Z., Bai, M. and Ren, D. A new fossil Hybosoridae (Coleoptera: Scarabaeoidea) from the Yixian Formation of China. *Zootaxa* 3478, 201–204 (2012).
Yu, Y., Davis, S. R., Shih, C., Ren, D. and Pang, H. The earliest fossil record of Belidae and its implications for the early evolution of Curculionoidea (Coleoptera). *J. Syst. Palaeontol.* 17, 2105–2117 (2019).
Zhang, J. (2015). Archisargoid flies (Diptera, Brachycera, Archisargidae and Kovalevisargidae) from the Jurassic Daohugou biota of China, and the related biostratigraphical correlation and geological age. *J. Syst. Palaeontol*. 13, 857–881.
Zhang, S.-Q. et al. Evolutionary history of Coleoptera revealed by extensive sampling of genes and species. *Nat. Commun*. 9, 1–11 (2018).
Zheng, D. et al. Middle-Late Triassic insect radiation revealed by diverse fossils and isotopic ages from China. *Sci. Adv.* 4, eaat1380 (2018).
Zherikhin, V. V. Development and changes of the Cretaceous and Cenozoic faunal assemblages (Tracheata and Chelicerata). *Trudy Paleontol. Instit. Akad. Nauk SSSR* 165, 1–198 (1978).
Zherikhin, V. V. and Gratshev, V. G. Fossil weevil beetles (Coleoptera, Curculionoidea) from the Lower Cretaceous of northeastern Brazil. *Paleontol. J.* 38, 528–537 (2004).
Zwick, P. and Zwick, H. Beitrag zur morphologie, systematik und biologie mitteleuropäischer Scirtidae (Coleoptera). *Entomol. Nachr. Berich.* 52, 185–191 (2008).
